## Supplementary material for "LiveCellMiner: A New Tool to Analyze Mitotic Progression": Note S1

### S1 Note: Experimental Details

#### LSM5live Experiments. LSD1 Data Set [1]

HeLa cells expressing H2B-mCherry and $\alpha$-tubulin-EGFP were transfected with the indicated siRNA oligonucleotides and seeded in eight-well µ-slide chambers (Ibidi). Starting at 30 h post-transfection, cells were imaged using a Plan-Apochromat 10× NA 0.45 objective and a 561-nm diode laser on a LSM5 live confocal microscope (Zeiss) equipped with a heating and CO2 incubation system (Ibidi). ZEN software (Zeiss) was used to acquire images from five 7.5-µm-spaced optical z-sections at various xy positions every 3 min. Single position *.ome files were generated from the maximum intensity projections in ZEN and converted into image sequences with Fiji software.

#### LSM5live Experiments. RecQL4 Data Set [2]

HeLa cells expressing H2B-mCherry and EGFP-$\alpha$-tubulin were transfected with the indicated siRNA oligonucleotides in eight-well µ-slide chambers (Ibidi) and, after 24 h, were imaged for 48 h in a LSM 5 live confocal microscope (Zeiss) equipped with a heating and CO2 incubation system (Ibidi). Seven 3.6-µm-spaced optical z-sections at various positions every 3 min were acquired with a Plan-Apochromat 20× NA 0.8 objective and a 488-nm and 561-nm diode lasers controlled by ZEN software. For the analysis, maximum intensity projections in Z were generated in ZEN for every position and converted into temporal image sequences with the free licensed AxioVision software (LE64; V4.9.1.0).

#### LSM710 Experiments. CTRL vs PP2A Data Set (Unpublished)

HeLa cells expressing H2B-mCherry were transfected with the indicated siRNA oligonucleotides in eight-well µ-slide chambers (Ibidi) and, after 24 h, were imaged for 48 h in a LSM 710 confocal microscope (Zeiss) equipped with an Incubator XL S1(Zeiss). Seven 2.6-µm-spaced optical z-sections at various positions every 3 min were acquired with a Plan-Apochromat 20× NA 0.8 objective and a 488-nm Argon and 561-nm DPSS lasers controlled by ZEN software (Zeiss). For the analysis, maximum intensity projections in Z were generated for every position and converted into temporal image sequences in ZEN 2.3 software (Zeiss).

### Nikon Experiments. VPS72, INO80, SRCAP, EP400 and H2A.Z. Data Set [3]

HeLa cells expressing H2B-mCherry were transfected with the indicated siRNA oligonucleotides in eight-well $\mu$-slide chambers (Ibidi) and, after 48 h, were imaged for 48 h with the widefield module of a Ti2 Eclipse (Nikon) equipped with a LED light engine SpectraX (Lumecor) and GFP/mCherry filter sets, a Plan-Apochromat 10x NA 0.5 objective and environmental control system (Ibidi). Elements software (Nikon) was used to perform fluorescence multi-position imaging every three minutes and the subsequent conversion to image sequences.

### Nikon Experiments. RPE cells. Data Set [3]

RPE cells expressing H2B-mCherry were transfected with 20nM of the indicated siRNA oligonucleotides in eight-well µ-slide chambers (Ibidi) and, after 48 h, were imaged for 48 h with the widefield module of a Ti2 Eclipse (Nikon) equipped with a LED light engine SpectraX (Lumecor) and GFP/mCherry filter sets, a Plan-Apochromat 20x NA 0.75 air objective and environmental control system (Ibidi). Elements software (Nikon) was used to perform fluorescence multi-position imaging every three minutes and the subsequent conversion to image sequences.

### Screening Data Set by Hériché *et al.* [4]

The screening data set is publicly available at Image Data Resource (IDR) (`https://idr.openmicroscopy.org/webclient/?show=screen-102`). HeLa cells stably expressing HIST1H2BJ-mCherry and LMNA-eGFP in each well of siRNA-coated 96-well plates. The images were acquired with an Olimpus IX-81 automated epifluorescence microscope with a 20× objective and a time interval of 8.5 min for 44 h. Four independent replicates were acquired for each siRNA treatment.
