## Supplementary material for "LiveCellMiner: A New Tool to Analyze Mitotic Progression": File S1: LSD1_FusedProjects_CARSync_AdditionalFeatures_FeatureReport.htm

|  |  |  |  |  |  |  |  |  |  |  |  |  |  |  |  |
| --- | --- | --- | --- | --- | --- | --- | --- | --- | --- | --- | --- | --- | --- | --- | --- |
| **Time Series Name** | **Min** | **Max** | **Mean** | **Std** | **Median** | **n-Fold Inc. I->P (All)** | **n-Fold Inc. I->P (scrambled)** | **n-Fold Inc. I->P (LSD1-2)** | **n-Fold Inc. I->P (LSD1-6)** | **n-Fold Inc. I->P (PP2A)** | **n-Fold Inc. I->A (All)** | **n-Fold Inc. I->A (scrambled)** | **n-Fold Inc. I->A (LSD1-2)** | **n-Fold Inc. I->A (LSD1-6)** | **n-Fold Inc. I->A (PP2A)** |
| Orientation | -73.56 | 73.97 | 0.05 | 39.12 | 0.17 | -153.08 | -26.31 | 104.15 | 792.99 | -1546.79 | -777.76 | -41.32 | 10.02 | -2831.00 | 91.50 |
| StdIntensity | 5.58 | 49.21 | 12.77 | 11.34 | 7.70 | 179.04 | 190.75 | 167.99 | 175.09 | 180.04 | 497.80 | 537.95 | 455.15 | 504.22 | 482.92 |
| StdIntensity-Normalized | 0.15 | 1.32 | 0.34 | 0.30 | 0.20 | 179.04 | 190.75 | 167.99 | 175.09 | 180.04 | 497.80 | 537.95 | 455.15 | 504.22 | 482.92 |
| StdIntensityGradMag | 11.99 | 81.39 | 22.58 | 16.68 | 15.22 | 118.42 | 126.52 | 130.03 | 115.05 | 104.25 | 370.33 | 399.76 | 364.26 | 367.18 | 347.76 |
| Ch2-StdIntensity-Ext-disk-r=3 | 4.53 | 40.09 | 10.73 | 8.99 | 6.88 | 273.40 | 260.39 | 249.83 | 315.67 | 259.20 | 221.00 | 211.99 | 216.58 | 238.77 | 214.30 |
| AngularSecondMoment | 0.00 | 0.05 | 0.01 | 0.01 | 0.01 | 50.48 | 52.76 | 50.07 | 67.51 | 29.48 | 217.49 | 208.41 | 227.67 | 212.96 | 223.86 |
| manualSynchronization | 1.00 | 3.00 | 2.46 | 0.82 | 3.00 | 100.00 | 100.00 | 100.00 | 100.00 | 100.00 | 200.00 | 200.00 | 200.00 | 200.00 | 200.00 |
| DifferenceVariance | 13.08 | 56.25 | 27.51 | 8.12 | 26.75 | 26.26 | 23.56 | 50.19 | 14.97 | 22.28 | 142.84 | 138.30 | 179.66 | 123.80 | 138.96 |
| Contrast | 13.16 | 56.42 | 27.63 | 8.14 | 26.87 | 26.12 | 23.44 | 49.94 | 14.88 | 22.17 | 142.19 | 137.68 | 178.73 | 123.29 | 138.37 |
| InformationMeasureofCorrelationI | -0.47 | -0.19 | -0.27 | 0.07 | -0.24 | -42.57 | -44.10 | -27.76 | -49.00 | -45.81 | -108.38 | -111.27 | -87.24 | -121.37 | -108.02 |
| MeanIntensity-NormalizedIntMean | 0.86 | 2.43 | 1.15 | 0.41 | 0.98 | 40.95 | 43.71 | 42.91 | 42.77 | 34.43 | 99.15 | 104.48 | 95.40 | 96.75 | 99.26 |
| MeanIntensity | 53.32 | 150.12 | 71.80 | 25.32 | 61.11 | 40.95 | 43.71 | 42.91 | 42.77 | 34.43 | 99.15 | 104.48 | 95.40 | 96.75 | 99.26 |
| MeanIntensity-Normalized | 0.46 | 1.29 | 0.62 | 0.22 | 0.53 | 40.95 | 43.71 | 42.91 | 42.77 | 34.43 | 99.15 | 104.48 | 95.40 | 96.75 | 99.26 |
| Ch2-MeanIntensity-Ext-disk-r=3 | 42.08 | 114.16 | 56.16 | 20.12 | 47.28 | 67.45 | 64.71 | 75.84 | 74.56 | 55.64 | 94.24 | 89.07 | 98.31 | 99.12 | 90.96 |
| Ch2-MaxIntensity-Ext-disk-r=3 | 59.08 | 202.96 | 89.86 | 37.58 | 73.39 | 109.70 | 108.53 | 110.48 | 119.11 | 99.84 | 83.63 | 81.78 | 85.10 | 90.00 | 77.33 |
| Area | 53.69 | 308.80 | 164.02 | 65.65 | 152.24 | -29.88 | -31.71 | -29.91 | -33.26 | -24.16 | -72.44 | -73.20 | -70.26 | -73.06 | -72.70 |
| MinorAxisLength | 6.14 | 16.83 | 11.44 | 2.66 | 11.23 | -15.27 | -16.31 | -14.29 | -17.11 | -12.92 | -50.88 | -51.18 | -47.97 | -51.25 | -52.54 |
| MajorAxisLength | 10.57 | 26.31 | 18.07 | 3.77 | 17.69 | -15.42 | -16.90 | -15.90 | -17.45 | -11.23 | -45.74 | -47.07 | -44.20 | -46.09 | -45.21 |
| Ratio-MajorAxisLength-vs-MinorAxisLength | 1.12 | 2.41 | 1.62 | 0.28 | 1.60 | 0.89 | 0.22 | -0.94 | 0.87 | 3.10 | 15.60 | 13.25 | 11.42 | 16.01 | 21.00 |
| DifferenceEntropy | 1.90 | 2.58 | 2.29 | 0.14 | 2.31 | 5.91 | 5.58 | 8.00 | 5.15 | 5.40 | 13.27 | 12.88 | 15.01 | 12.63 | 12.97 |
| Ch1-Ch2-MI-Ratio-Ext-disk-r=3 | 0.91 | 1.41 | 1.18 | 0.11 | 1.18 | -18.68 | -15.94 | -21.23 | -21.13 | -16.79 | -10.14 | -5.15 | -12.87 | -13.51 | -9.43 |
| MaximalCorrelationCoefficient | 0.73 | 0.94 | 0.84 | 0.05 | 0.83 | 10.24 | 10.52 | 5.85 | 11.80 | 11.75 | 10.06 | 10.56 | 6.72 | 12.21 | 9.86 |
| Correlation | 0.71 | 0.94 | 0.82 | 0.05 | 0.81 | 10.78 | 11.06 | 6.26 | 12.27 | 12.48 | 10.01 | 10.58 | 6.69 | 12.18 | 9.70 |
| InformationMeasureofCorrelationII | 0.85 | 1.00 | 0.93 | 0.04 | 0.92 | 7.16 | 7.43 | 4.47 | 8.49 | 7.59 | 9.68 | 9.98 | 6.80 | 11.43 | 9.76 |
| SumAverage | 59.01 | 104.78 | 88.99 | 10.58 | 90.92 | -3.56 | -3.75 | -3.91 | -1.29 | -5.59 | -9.15 | -10.61 | -10.60 | -8.73 | -6.88 |
| MinMajAxisRatio | 0.42 | 0.90 | 0.64 | 0.11 | 0.64 | 1.93 | 2.60 | 3.79 | 2.59 | -1.00 | -8.24 | -6.47 | -4.82 | -8.29 | -12.82 |
| Ratio-MinorAxisLength-vs-MajorAxisLength | 0.42 | 0.90 | 0.64 | 0.11 | 0.64 | 1.93 | 2.60 | 3.79 | 2.59 | -1.00 | -8.24 | -6.47 | -4.82 | -8.29 | -12.82 |
| Circularity | 0.63 | 0.98 | 0.84 | 0.07 | 0.85 | -4.66 | -3.03 | -6.85 | -3.65 | -5.72 | 5.68 | 7.09 | 3.75 | 5.94 | 5.46 |
| SumEntropy | 3.42 | 4.32 | 3.92 | 0.18 | 3.94 | 6.74 | 6.62 | 5.48 | 6.33 | 8.34 | 4.24 | 4.21 | 3.66 | 4.63 | 4.30 |
| Variance | 1088.18 | 2829.47 | 2139.47 | 410.37 | 2194.63 | -0.68 | -2.52 | 0.51 | 2.83 | -3.61 | -3.93 | -7.39 | -4.38 | -3.35 | -0.56 |
| SumVariance | 3883.59 | 10525.30 | 7845.83 | 1561.71 | 8045.04 | -0.66 | -2.79 | 0.95 | 2.99 | -3.79 | -3.54 | -7.30 | -3.58 | -2.98 | -0.16 |
| Entropy | 6.75 | 8.74 | 7.81 | 0.39 | 7.82 | 7.37 | 7.85 | 5.13 | 7.30 | 8.76 | -1.54 | -1.03 | -3.54 | -0.75 | -1.32 |
| InverseDifferenceMoment(Homogeneity) | 0.21 | 0.40 | 0.28 | 0.04 | 0.27 | 3.09 | 3.23 | -2.63 | 7.83 | 2.29 | -1.00 | -0.31 | -6.33 | 3.52 | -2.45 |
| **Single Feature Name** | **Min** | **Max** | **Mean** | **Std** | **Median** |  |  |  |  |  |  |  |  |  |  |
| IPToMALength\_Frames | 4.00 | 29.00 | 10.80 | 3.43 | 10.00 | - | - | - | - | - | - | - | - | - | - |
| IPToMALength\_Minutes | 12.00 | 87.00 | 32.39 | 10.29 | 30.00 | - | - | - | - | - | - | - | - | - | - |
| InterphaseMeanIntensity | 35.41 | 155.28 | 62.37 | 14.76 | 59.58 | - | - | - | - | - | - | - | - | - | - |
| AccumulatedOrientationDiffPMA | 18.78 | 1133.52 | 236.77 | 165.89 | 188.19 | - | - | - | - | - | - | - | - | - | - |
| Ch2-MeanIntensity-Ext-disk-r=3-intMean | 29.56 | 101.69 | 46.72 | 6.95 | 45.56 | - | - | - | - | - | - | - | - | - | - |
| Ch2-MeanIntensity-Ext-disk-r=3-pmaMean | 30.23 | 218.06 | 101.76 | 23.94 | 99.20 | - | - | - | - | - | - | - | - | - | - |
| Ch2-MeanIntensity-Ext-disk-r=3-atiMean | 29.73 | 163.88 | 50.74 | 7.71 | 49.73 | - | - | - | - | - | - | - | - | - | - |
