## Supplementary figures and images for "LiveCellMiner: A New Tool to Analyze Mitotic Progression"

### LSD1_FusedProjects_CARSync_AdditionalFeatures_AccumulatedOrientationDiffPMA_BoxPlots.png

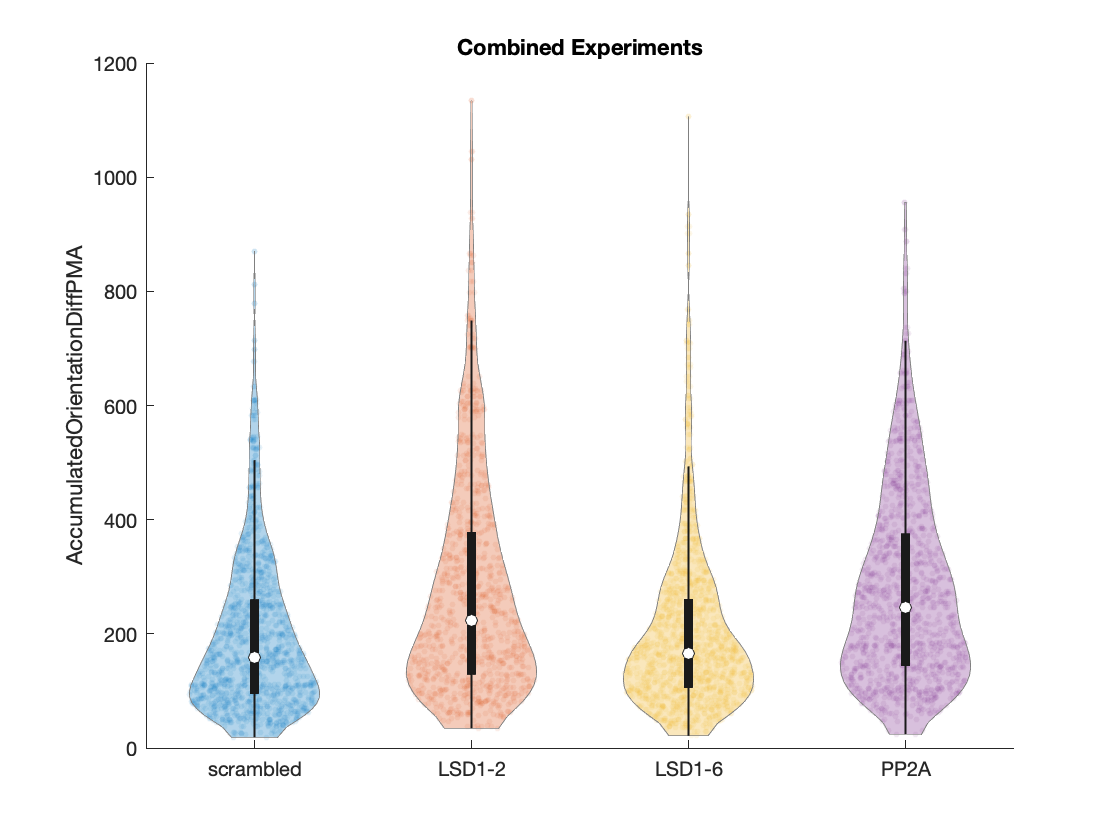

### LSD1_FusedProjects_CARSync_AdditionalFeatures_AccumulatedOrientationDiffPMA_Histograms.png

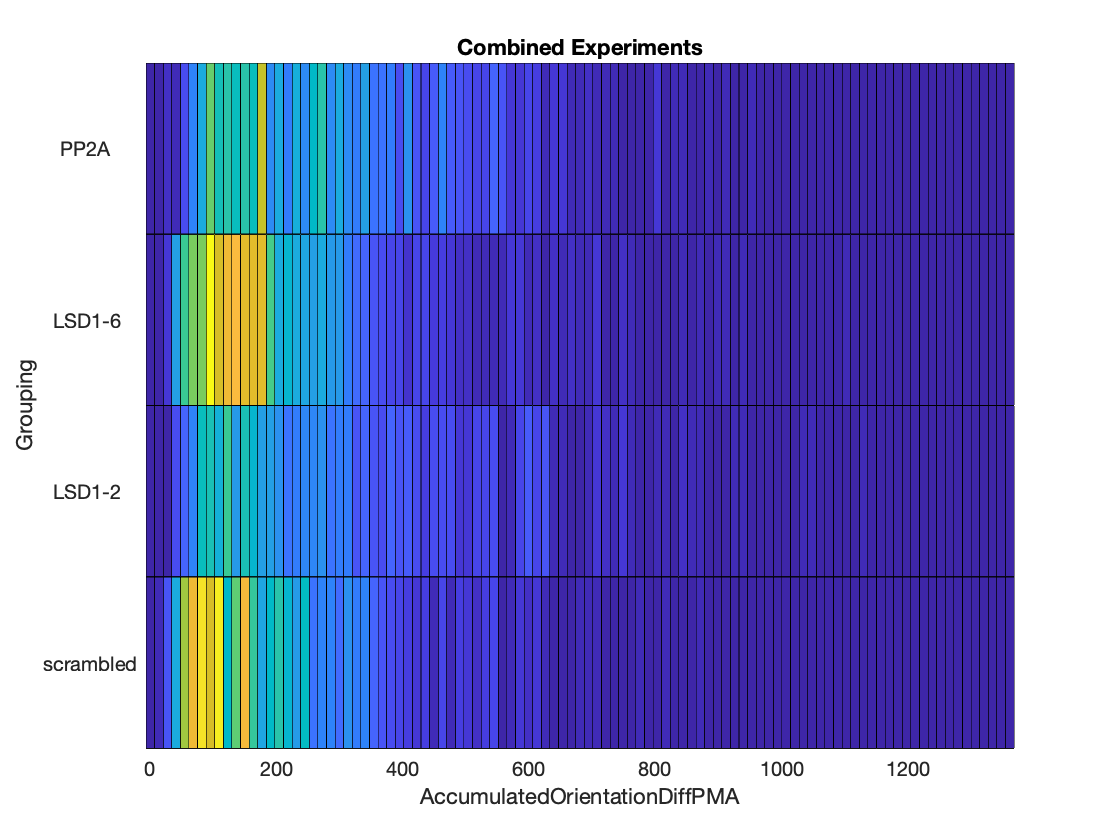

### LSD1_FusedProjects_CARSync_AdditionalFeatures_AngularSecondMoment_HeatMaps.png

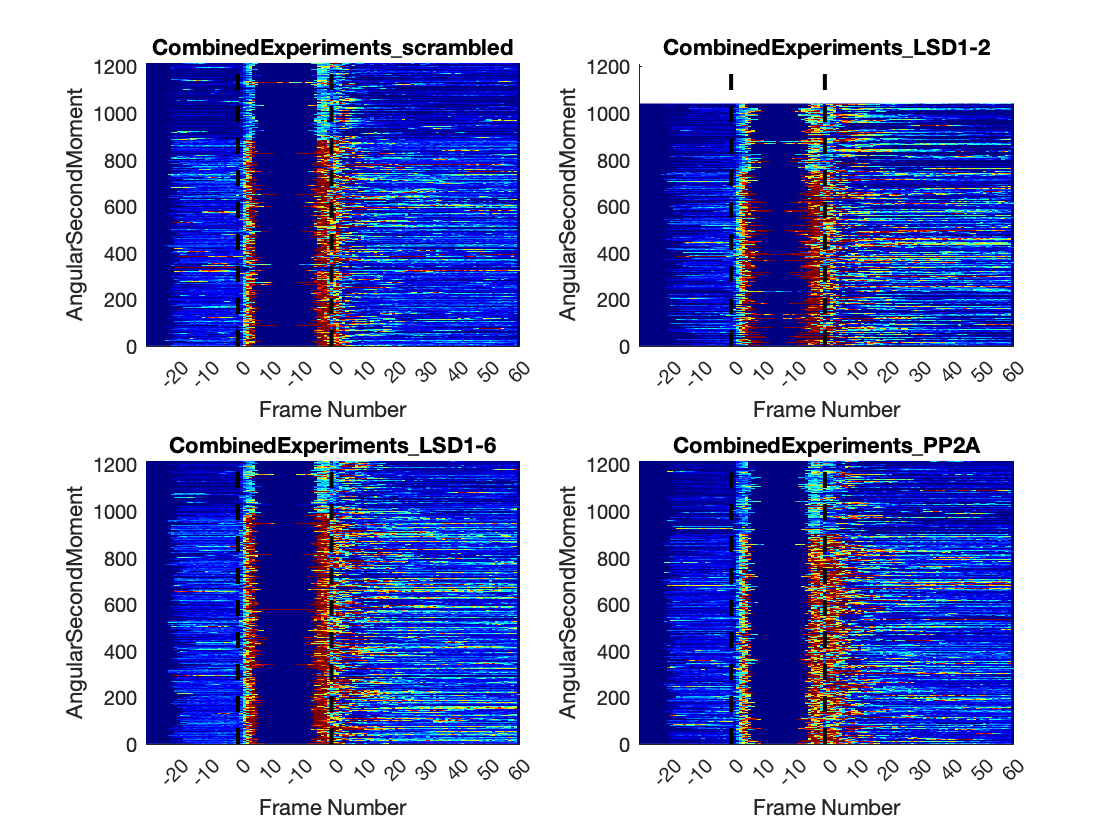

### LSD1_FusedProjects_CARSync_AdditionalFeatures_AngularSecondMoment_LinePlots.png

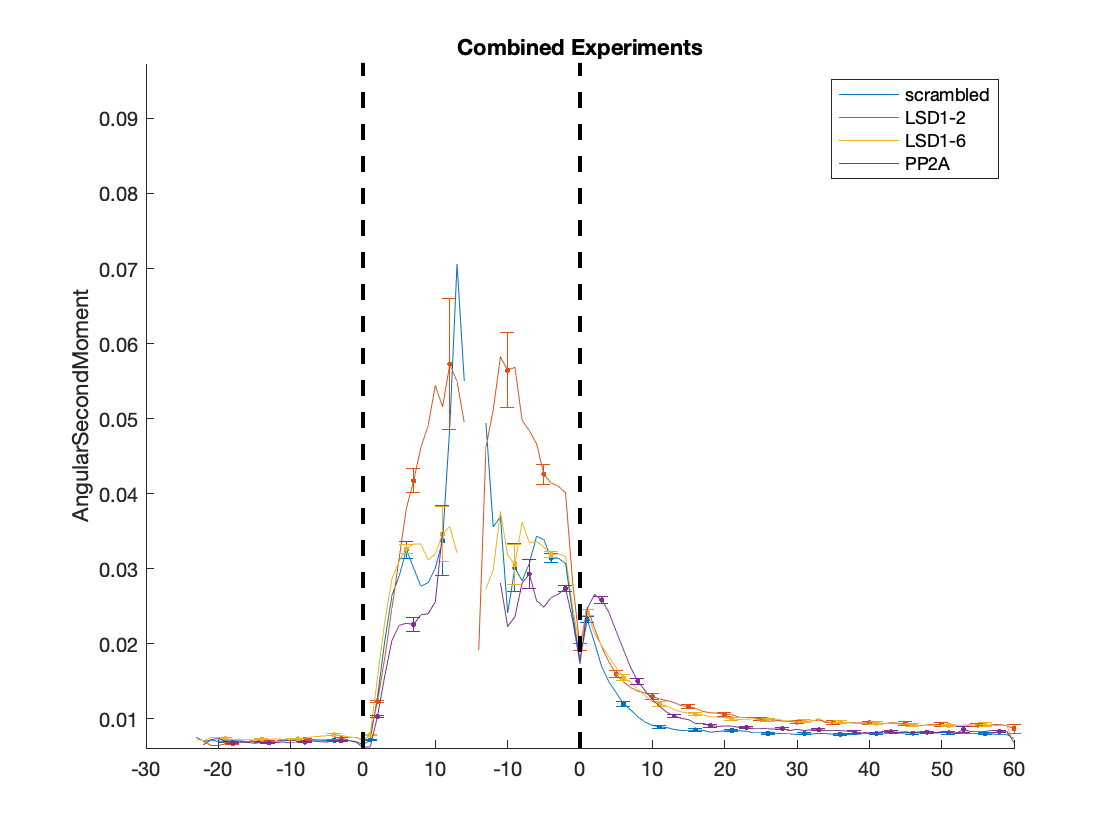

### LSD1_FusedProjects_CARSync_AdditionalFeatures_Area_HeatMaps.png

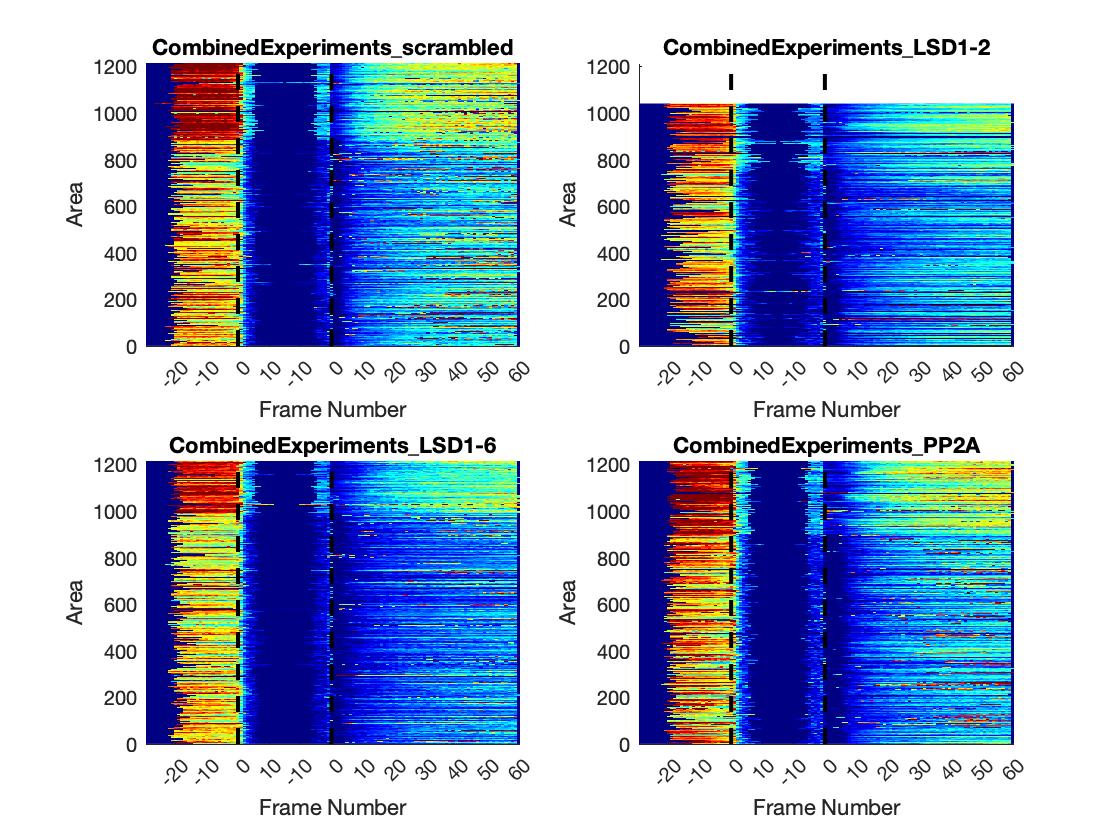

### LSD1_FusedProjects_CARSync_AdditionalFeatures_Area_LinePlots.png

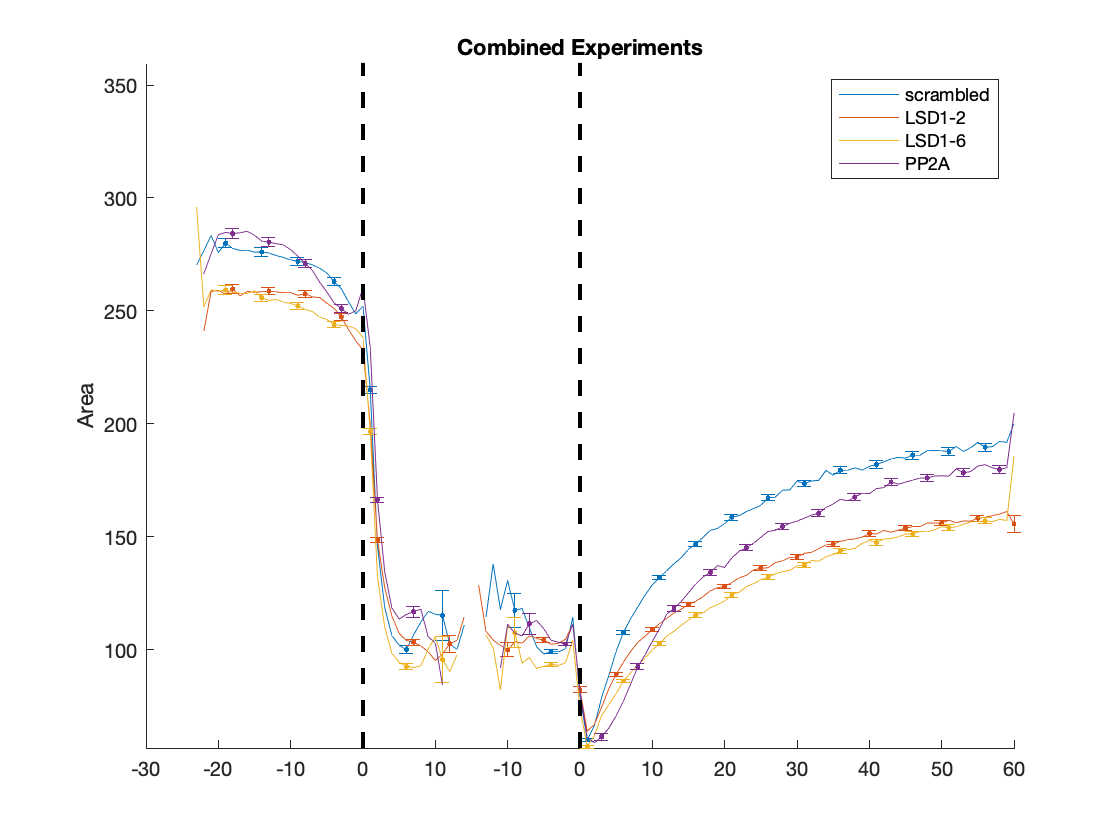

### LSD1_FusedProjects_CARSync_AdditionalFeatures_Ch1-Ch2-MI-Ratio-Ext-disk-r=3_HeatMaps.png

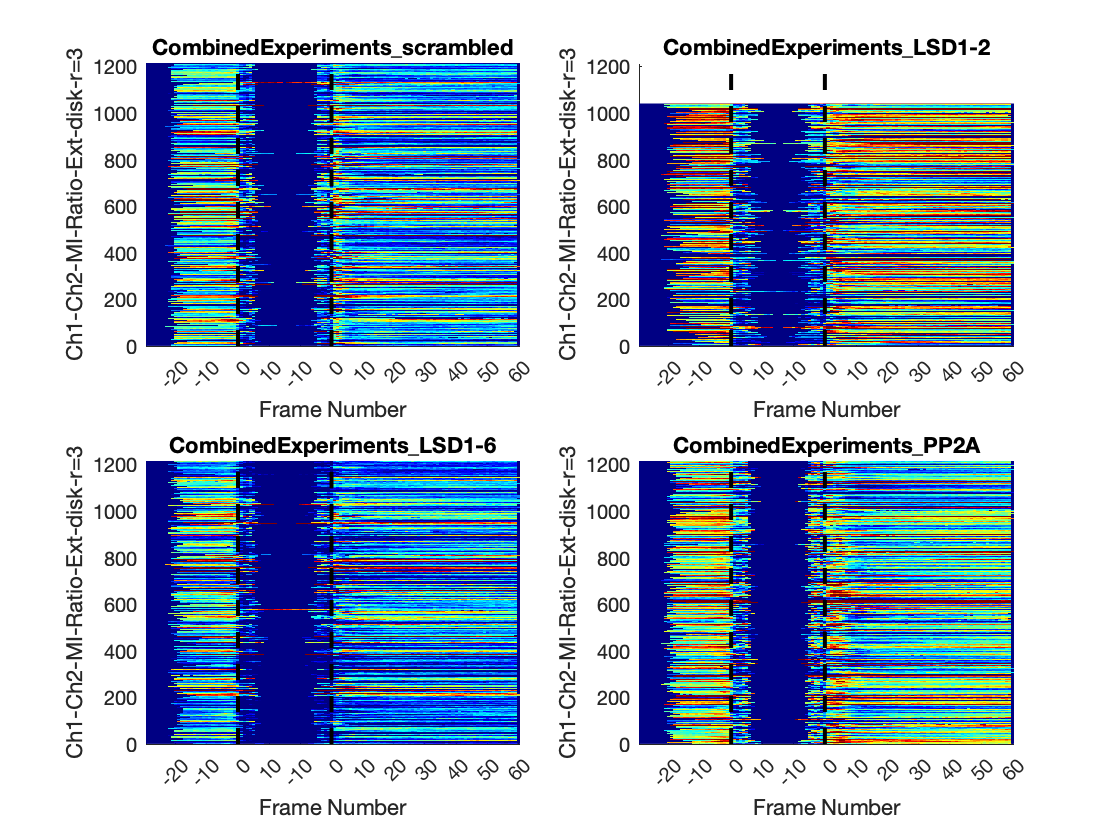

### LSD1_FusedProjects_CARSync_AdditionalFeatures_Ch1-Ch2-MI-Ratio-Ext-disk-r=3_LinePlots.png

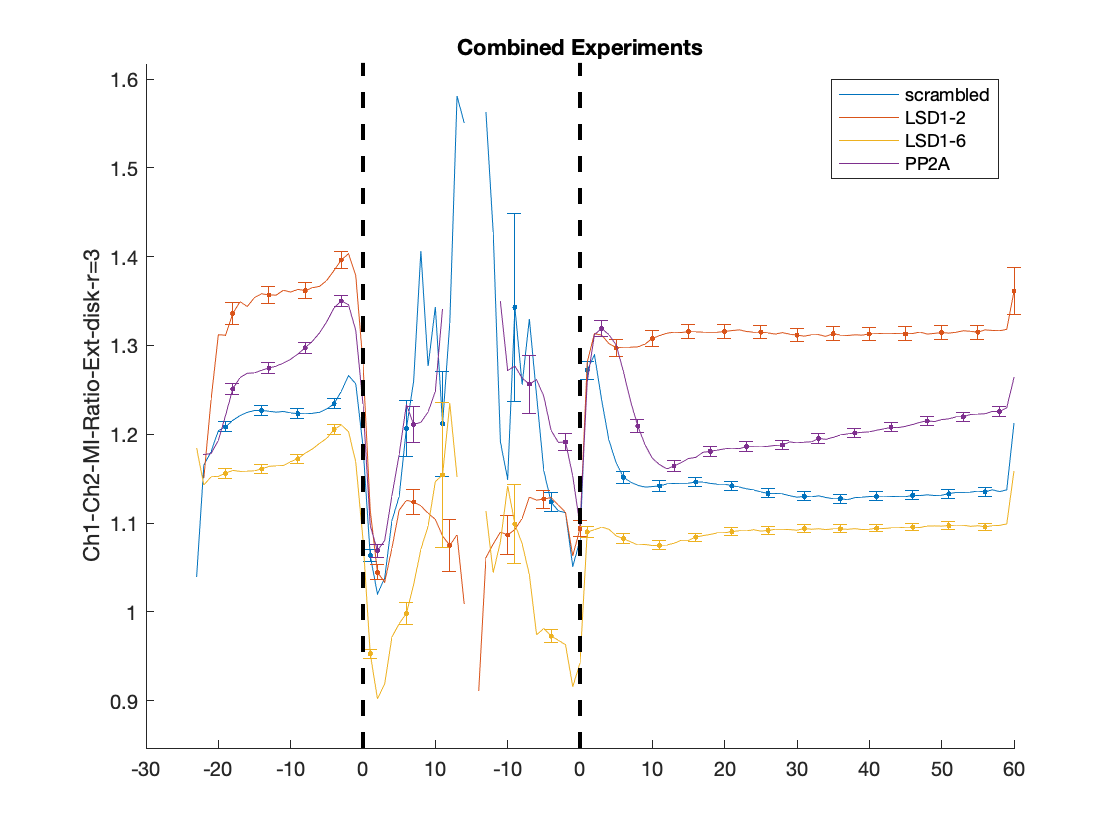

### LSD1_FusedProjects_CARSync_AdditionalFeatures_Ch2-MaxIntensity-Ext-disk-r=3_HeatMaps.png

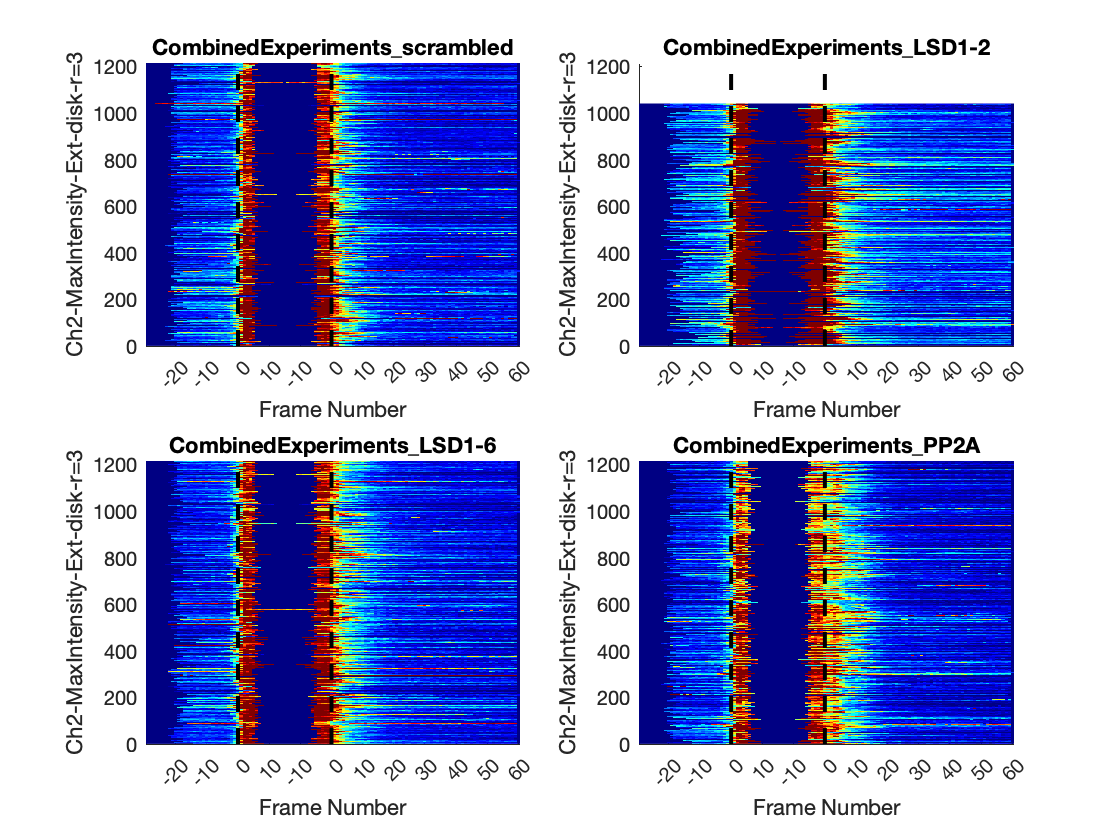

### LSD1_FusedProjects_CARSync_AdditionalFeatures_Ch2-MaxIntensity-Ext-disk-r=3_LinePlots.png

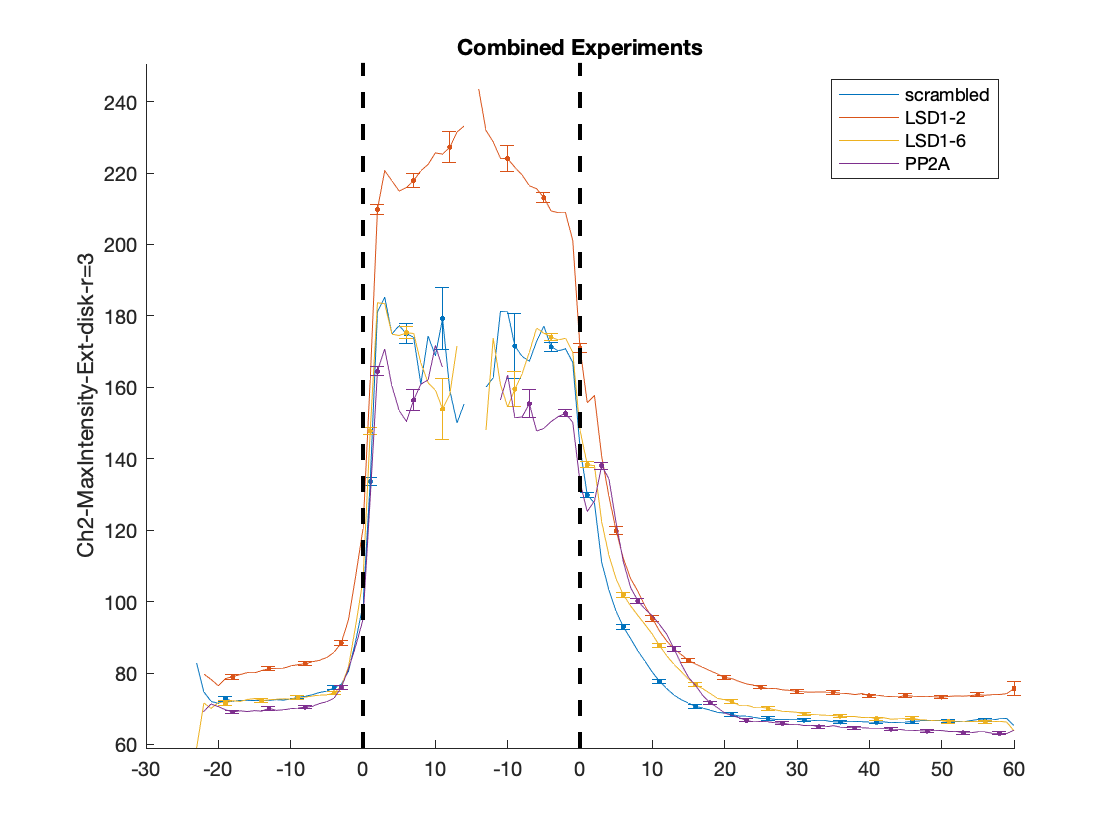

### LSD1_FusedProjects_CARSync_AdditionalFeatures_Ch2-MeanIntensity-Ext-disk-r=3-atiMean_BoxPlots.png

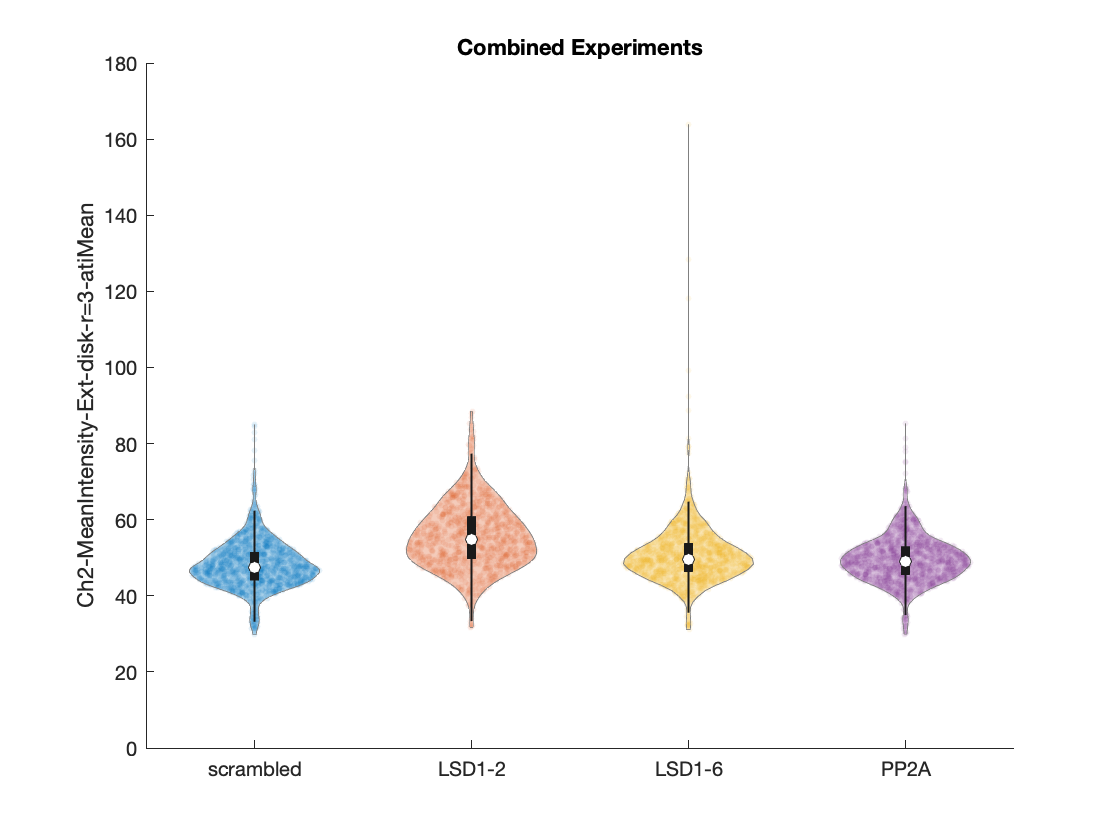

### LSD1_FusedProjects_CARSync_AdditionalFeatures_Ch2-MeanIntensity-Ext-disk-r=3-atiMean_Histograms.png

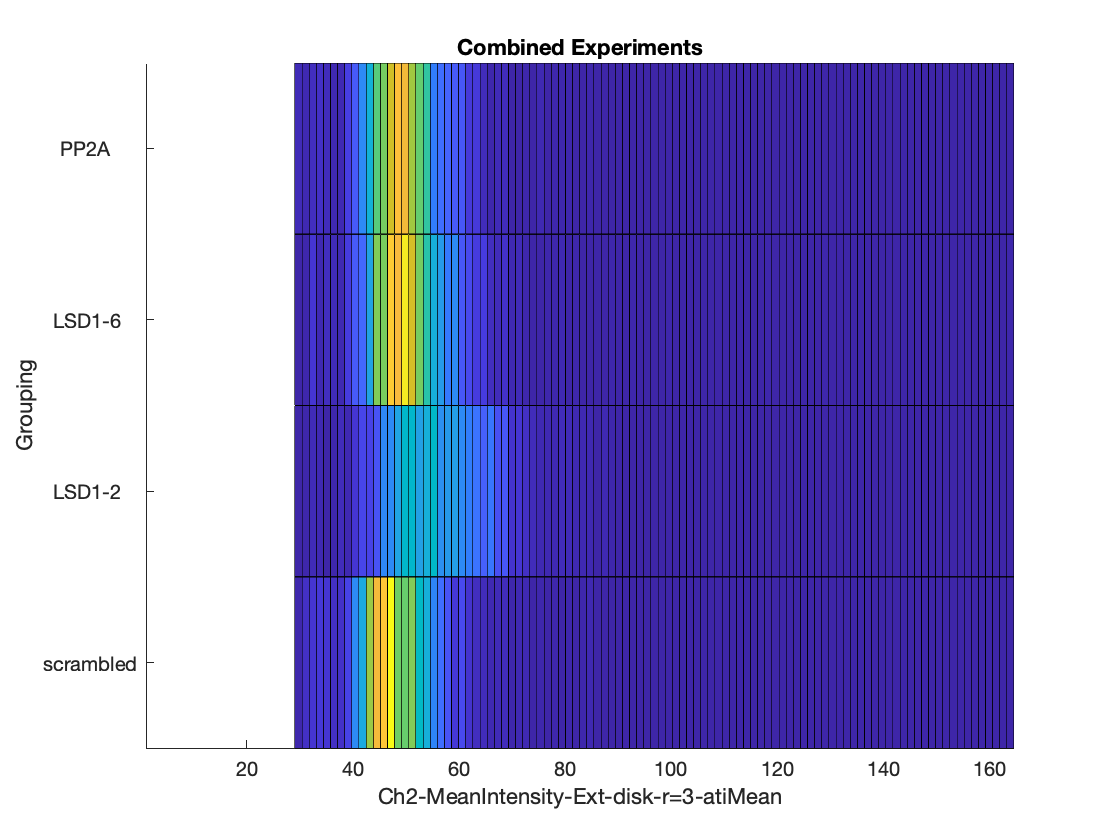

### LSD1_FusedProjects_CARSync_AdditionalFeatures_Ch2-MeanIntensity-Ext-disk-r=3-intMean_BoxPlots.png

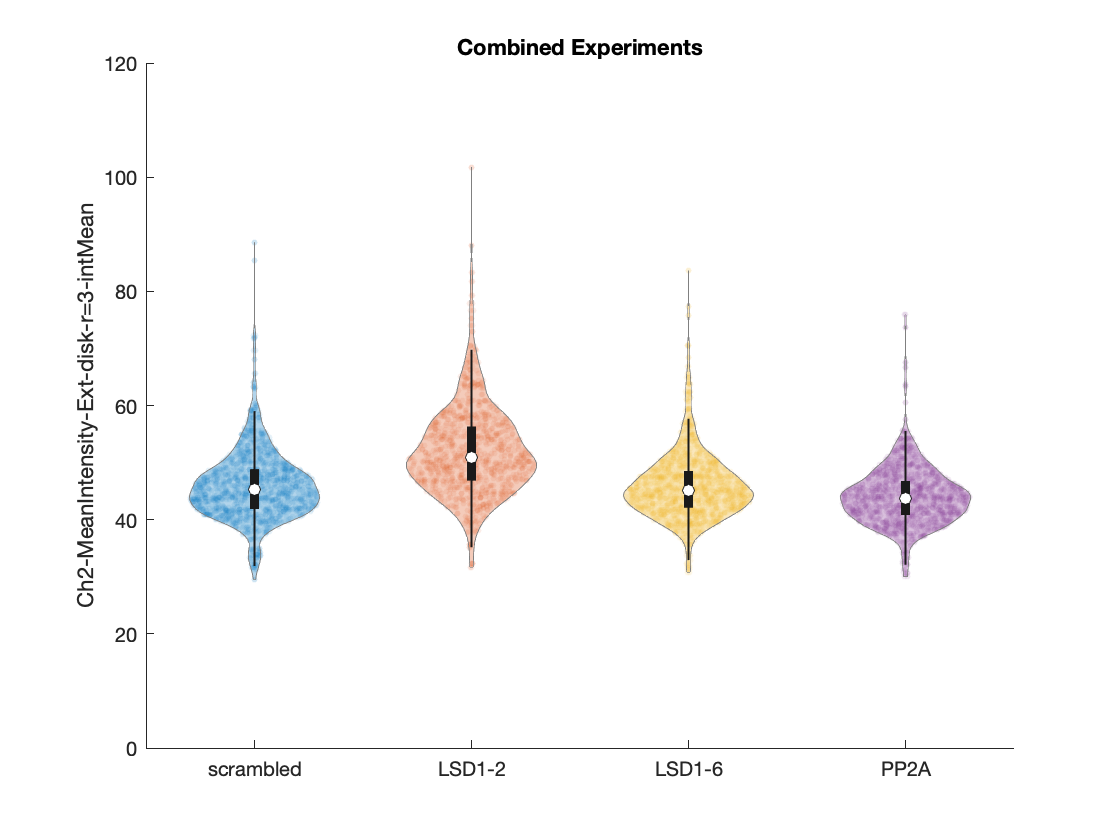

### LSD1_FusedProjects_CARSync_AdditionalFeatures_Ch2-MeanIntensity-Ext-disk-r=3-intMean_Histograms.png

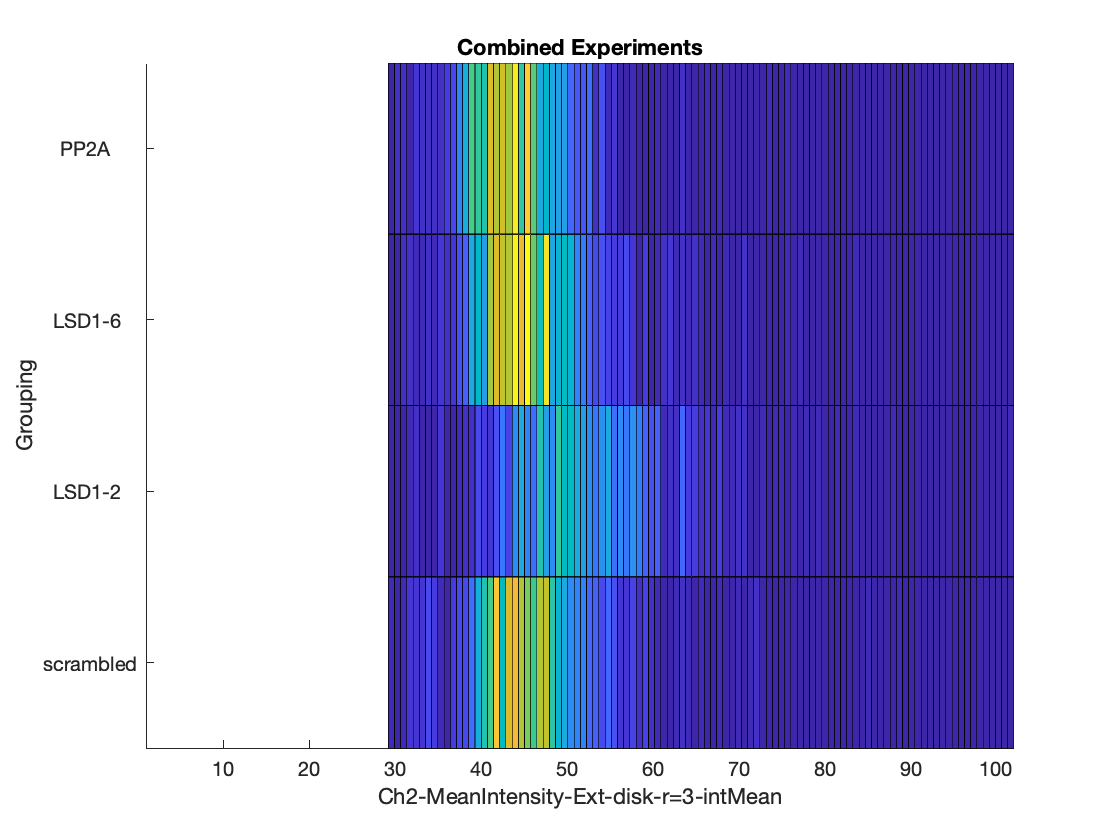

### LSD1_FusedProjects_CARSync_AdditionalFeatures_Ch2-MeanIntensity-Ext-disk-r=3-pmaMean_BoxPlots.png

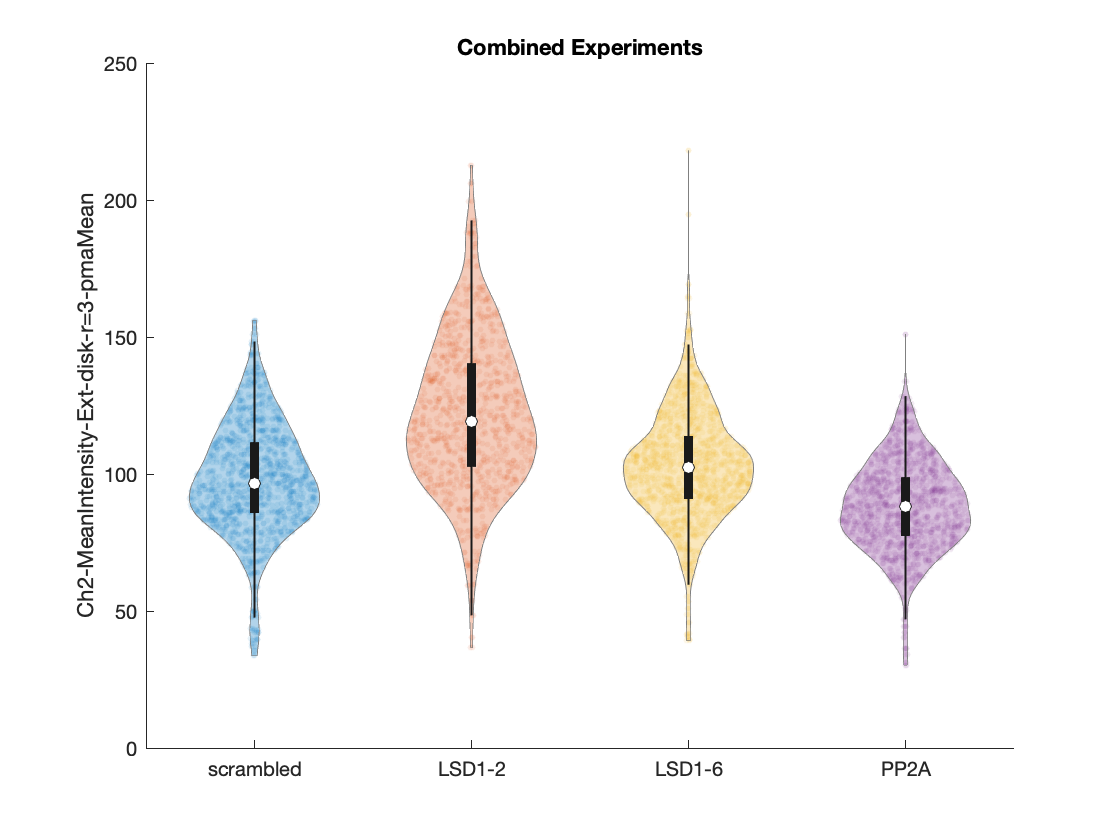

### LSD1_FusedProjects_CARSync_AdditionalFeatures_Ch2-MeanIntensity-Ext-disk-r=3-pmaMean_Histograms.png

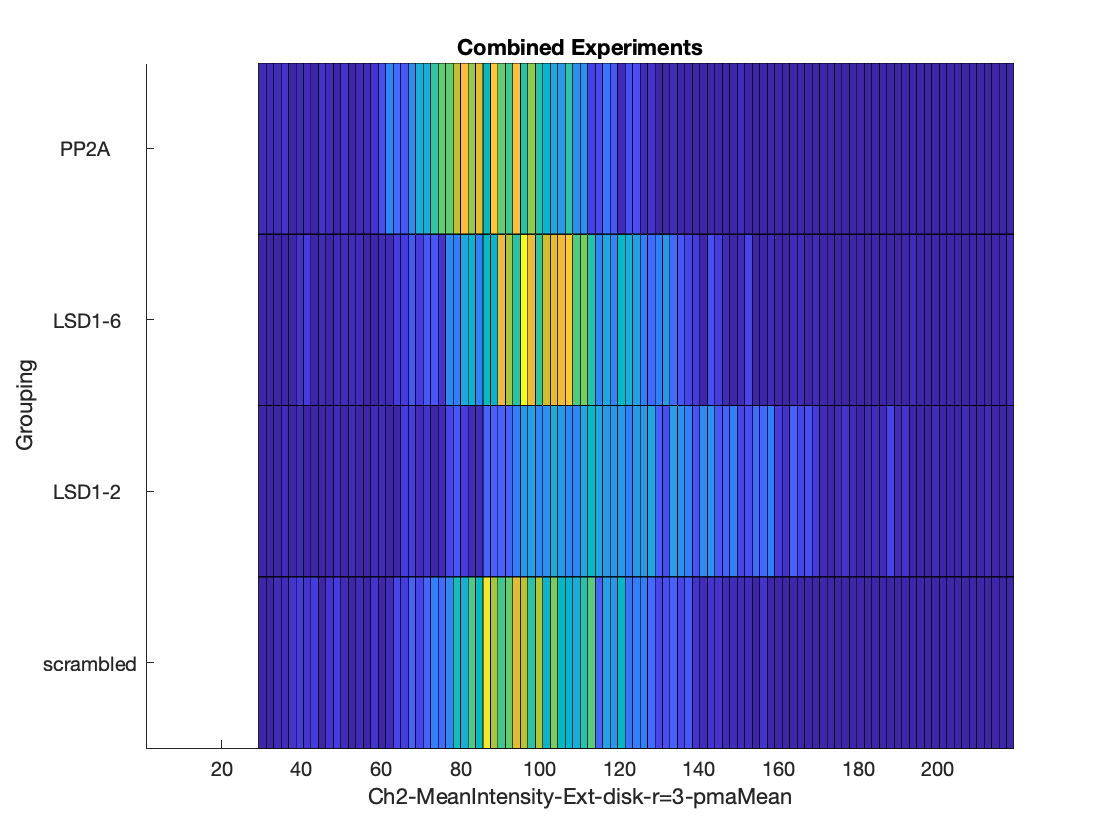

### LSD1_FusedProjects_CARSync_AdditionalFeatures_Ch2-MeanIntensity-Ext-disk-r=3_HeatMaps.png

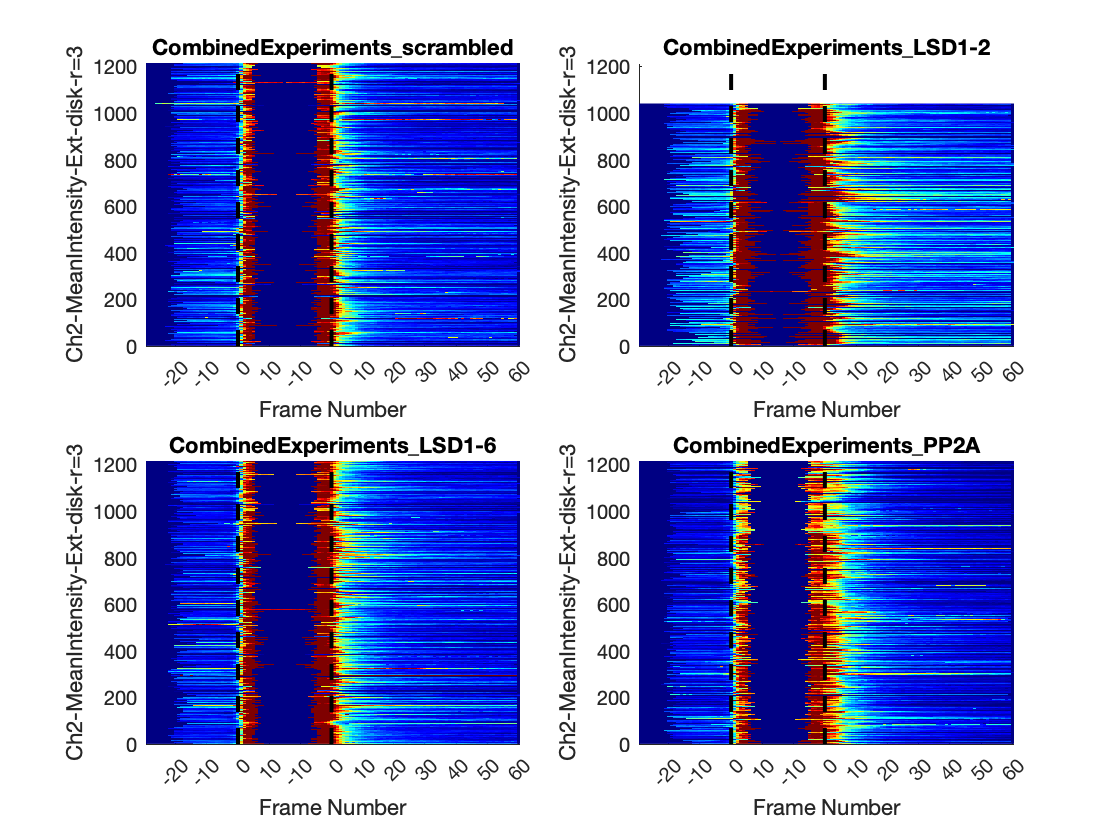

### LSD1_FusedProjects_CARSync_AdditionalFeatures_Ch2-MeanIntensity-Ext-disk-r=3_LinePlots.png

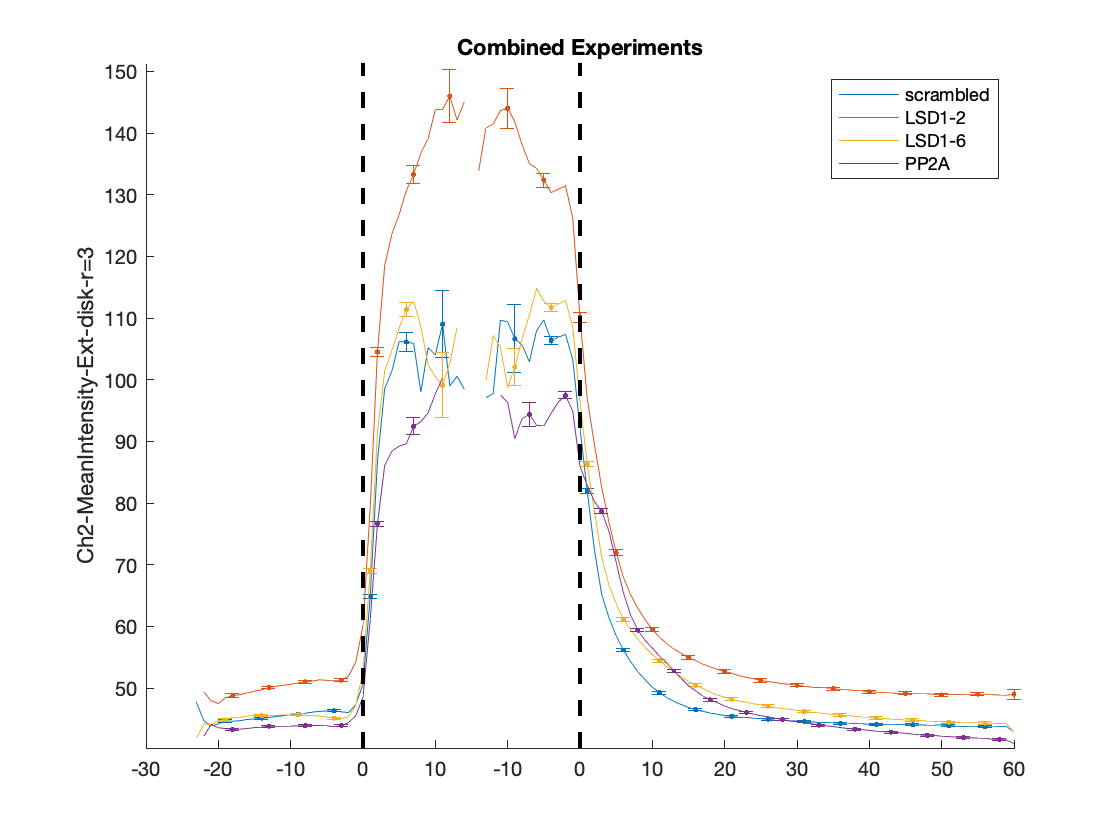

### LSD1_FusedProjects_CARSync_AdditionalFeatures_Ch2-StdIntensity-Ext-disk-r=3_HeatMaps.png

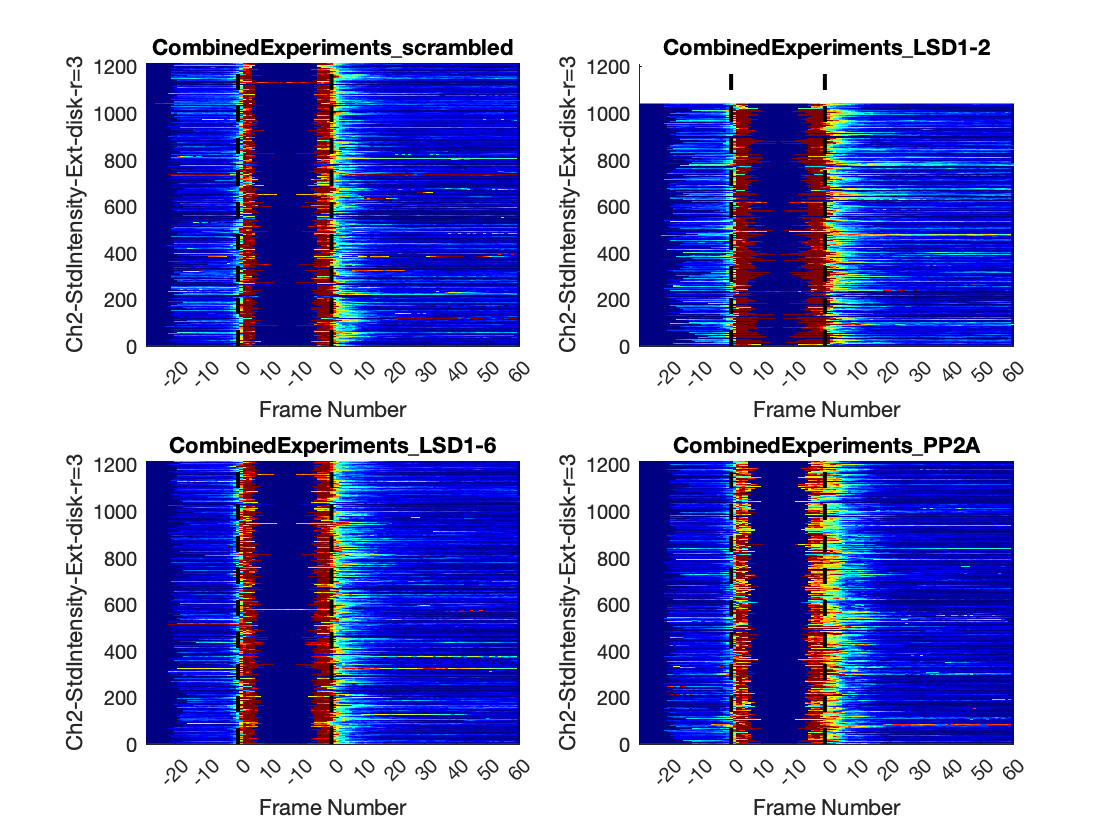

### LSD1_FusedProjects_CARSync_AdditionalFeatures_Ch2-StdIntensity-Ext-disk-r=3_LinePlots.png

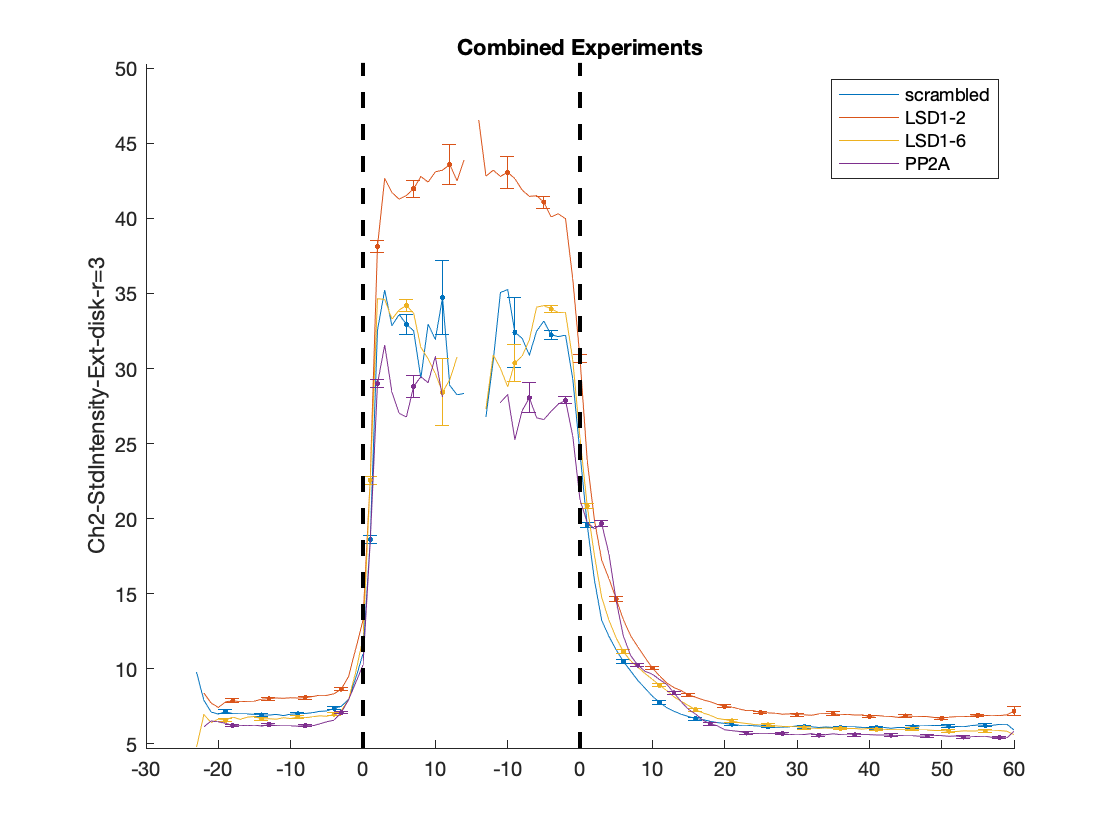

### LSD1_FusedProjects_CARSync_AdditionalFeatures_Circularity_HeatMaps.png

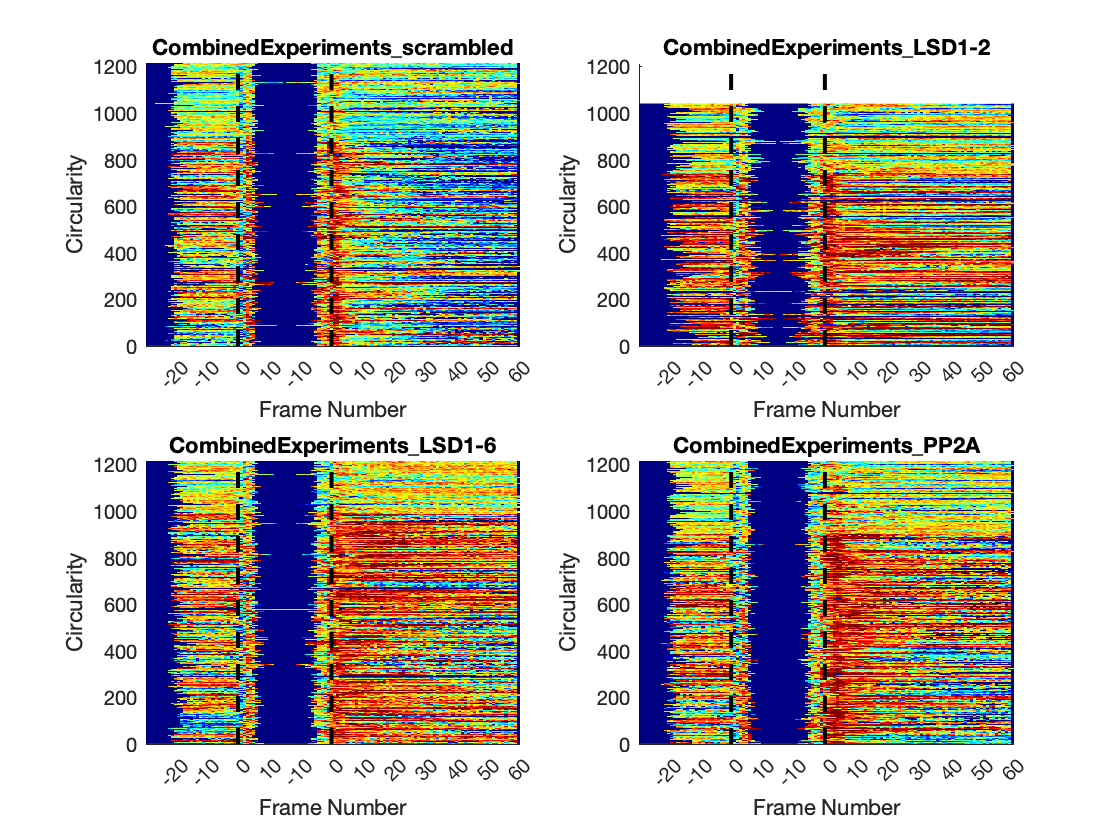

### LSD1_FusedProjects_CARSync_AdditionalFeatures_Circularity_LinePlots.png

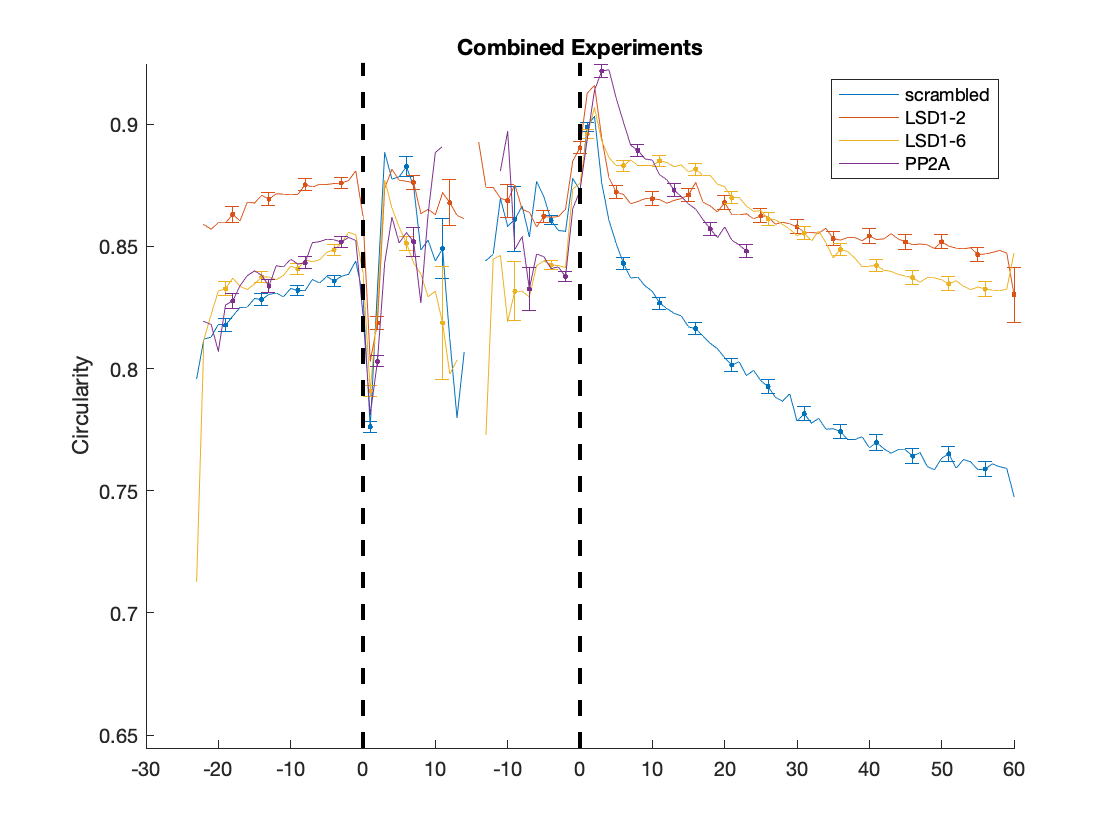

### LSD1_FusedProjects_CARSync_AdditionalFeatures_Contrast_HeatMaps.png

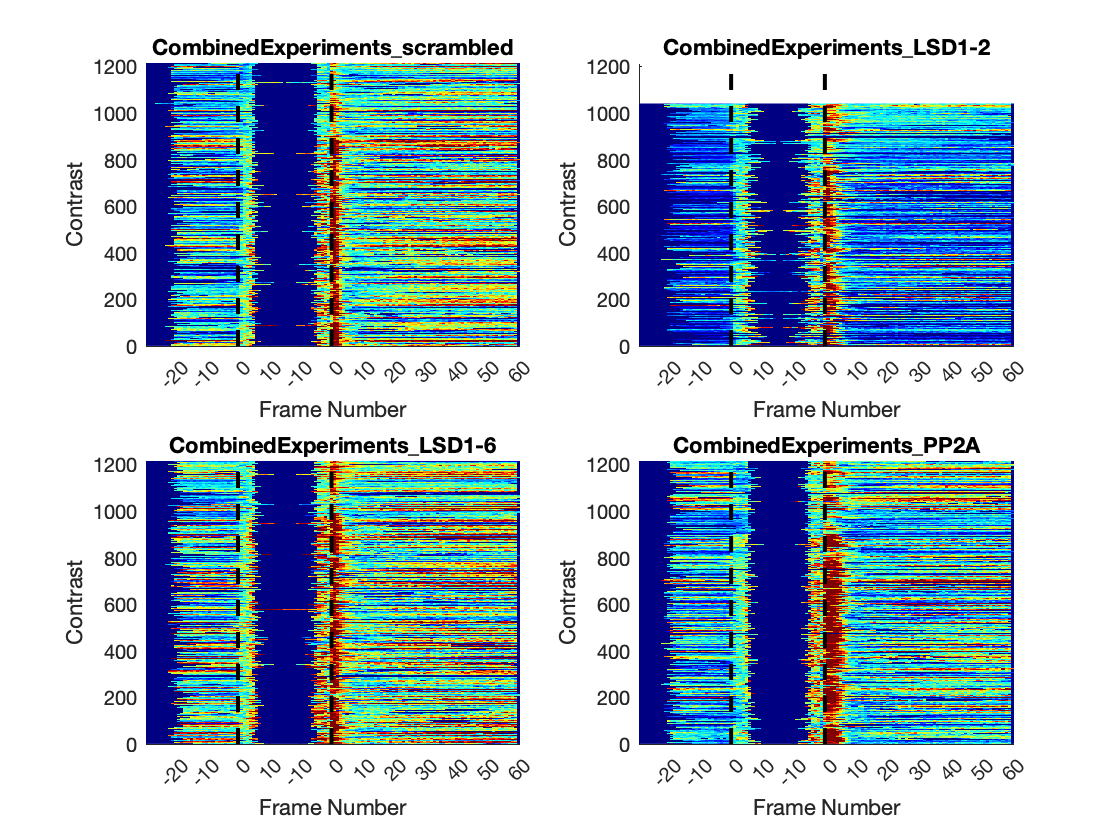

### LSD1_FusedProjects_CARSync_AdditionalFeatures_Contrast_LinePlots.png

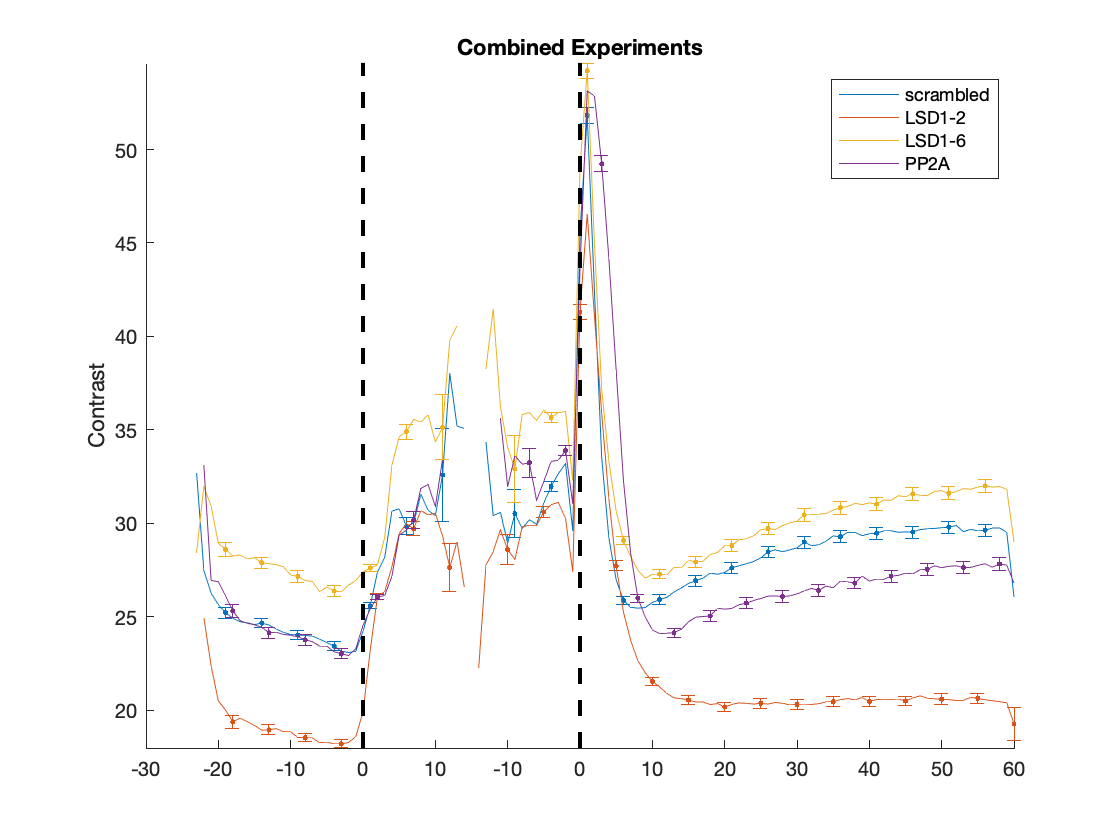

### LSD1_FusedProjects_CARSync_AdditionalFeatures_Correlation_HeatMaps.png

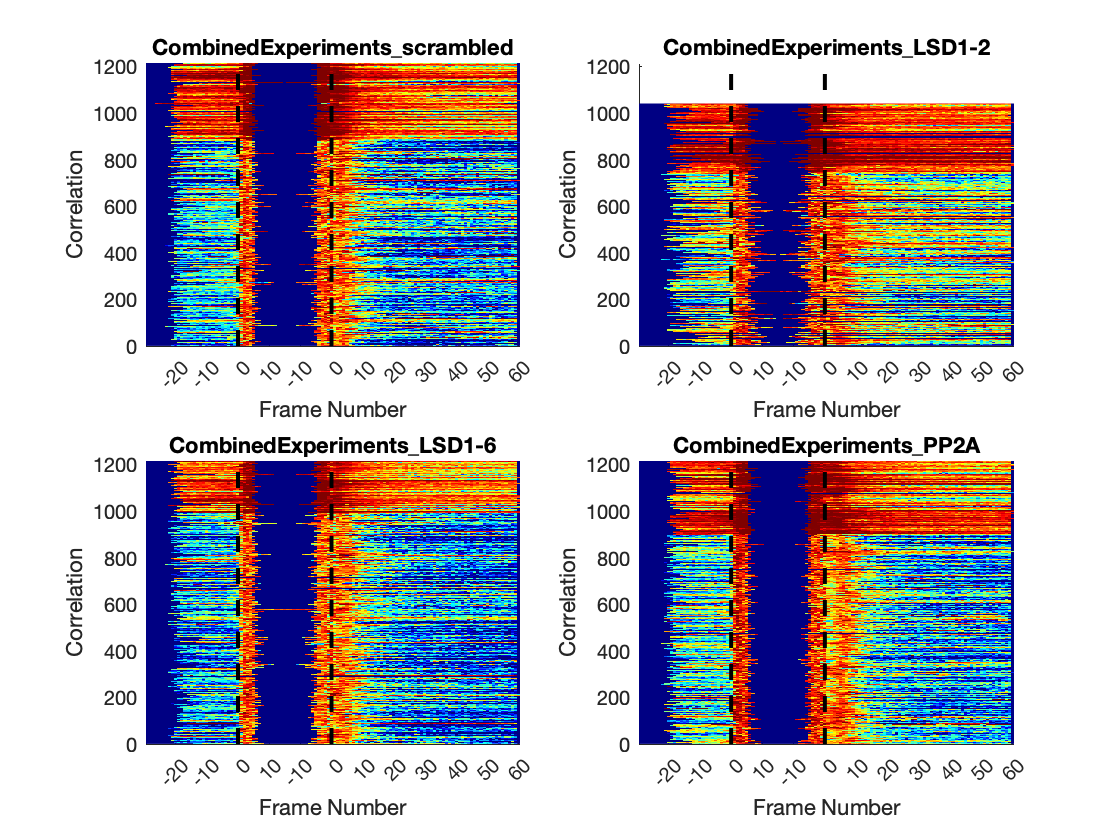

### LSD1_FusedProjects_CARSync_AdditionalFeatures_Correlation_LinePlots.png

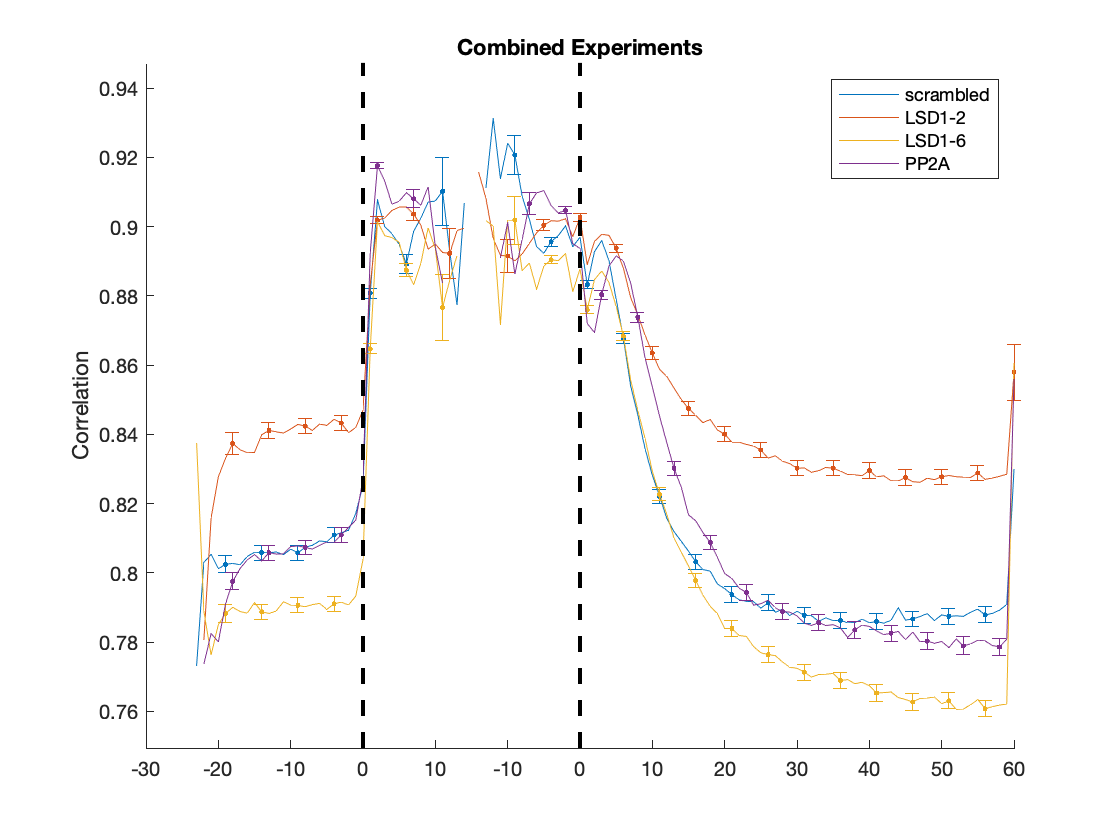

### LSD1_FusedProjects_CARSync_AdditionalFeatures_DifferenceEntropy_HeatMaps.png

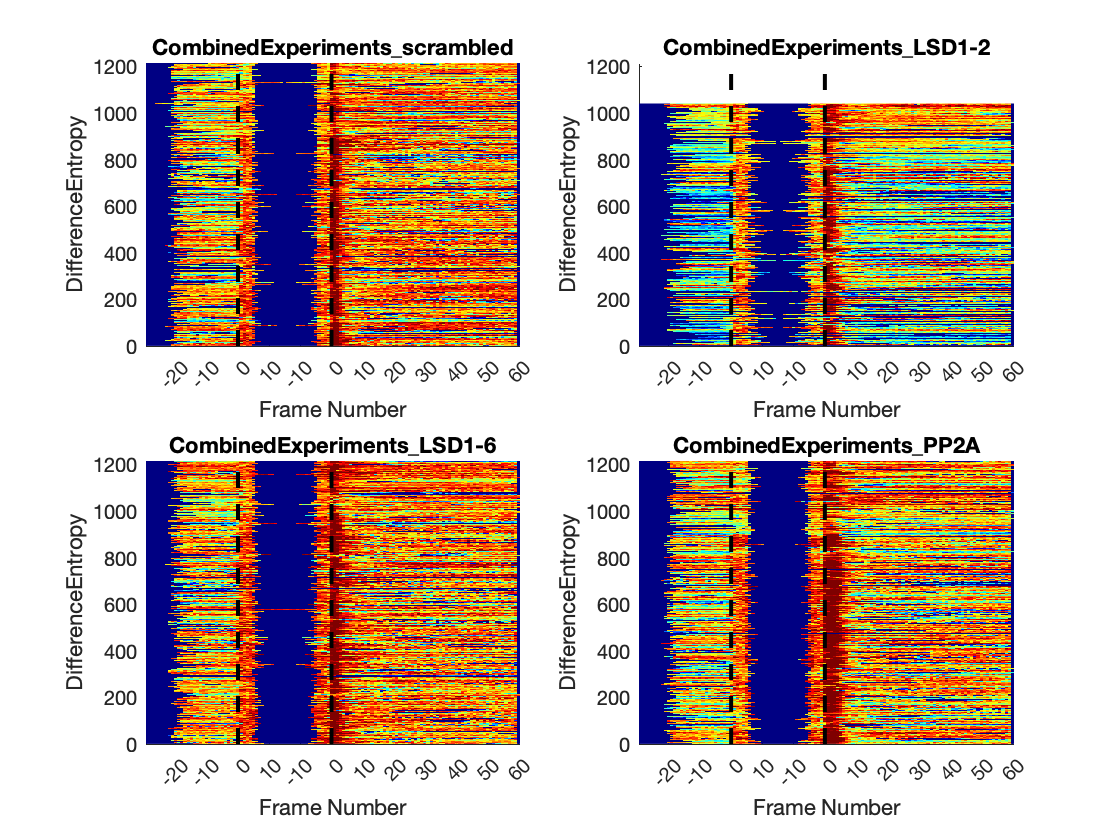

### LSD1_FusedProjects_CARSync_AdditionalFeatures_DifferenceEntropy_LinePlots.png

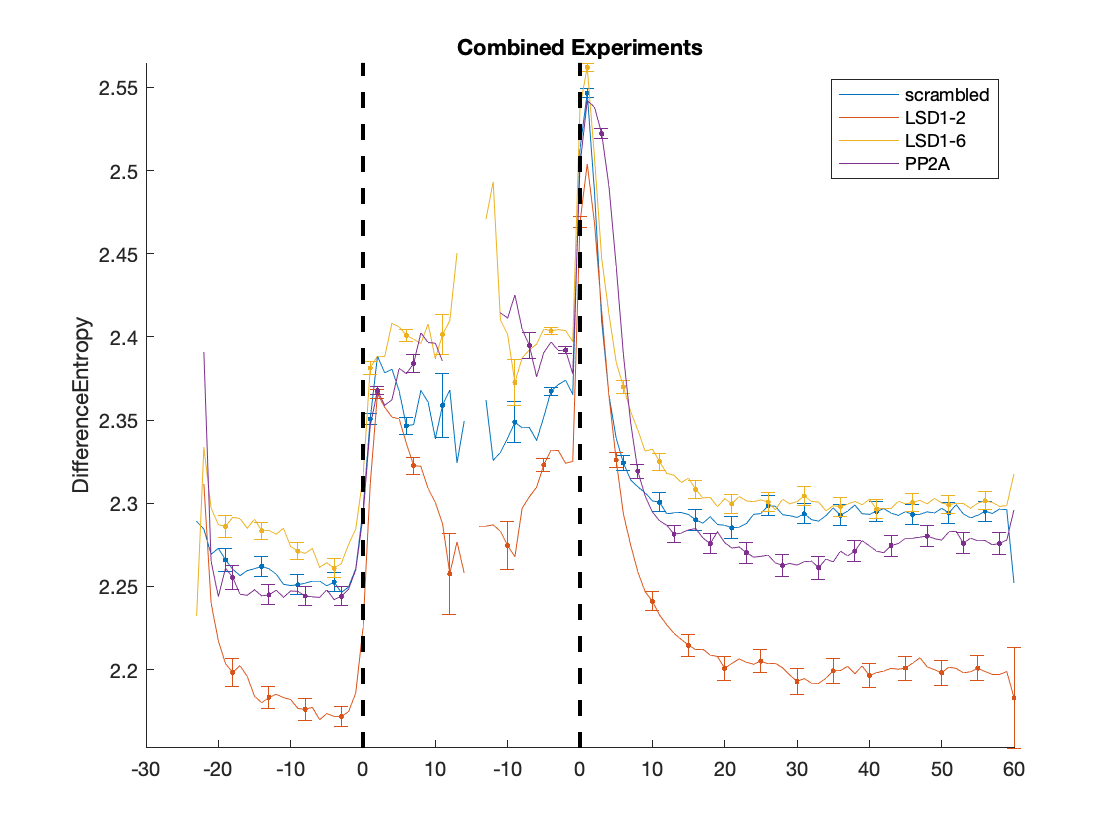

### LSD1_FusedProjects_CARSync_AdditionalFeatures_DifferenceVariance_HeatMaps.png

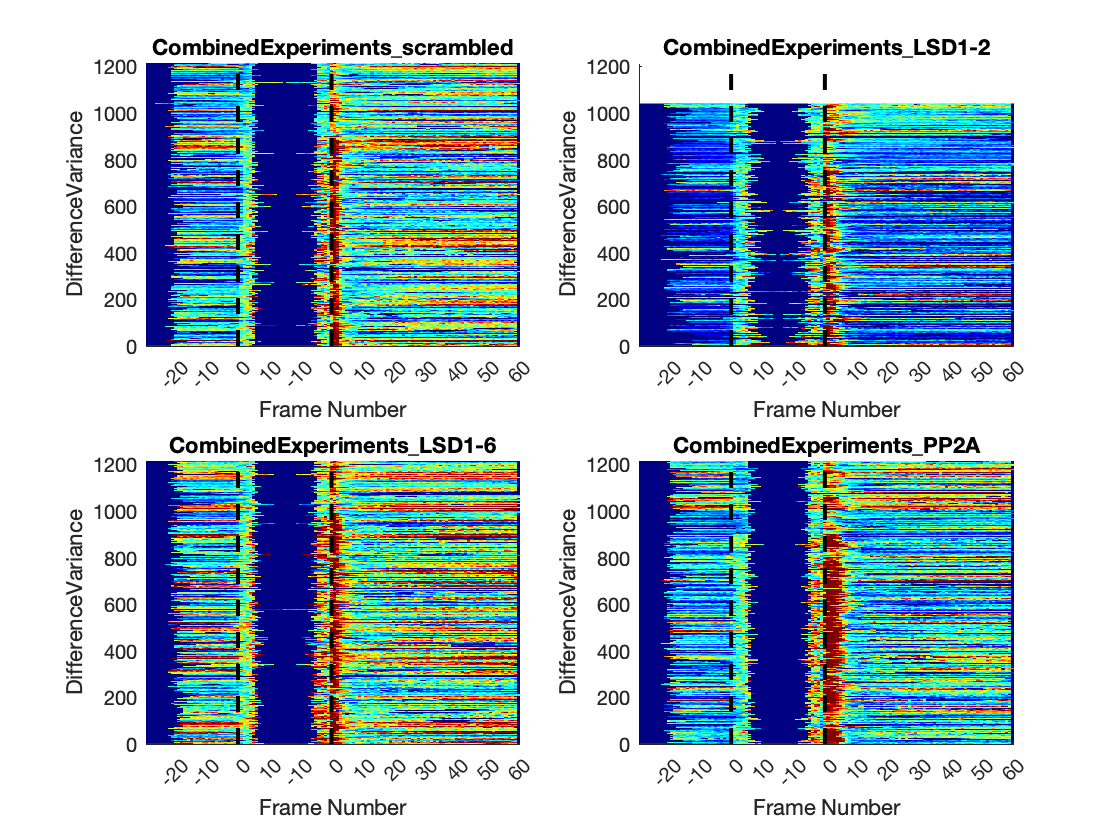

### LSD1_FusedProjects_CARSync_AdditionalFeatures_DifferenceVariance_LinePlots.png

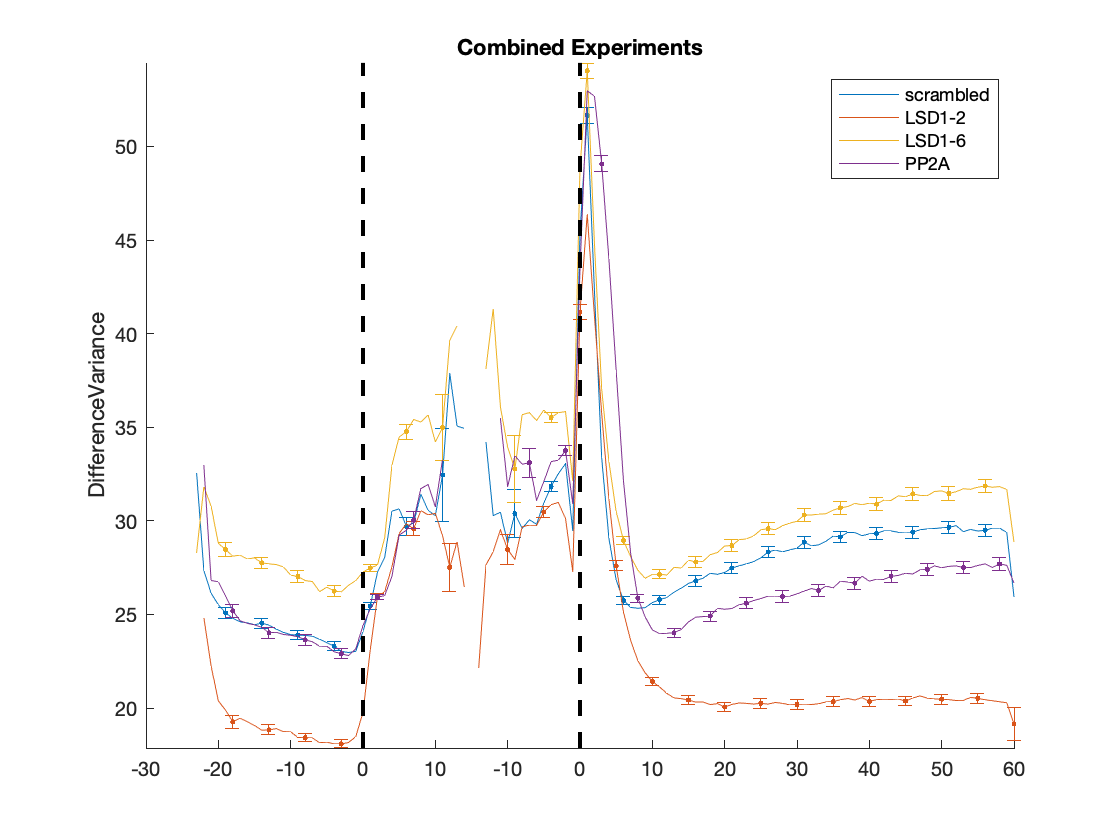

### LSD1_FusedProjects_CARSync_AdditionalFeatures_Entropy_HeatMaps.png

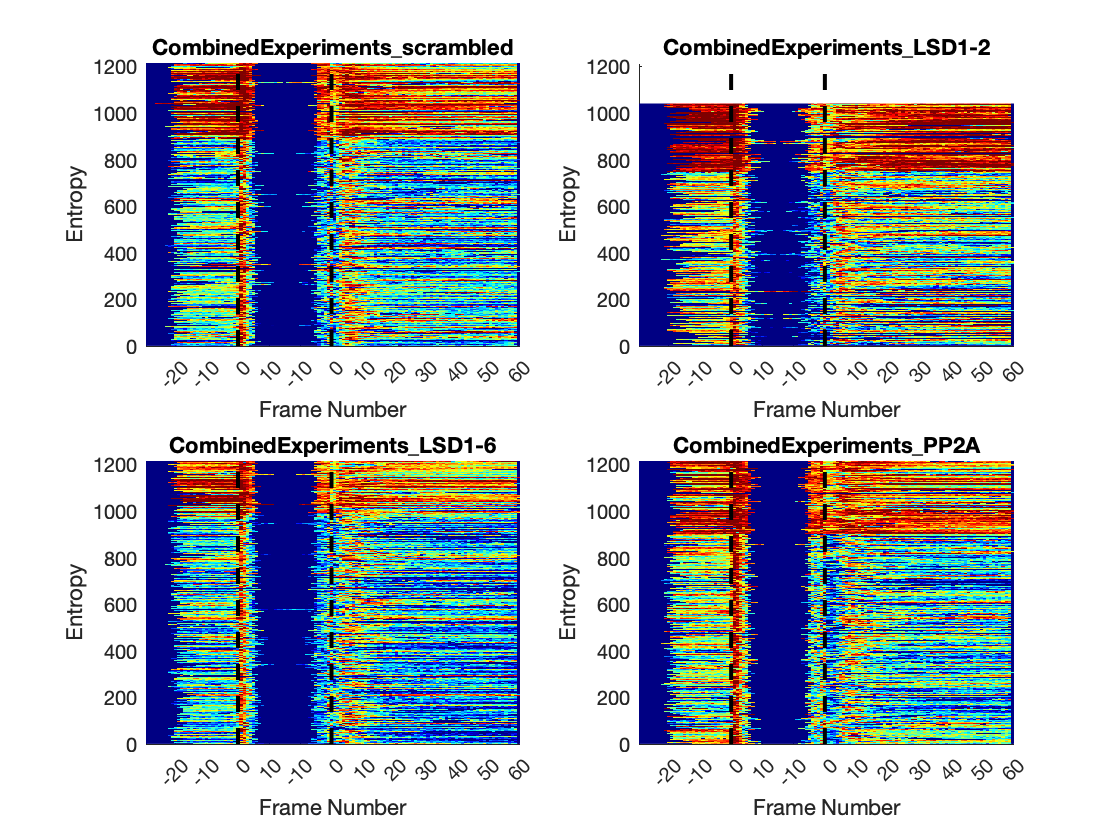

### LSD1_FusedProjects_CARSync_AdditionalFeatures_Entropy_LinePlots.png

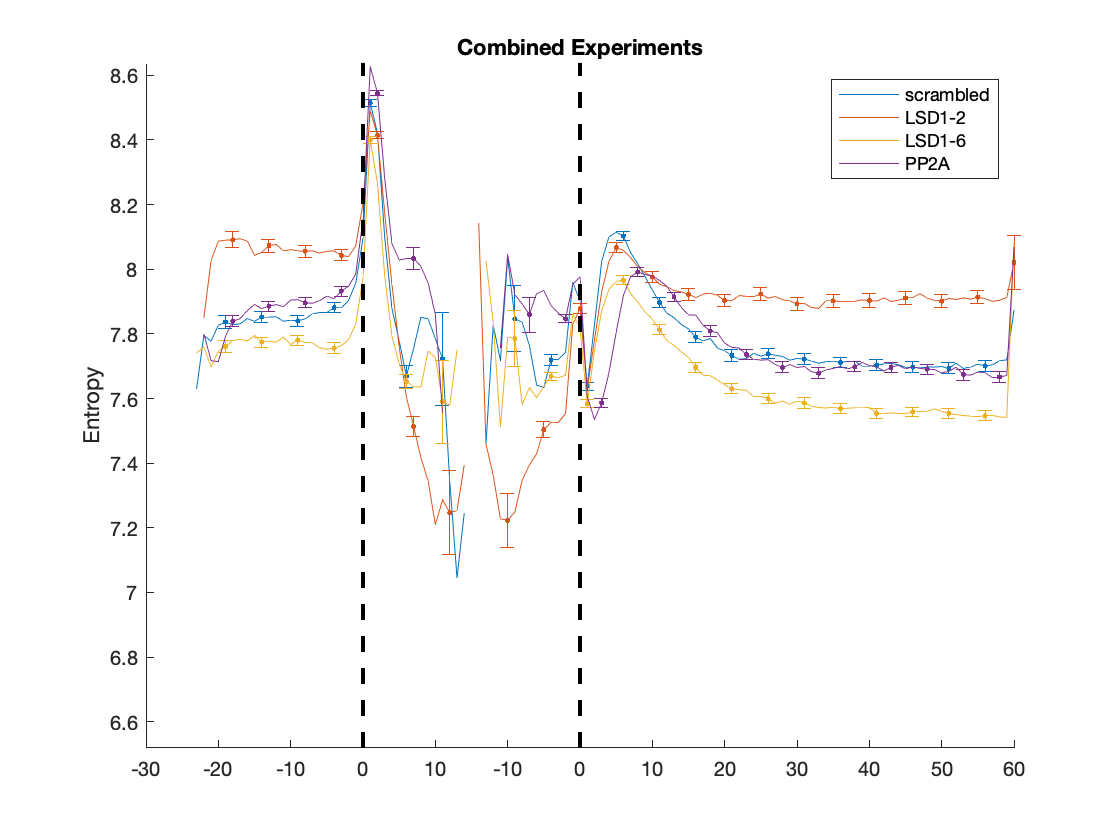

### LSD1_FusedProjects_CARSync_AdditionalFeatures_id_HeatMaps.png

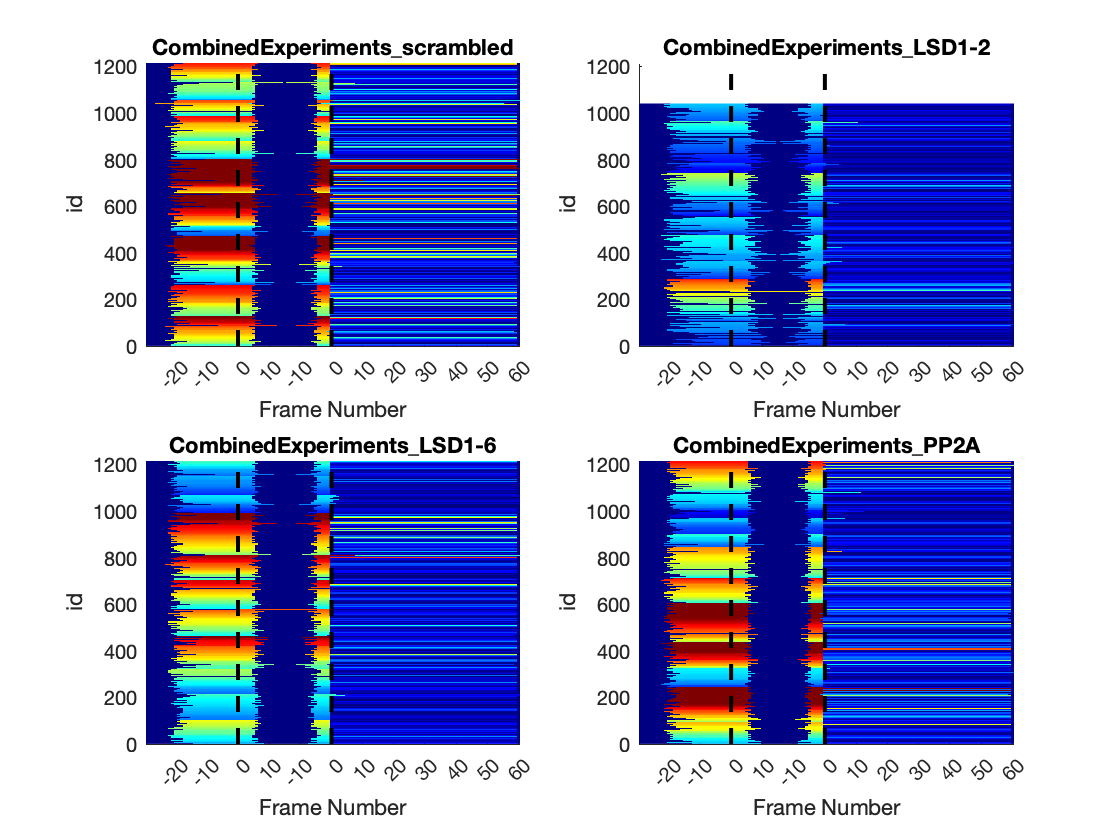

### LSD1_FusedProjects_CARSync_AdditionalFeatures_id_LinePlots.png

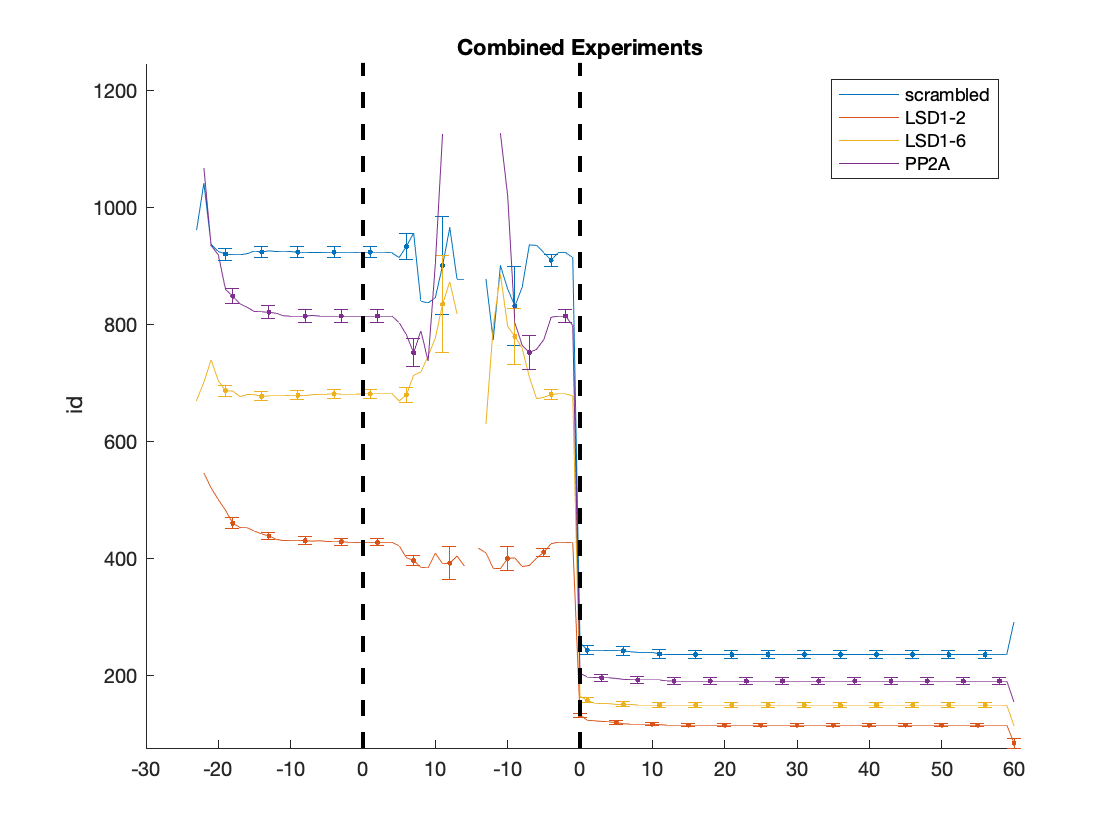

### LSD1_FusedProjects_CARSync_AdditionalFeatures_InformationMeasureofCorrelationI_HeatMaps.png

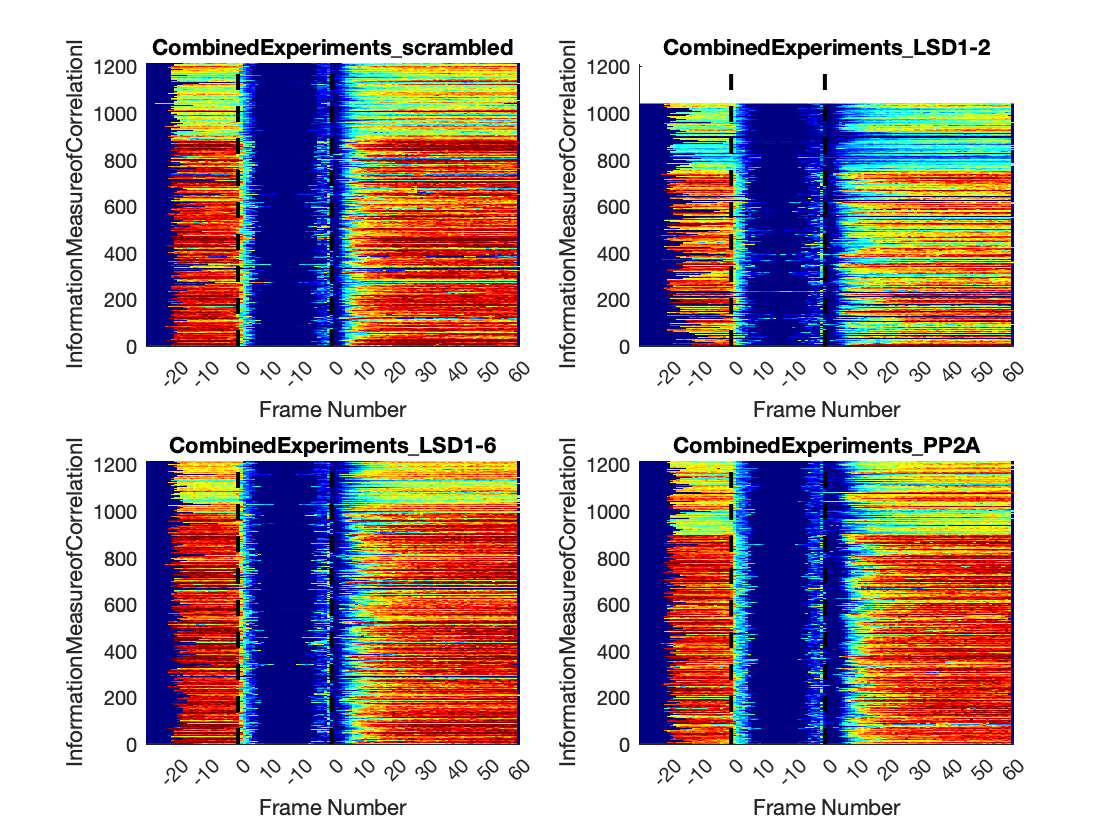

### LSD1_FusedProjects_CARSync_AdditionalFeatures_InformationMeasureofCorrelationI_LinePlots.png

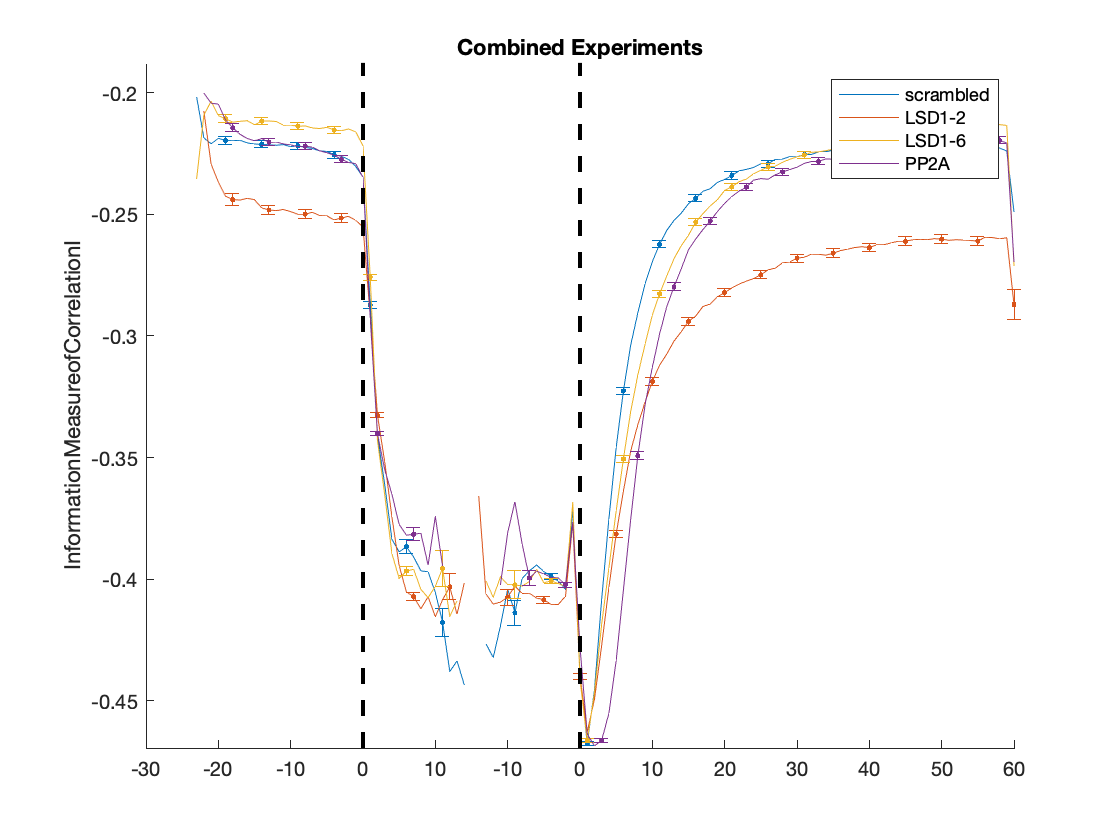

### LSD1_FusedProjects_CARSync_AdditionalFeatures_InformationMeasureofCorrelationII_HeatMaps.png

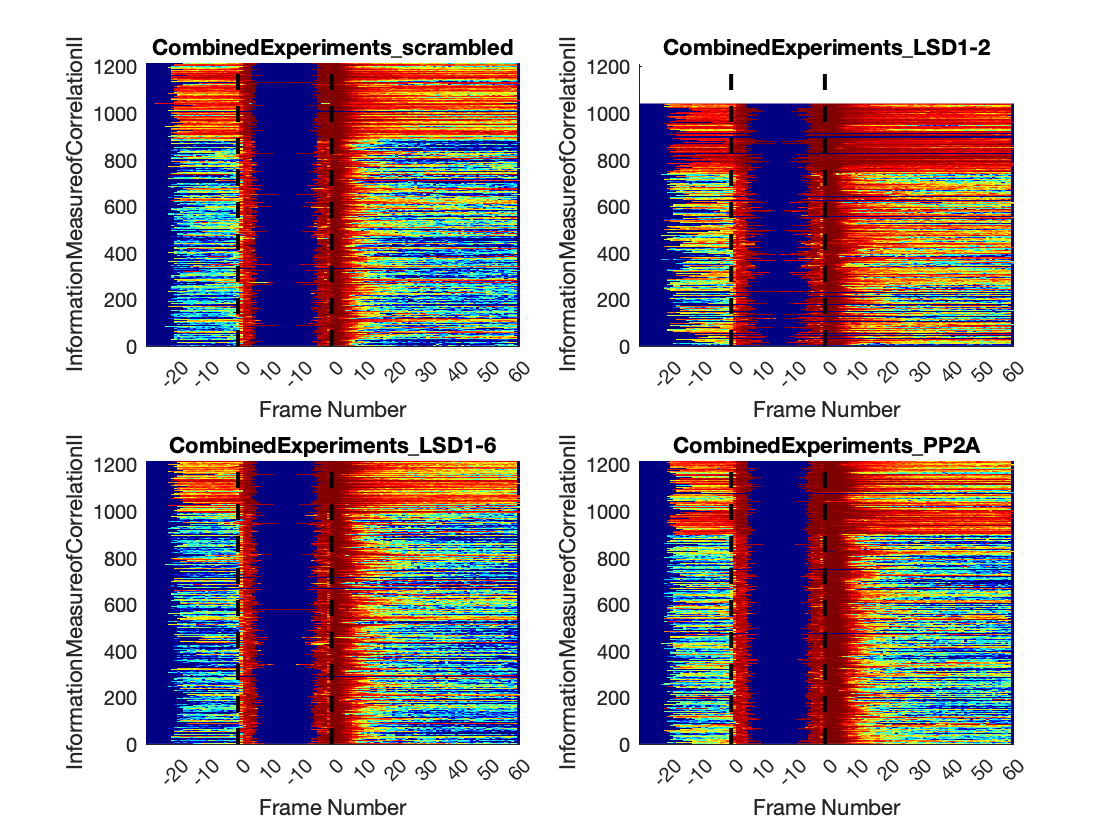

### LSD1_FusedProjects_CARSync_AdditionalFeatures_InformationMeasureofCorrelationII_LinePlots.png

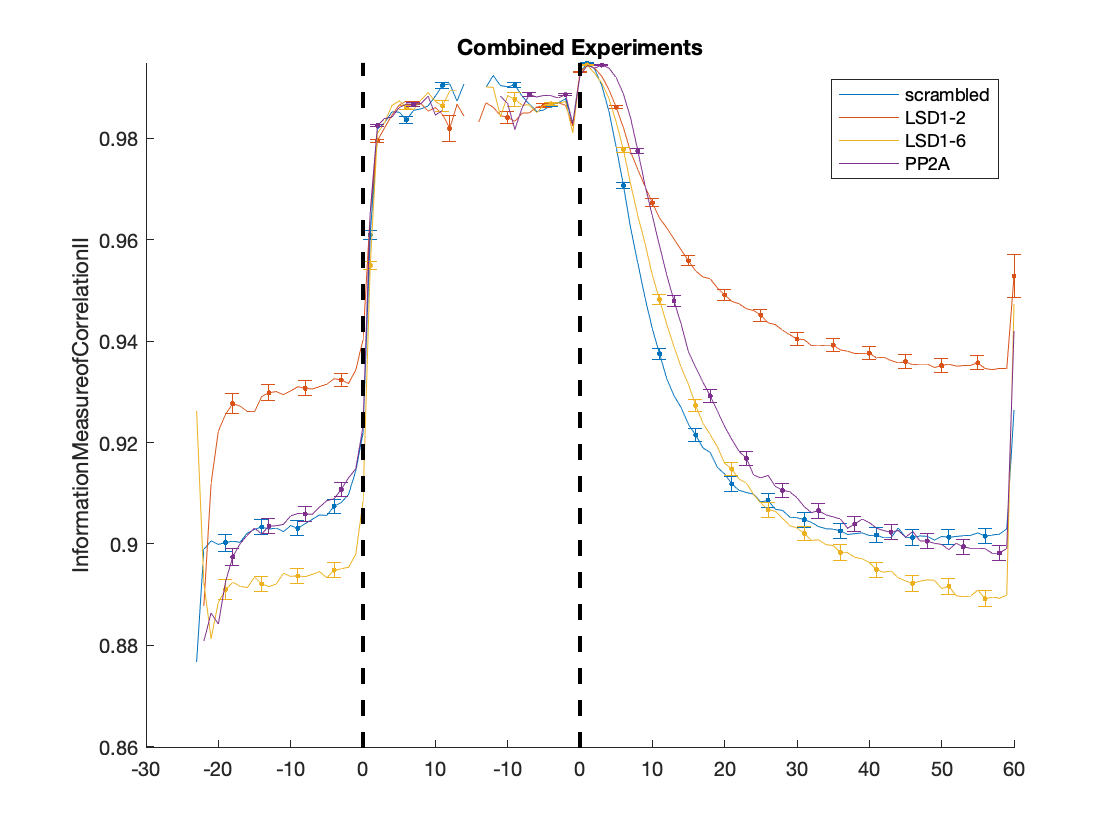

### LSD1_FusedProjects_CARSync_AdditionalFeatures_InterphaseMeanIntensity_BoxPlots.png

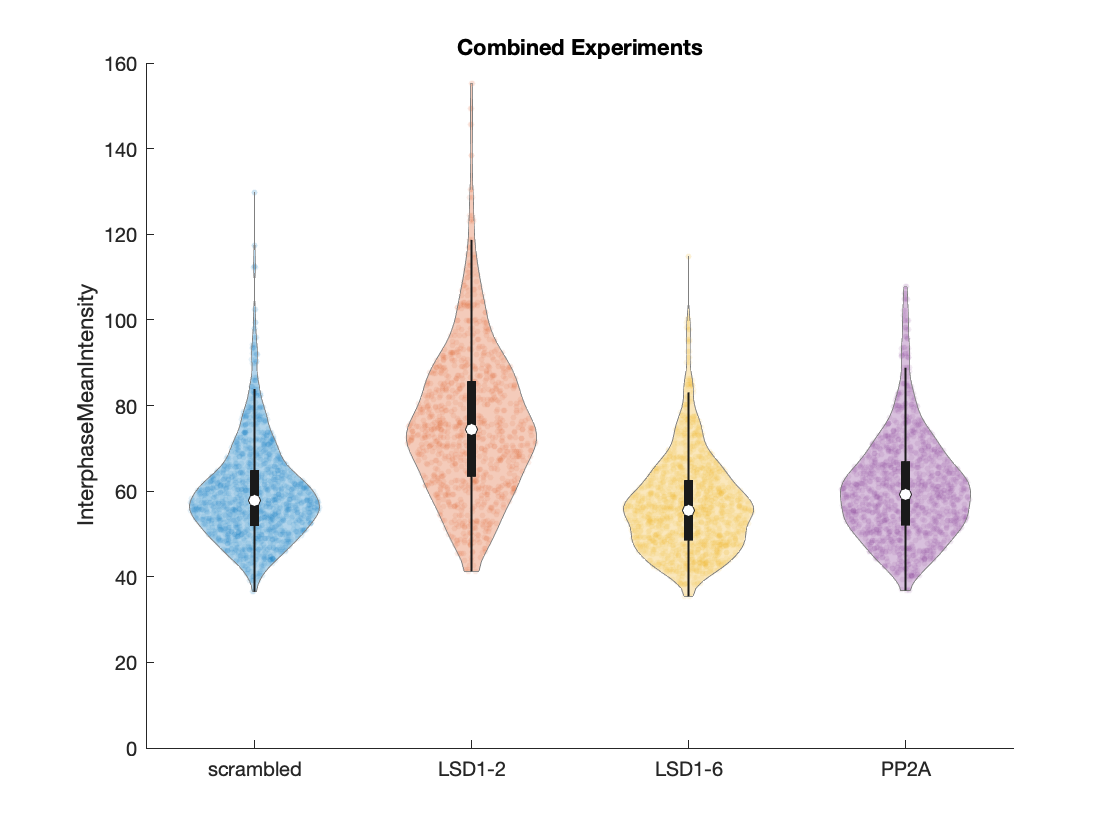

### LSD1_FusedProjects_CARSync_AdditionalFeatures_InterphaseMeanIntensity_Histograms.png

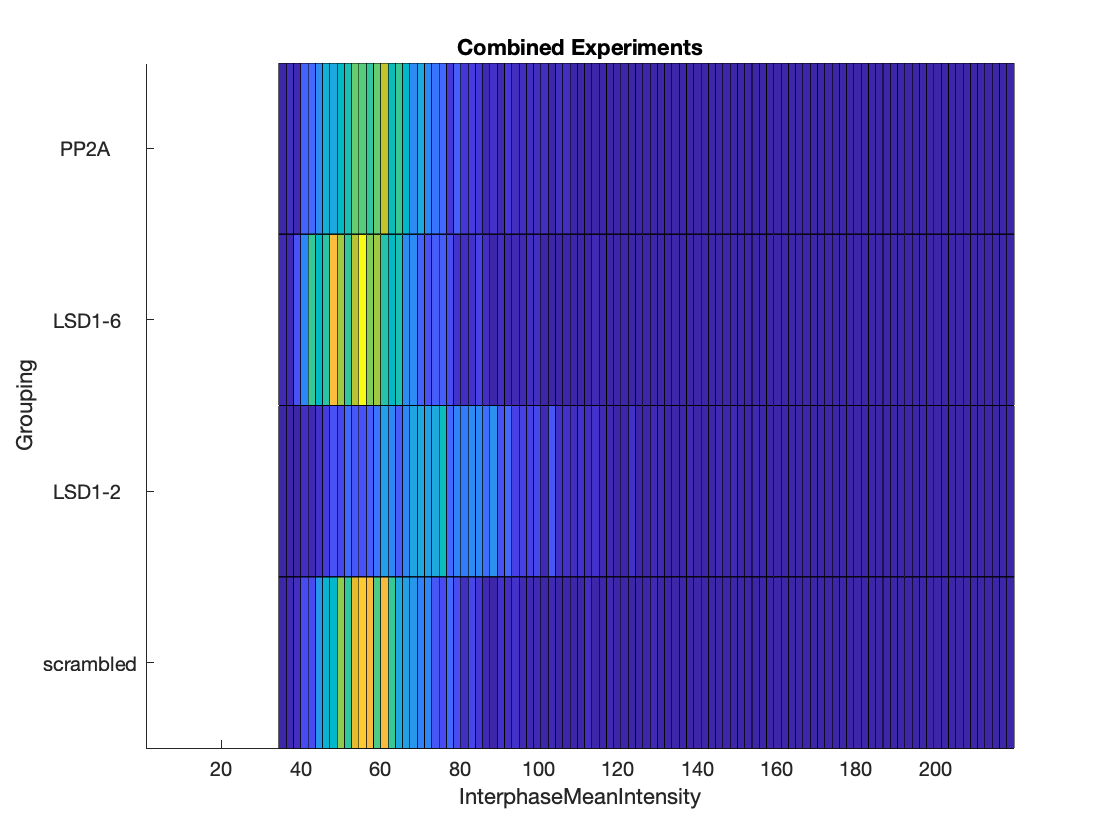

### LSD1_FusedProjects_CARSync_AdditionalFeatures_InverseDifferenceMoment(Homogeneity)_HeatMaps.png

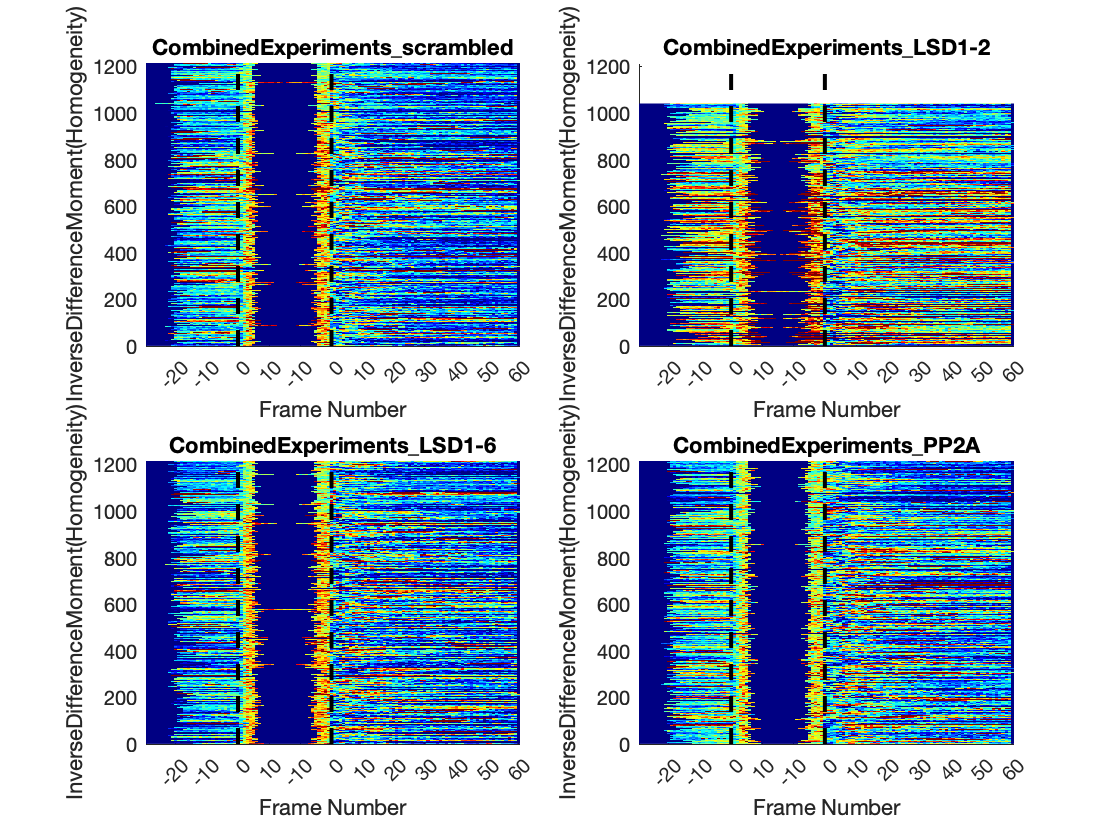

### LSD1_FusedProjects_CARSync_AdditionalFeatures_InverseDifferenceMoment(Homogeneity)_LinePlots.png

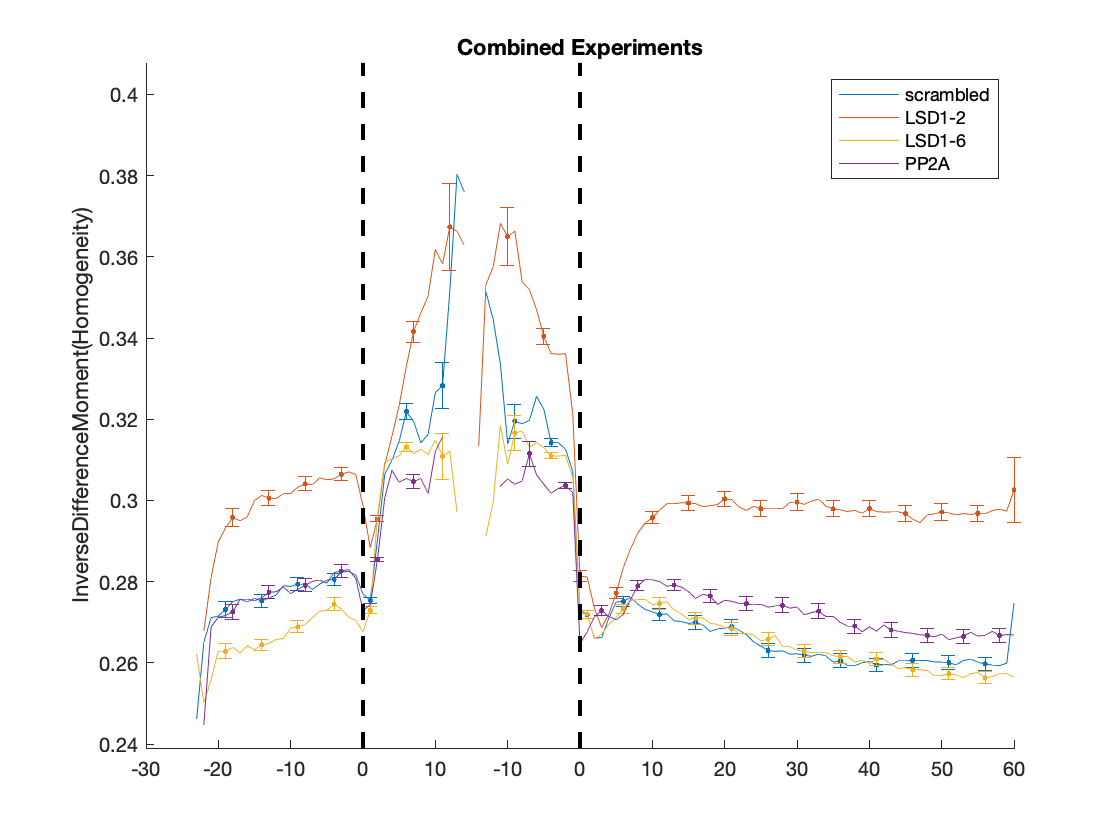

### LSD1_FusedProjects_CARSync_AdditionalFeatures_IPToMALength_Frames_BoxPlots.png

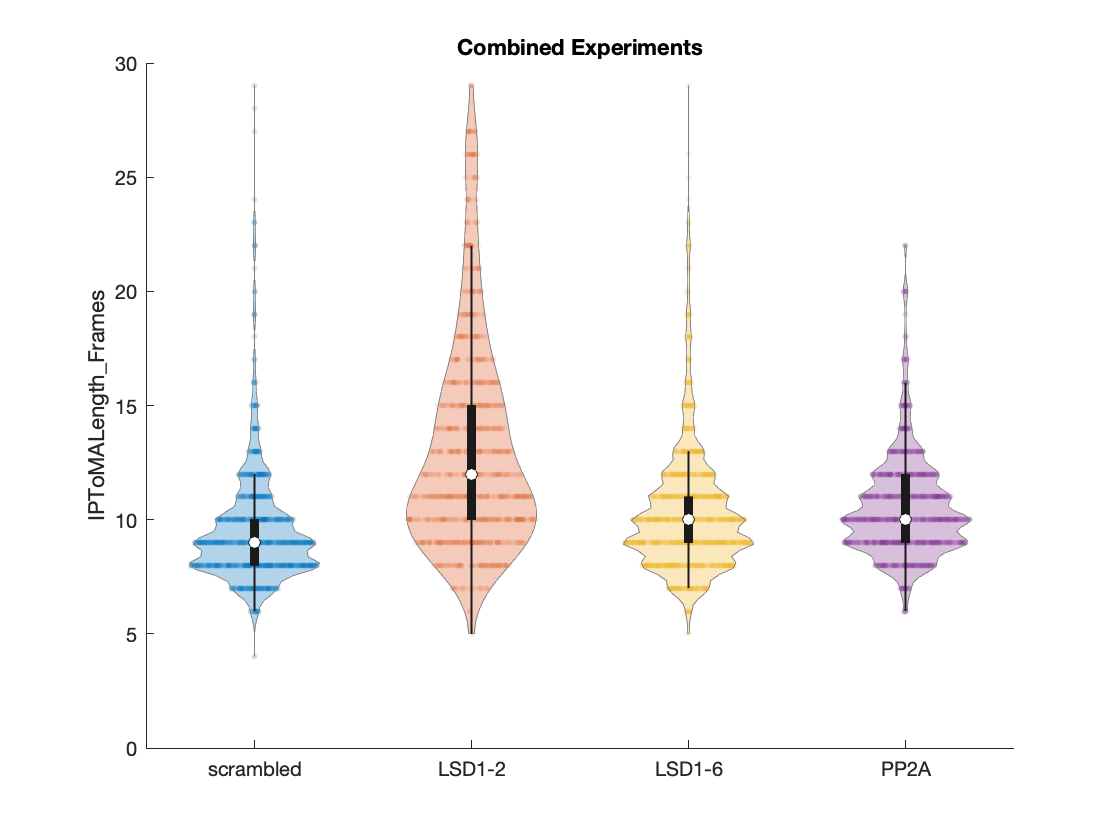

### LSD1_FusedProjects_CARSync_AdditionalFeatures_IPToMALength_Frames_Histograms.png

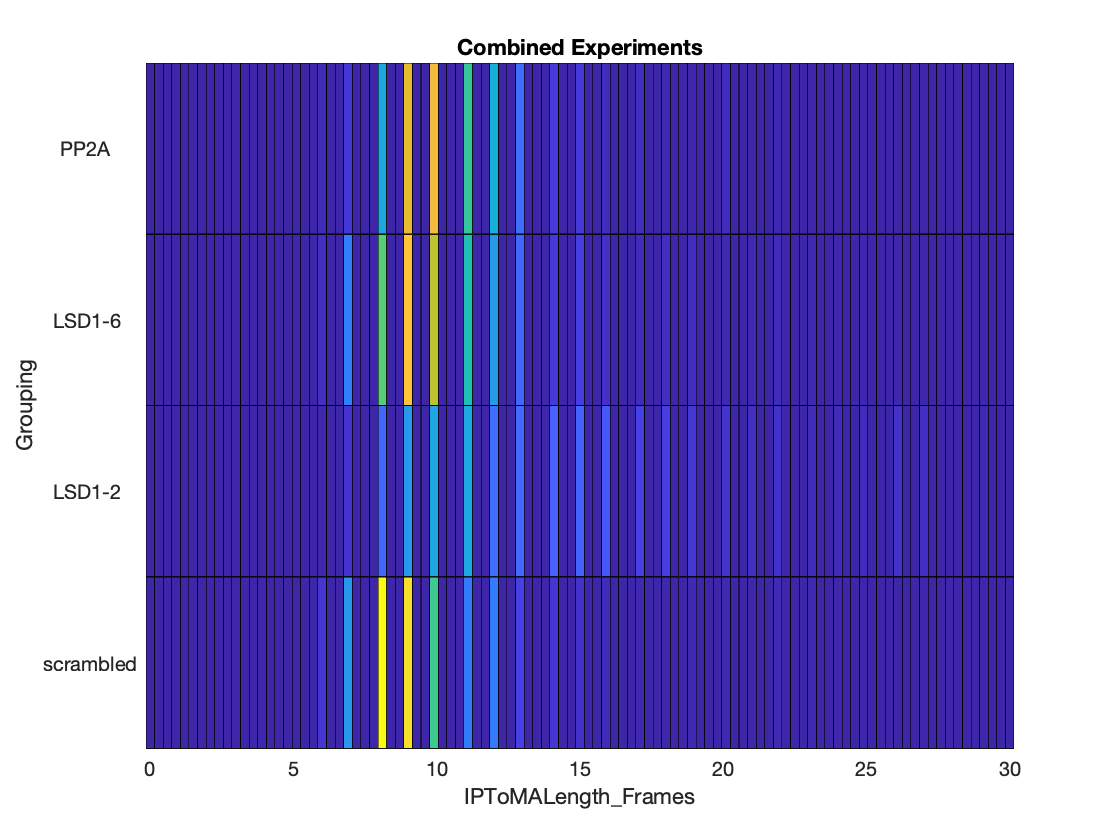

### LSD1_FusedProjects_CARSync_AdditionalFeatures_IPToMALength_Minutes_BoxPlots.png

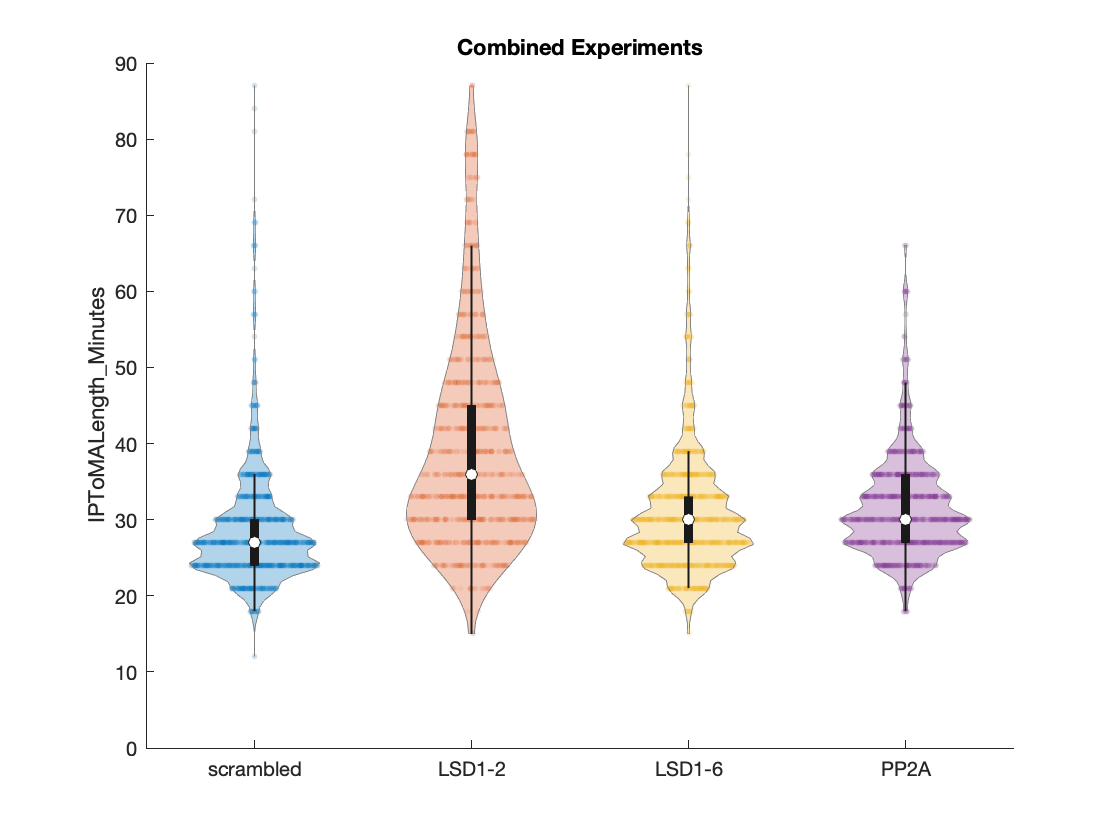

### LSD1_FusedProjects_CARSync_AdditionalFeatures_IPToMALength_Minutes_Histograms.png

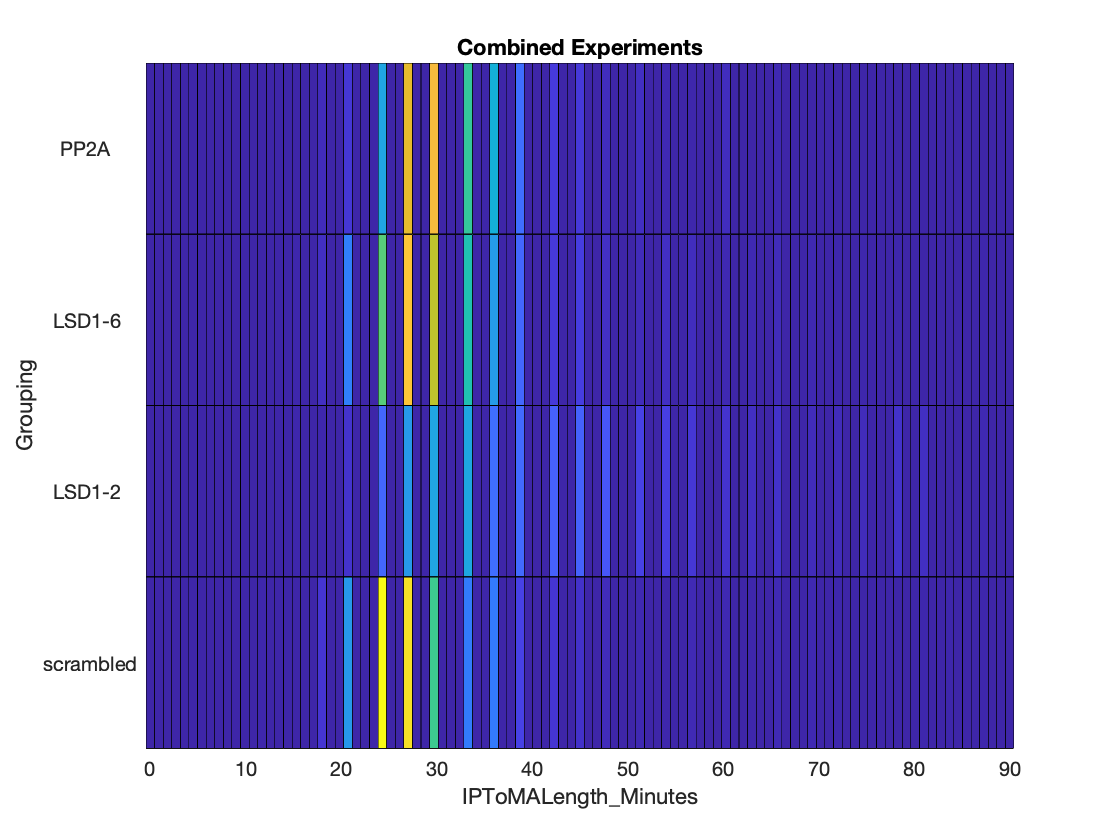

### LSD1_FusedProjects_CARSync_AdditionalFeatures_MajorAxisLength_HeatMaps.png

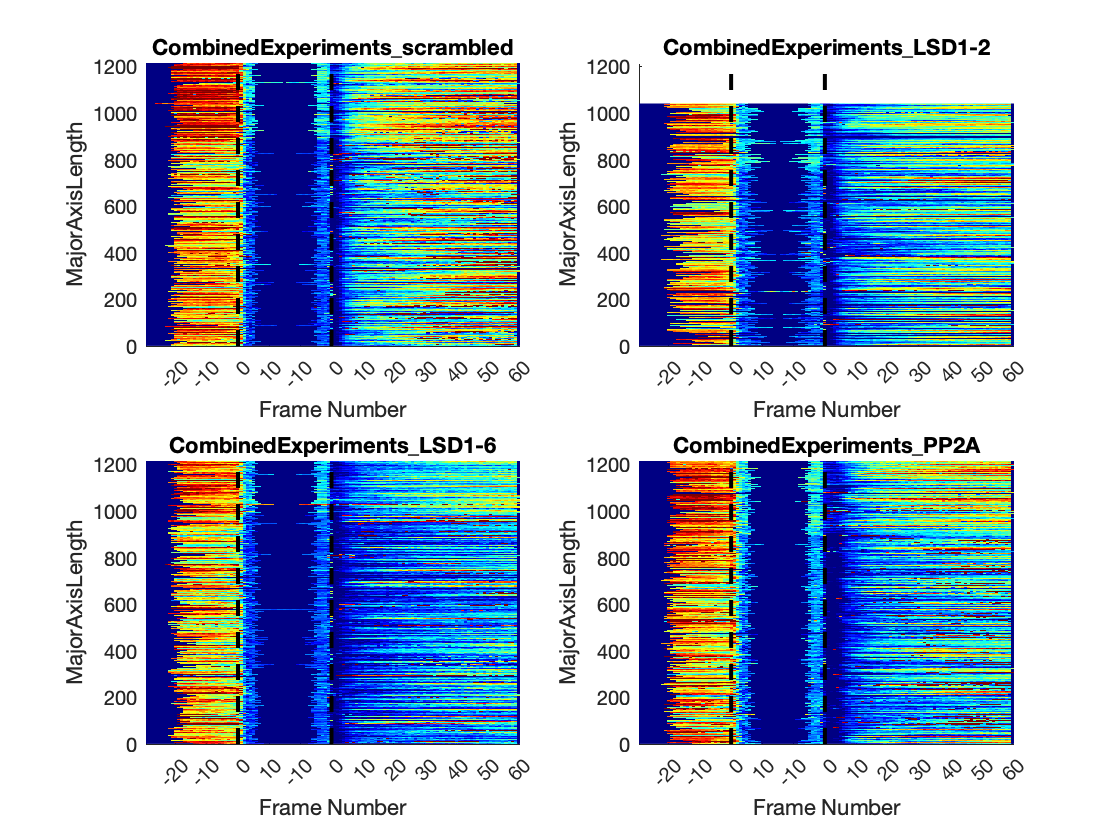

### LSD1_FusedProjects_CARSync_AdditionalFeatures_MajorAxisLength_LinePlots.png

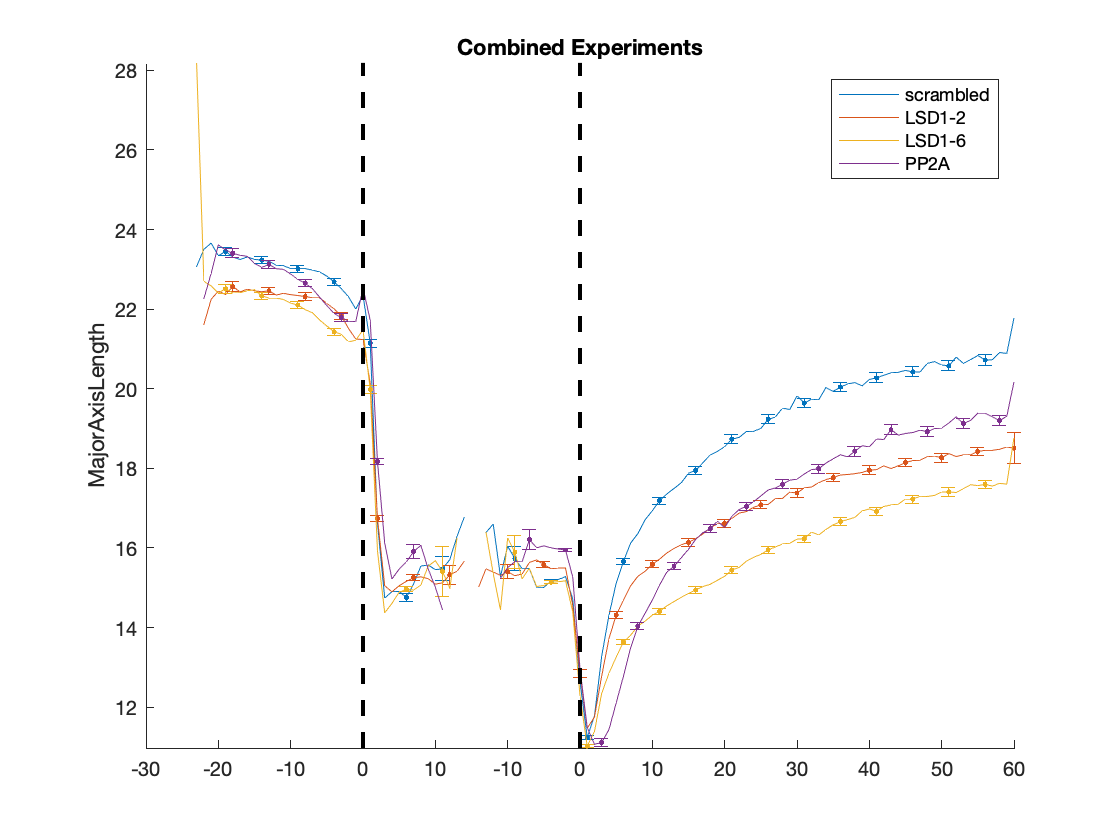

### LSD1_FusedProjects_CARSync_AdditionalFeatures_manuallyConfirmed_BoxPlots.png

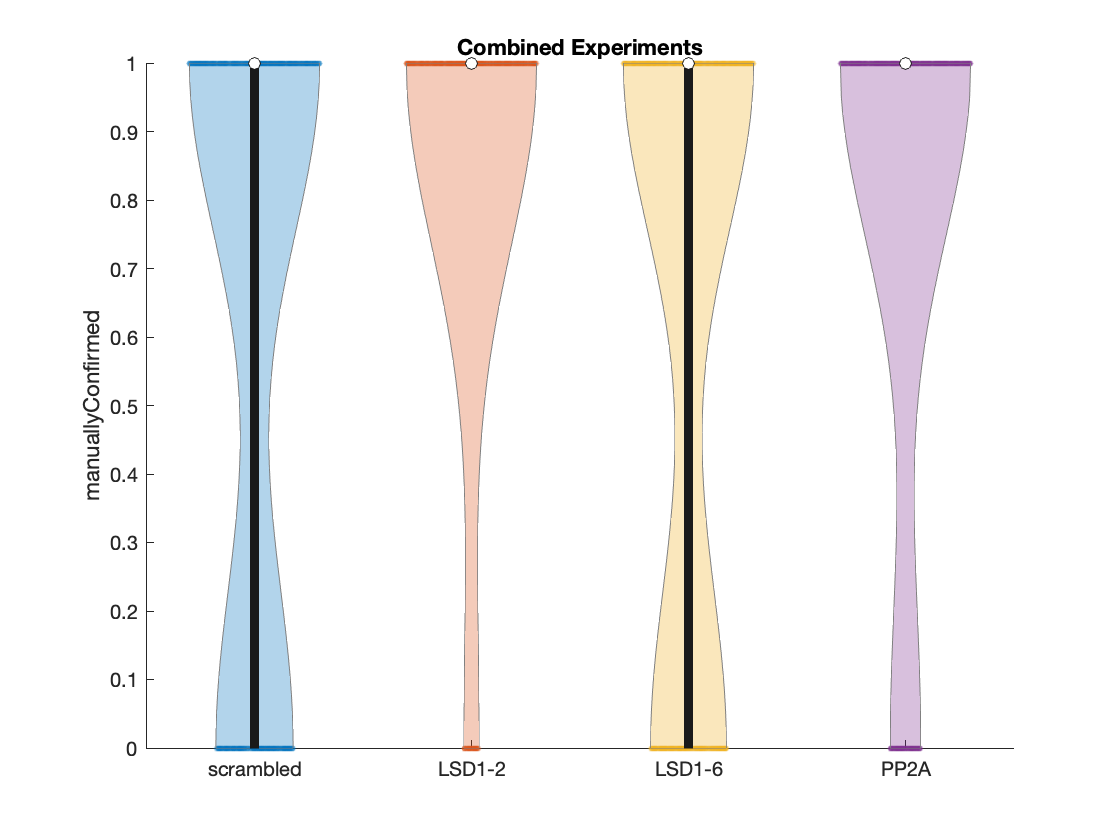

### LSD1_FusedProjects_CARSync_AdditionalFeatures_manuallyConfirmed_Histograms.png

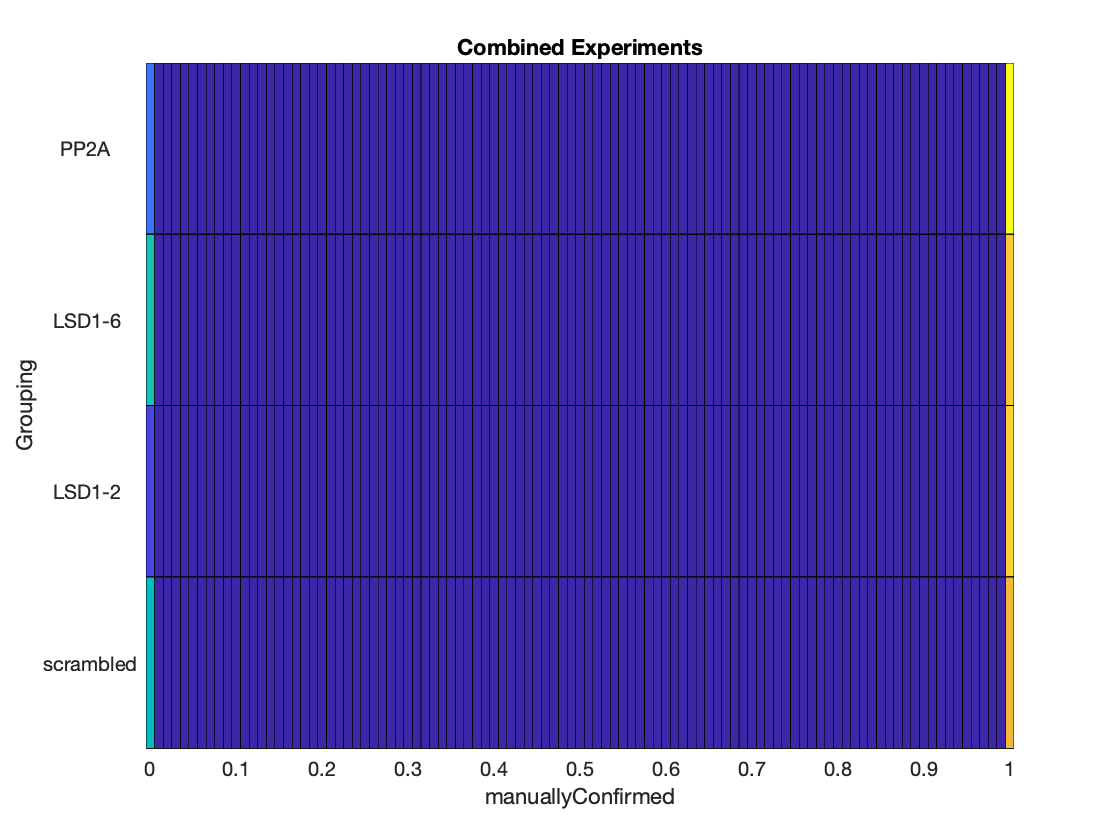

### LSD1_FusedProjects_CARSync_AdditionalFeatures_manualSynchronization_HeatMaps.png

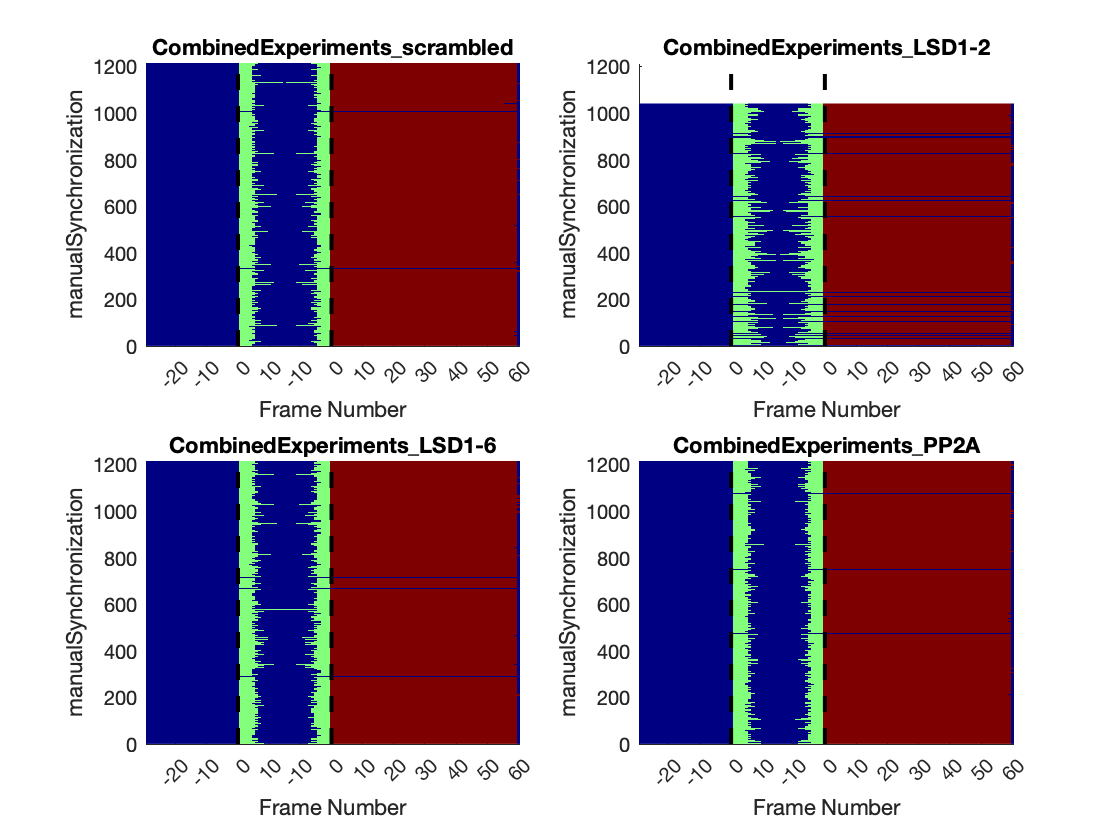

### LSD1_FusedProjects_CARSync_AdditionalFeatures_manualSynchronization_LinePlots.png

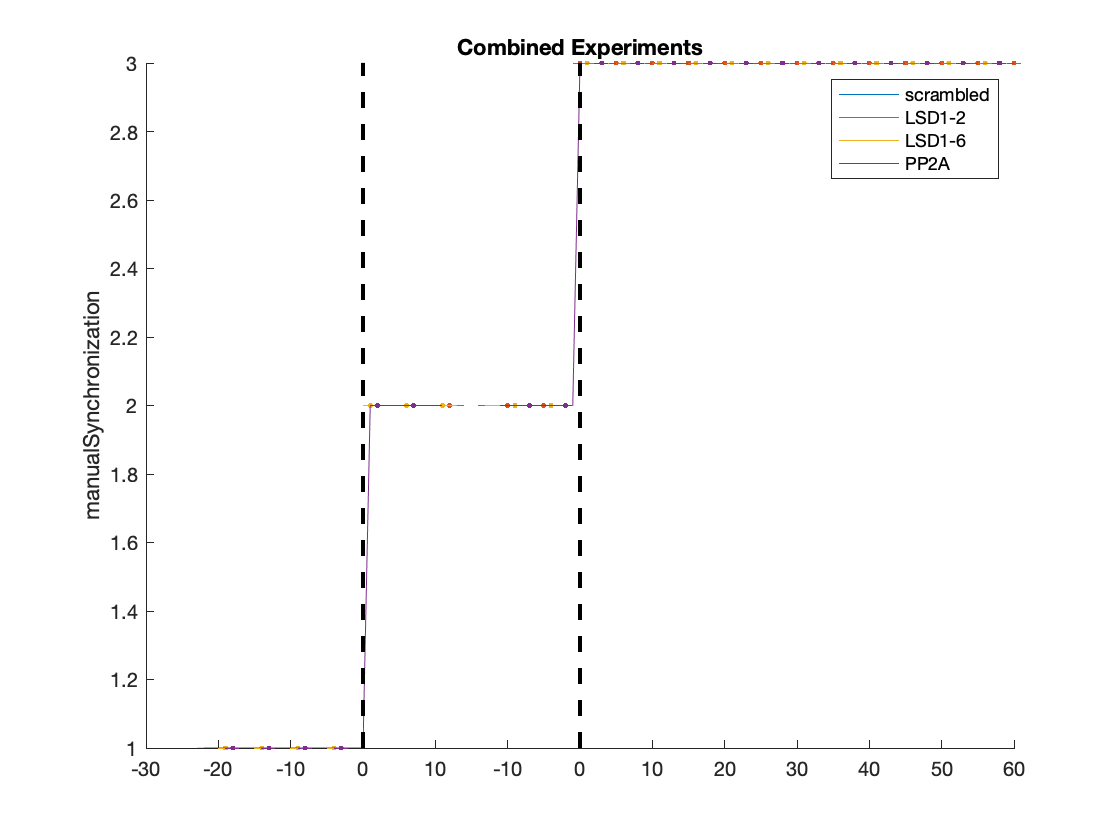

### LSD1_FusedProjects_CARSync_AdditionalFeatures_MaximalCorrelationCoefficient_HeatMaps.png

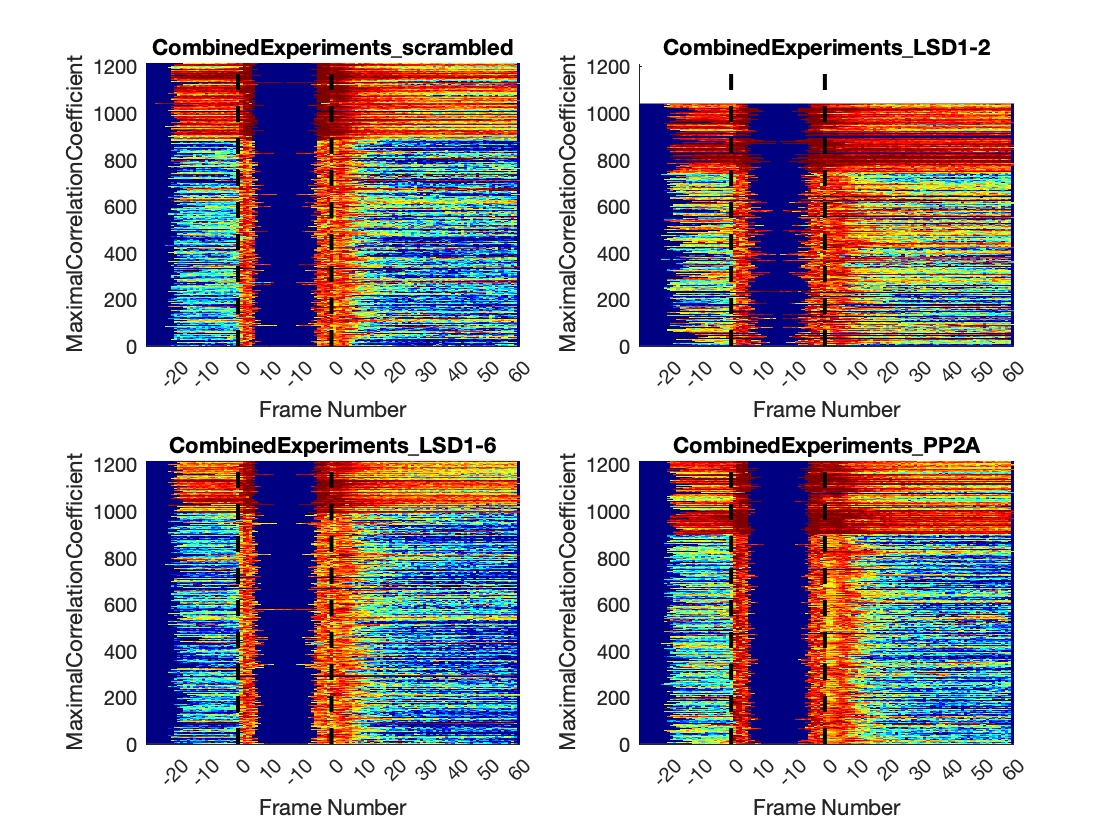

### LSD1_FusedProjects_CARSync_AdditionalFeatures_MaximalCorrelationCoefficient_LinePlots.png

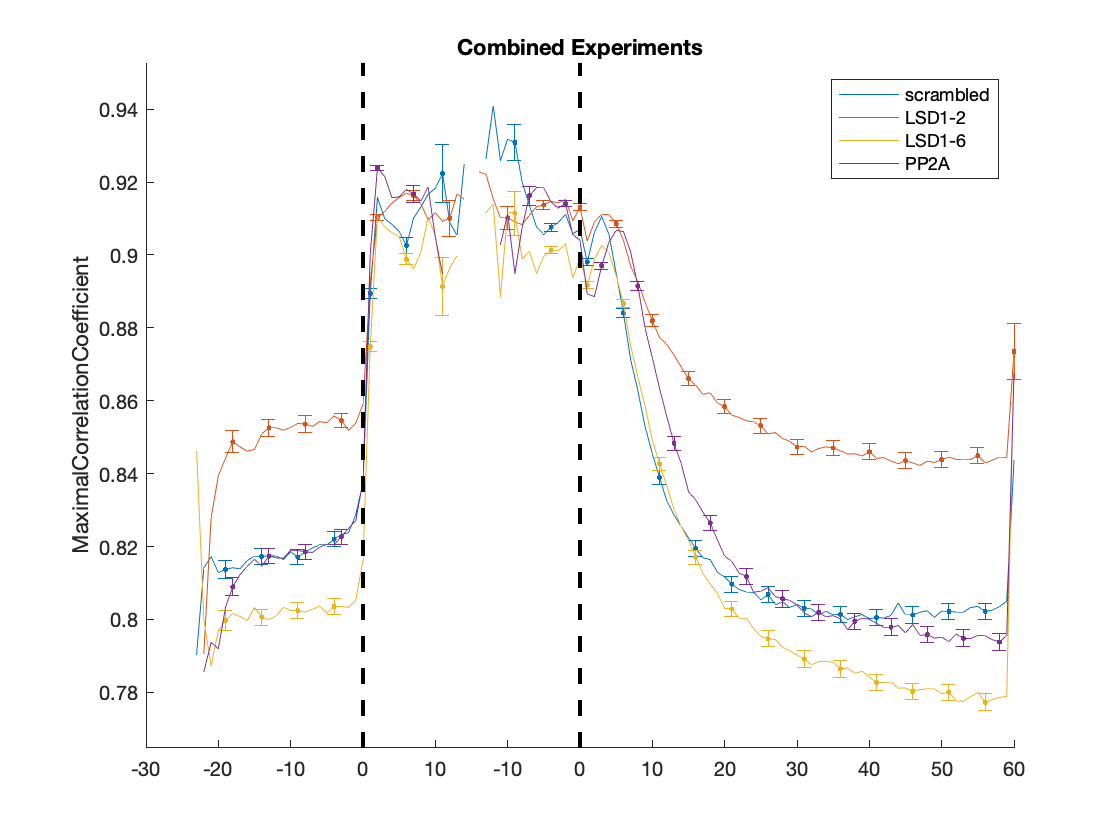

### LSD1_FusedProjects_CARSync_AdditionalFeatures_MeanIntensity-Normalized_HeatMaps.png

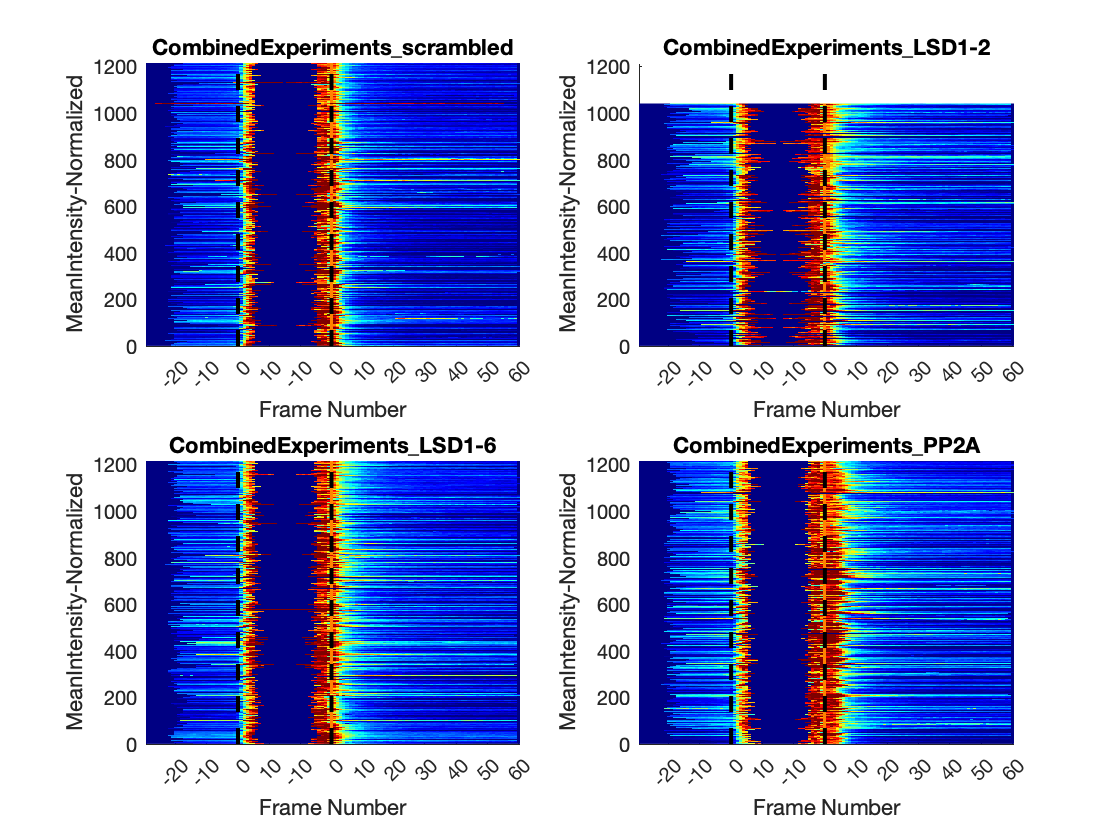

### LSD1_FusedProjects_CARSync_AdditionalFeatures_MeanIntensity-Normalized_LinePlots.png

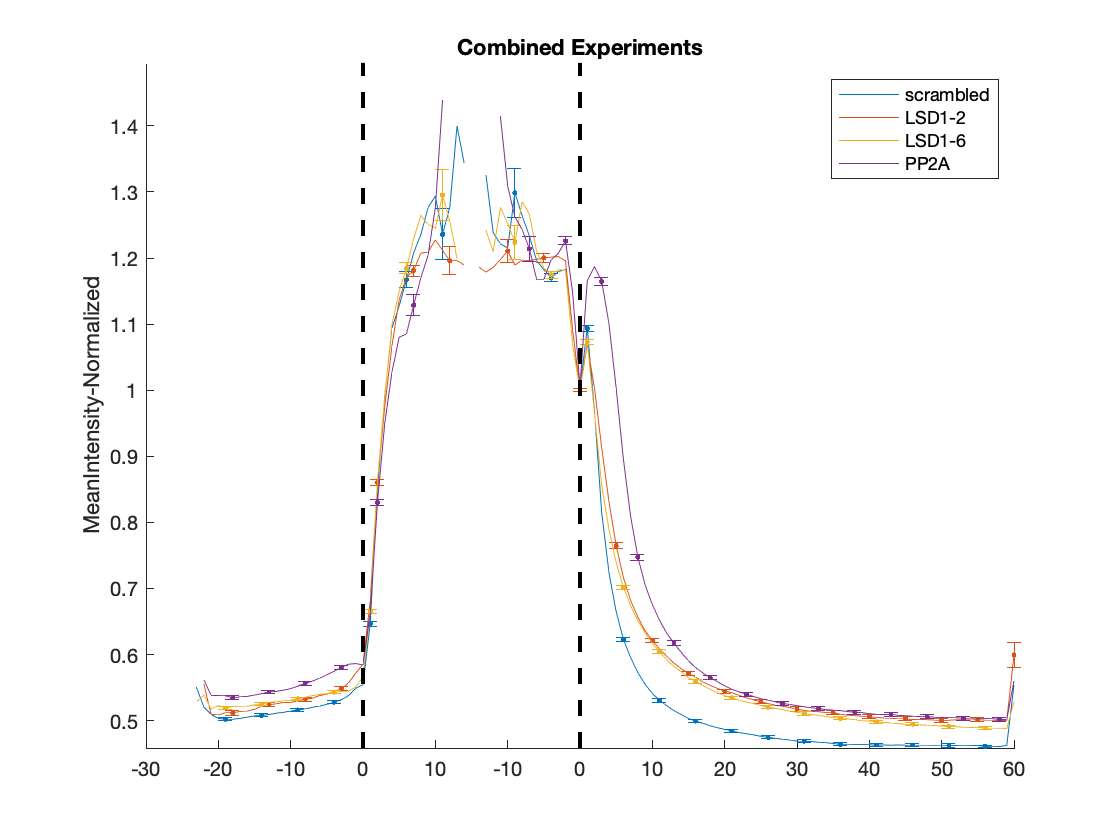

### LSD1_FusedProjects_CARSync_AdditionalFeatures_MeanIntensity-NormalizedIntMean_HeatMaps.png

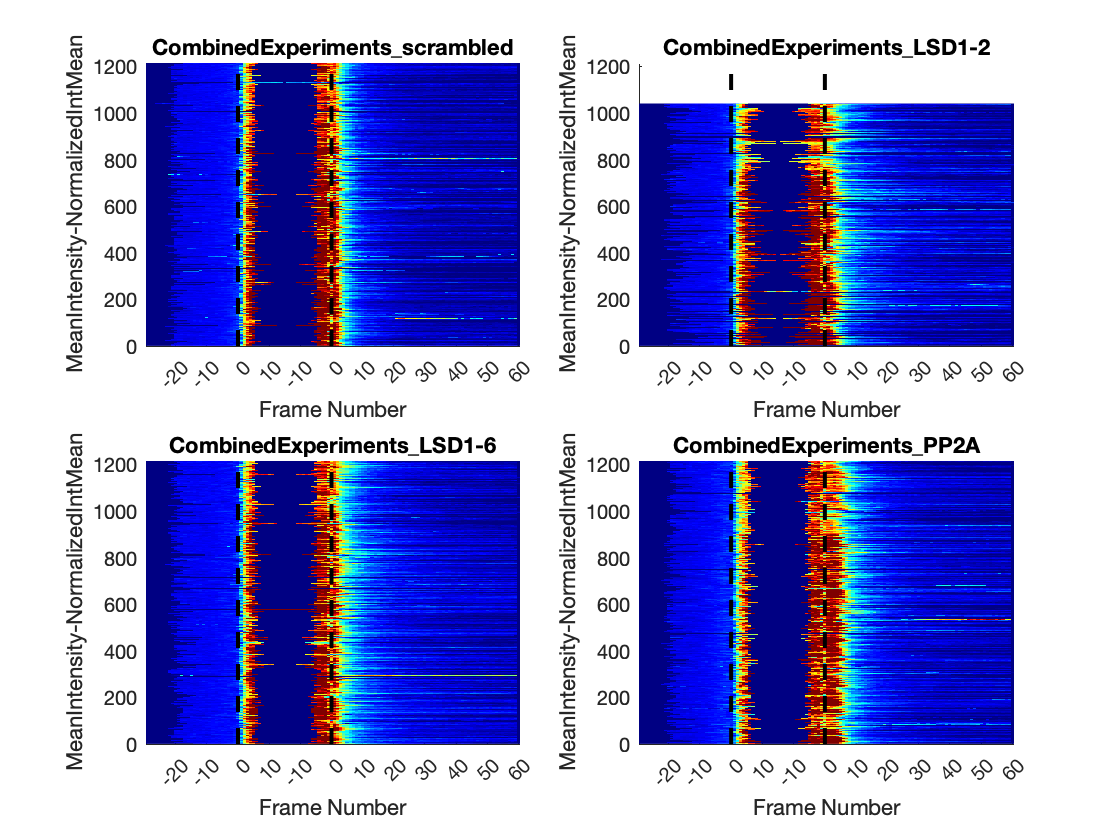

### LSD1_FusedProjects_CARSync_AdditionalFeatures_MeanIntensity-NormalizedIntMean_LinePlots.png

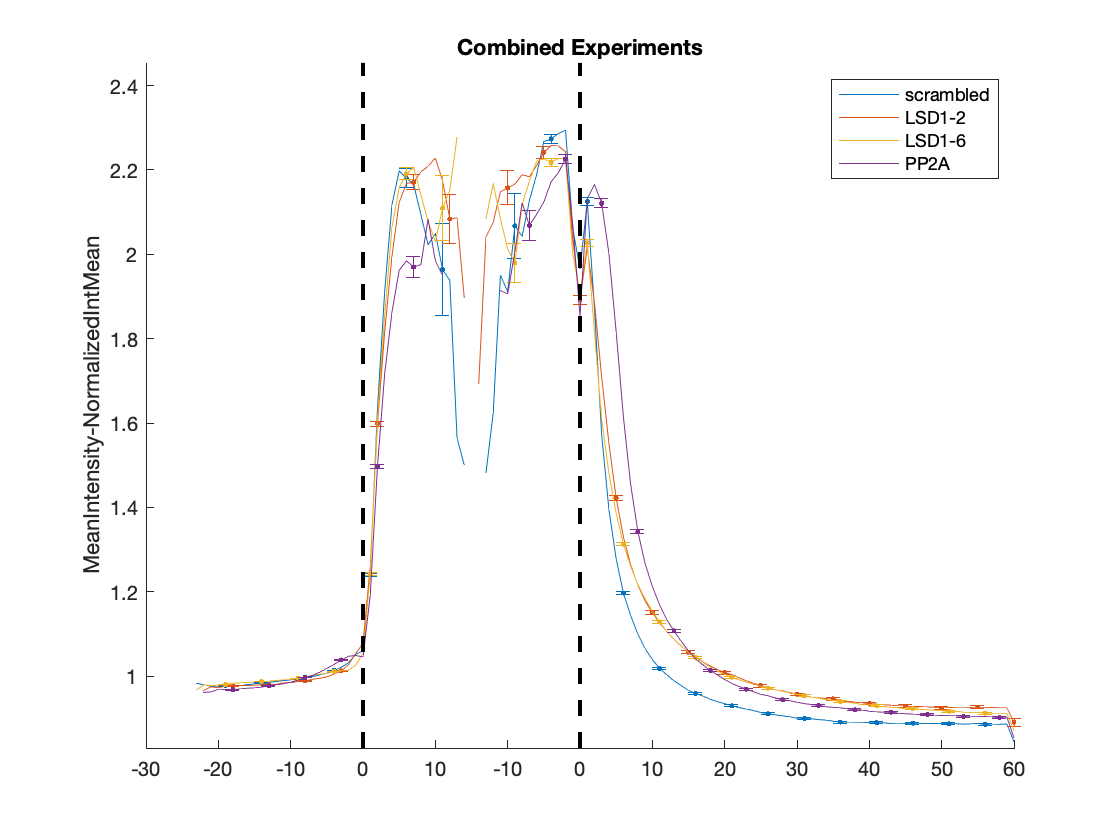

### LSD1_FusedProjects_CARSync_AdditionalFeatures_MeanIntensity_HeatMaps.png

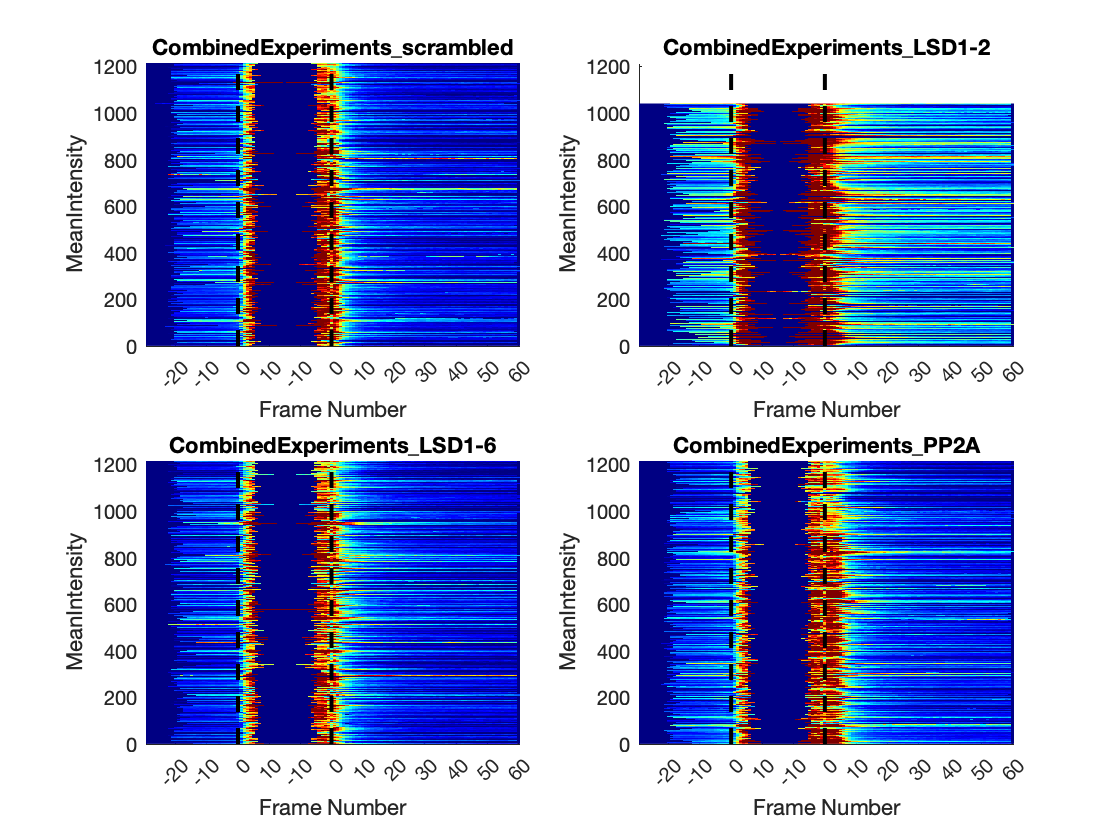

### LSD1_FusedProjects_CARSync_AdditionalFeatures_MeanIntensity_LinePlots.png

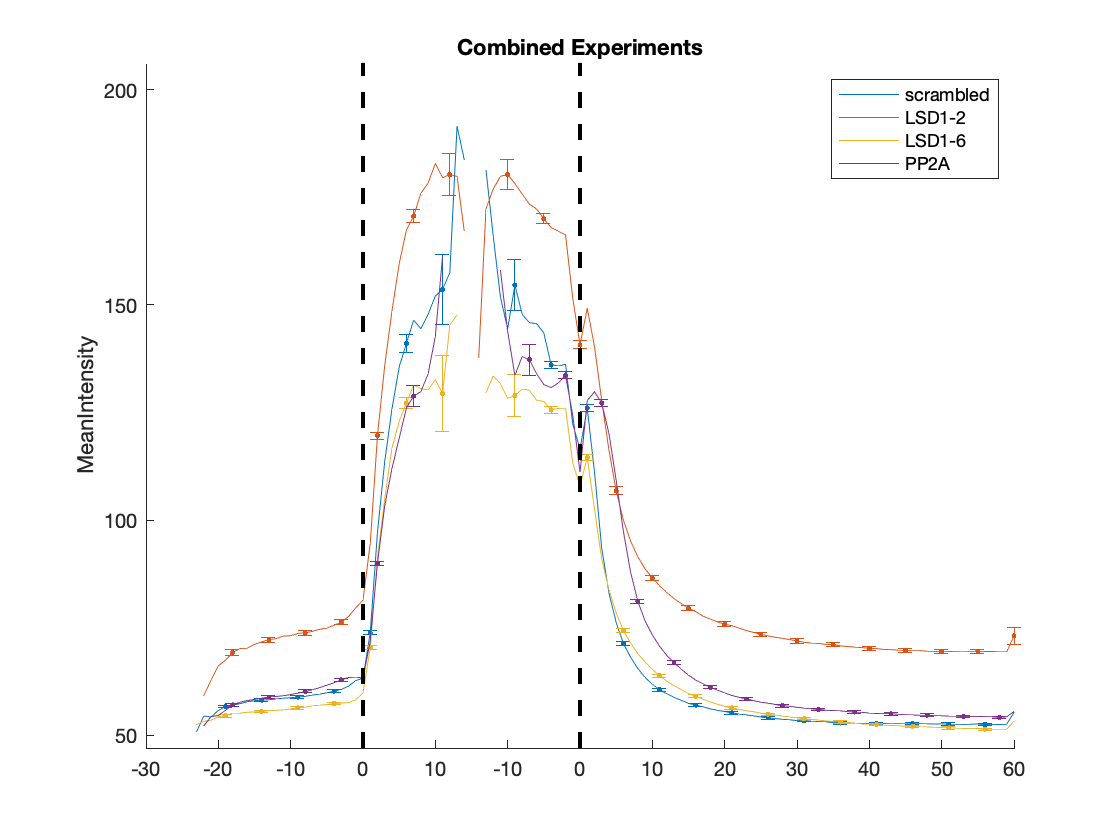

### LSD1_FusedProjects_CARSync_AdditionalFeatures_MinMajAxisRatio_HeatMaps.png

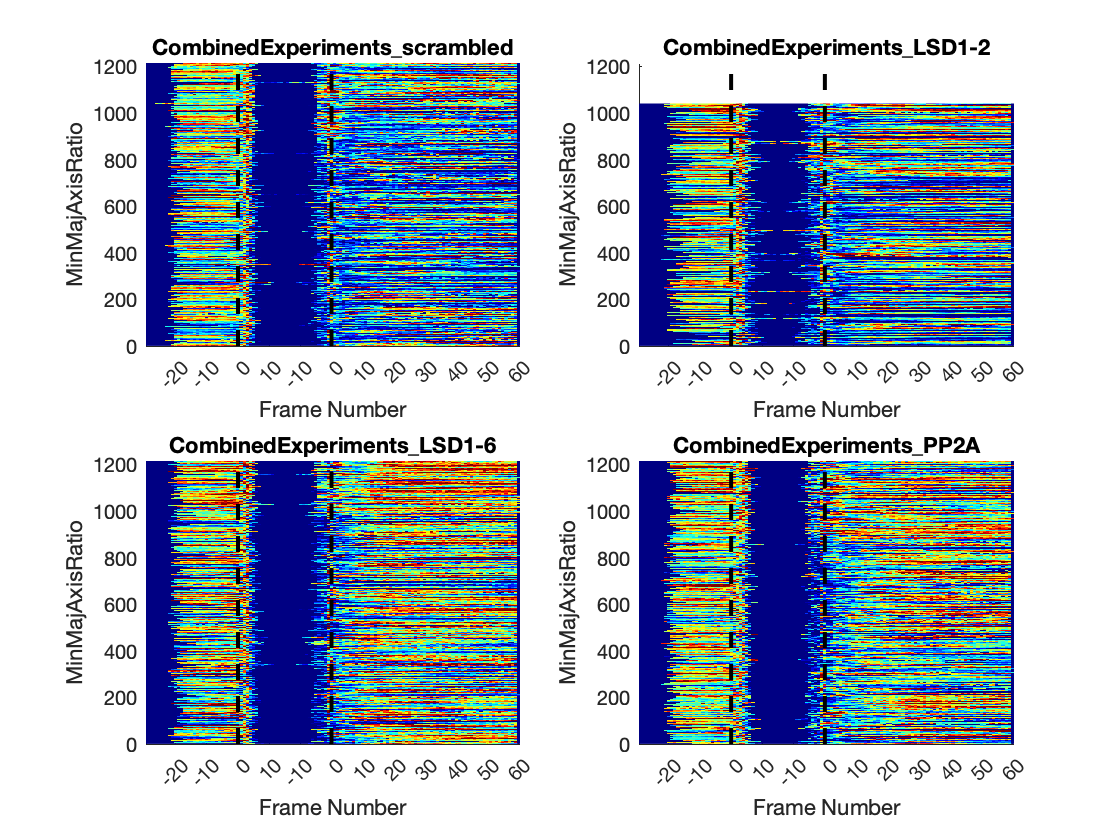

### LSD1_FusedProjects_CARSync_AdditionalFeatures_MinMajAxisRatio_LinePlots.png

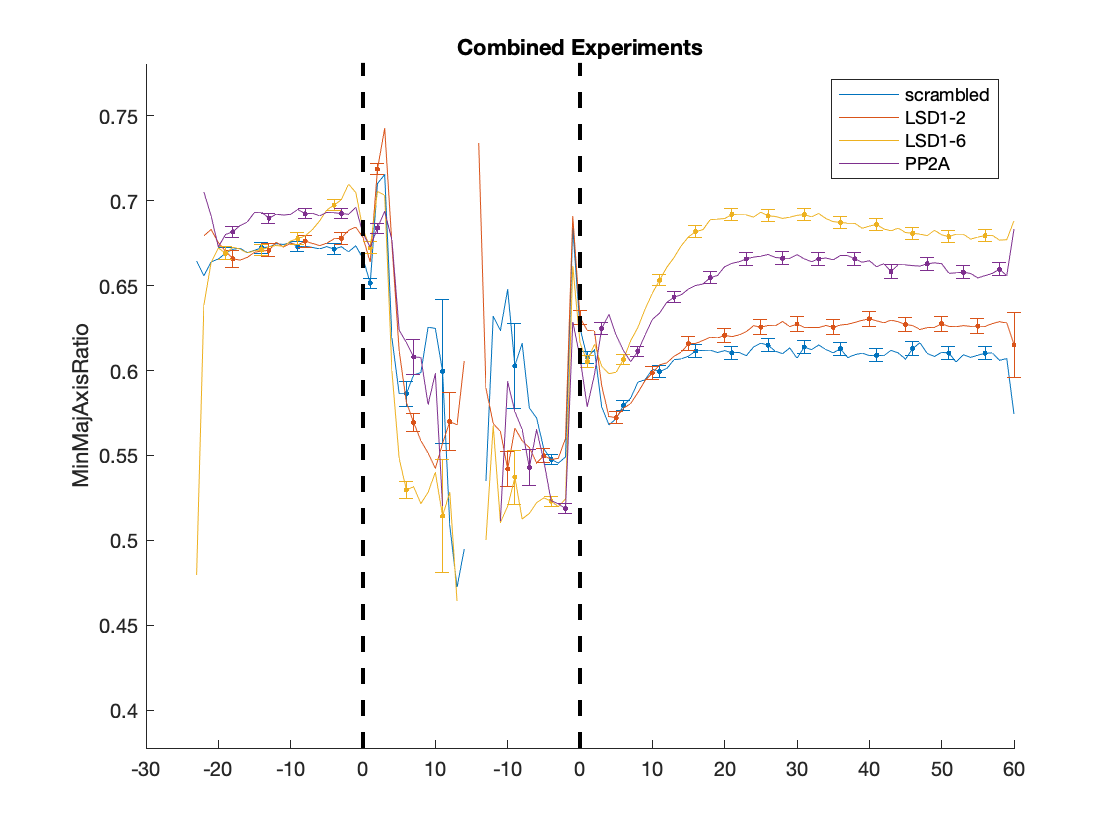

### LSD1_FusedProjects_CARSync_AdditionalFeatures_MinorAxisLength_HeatMaps.png

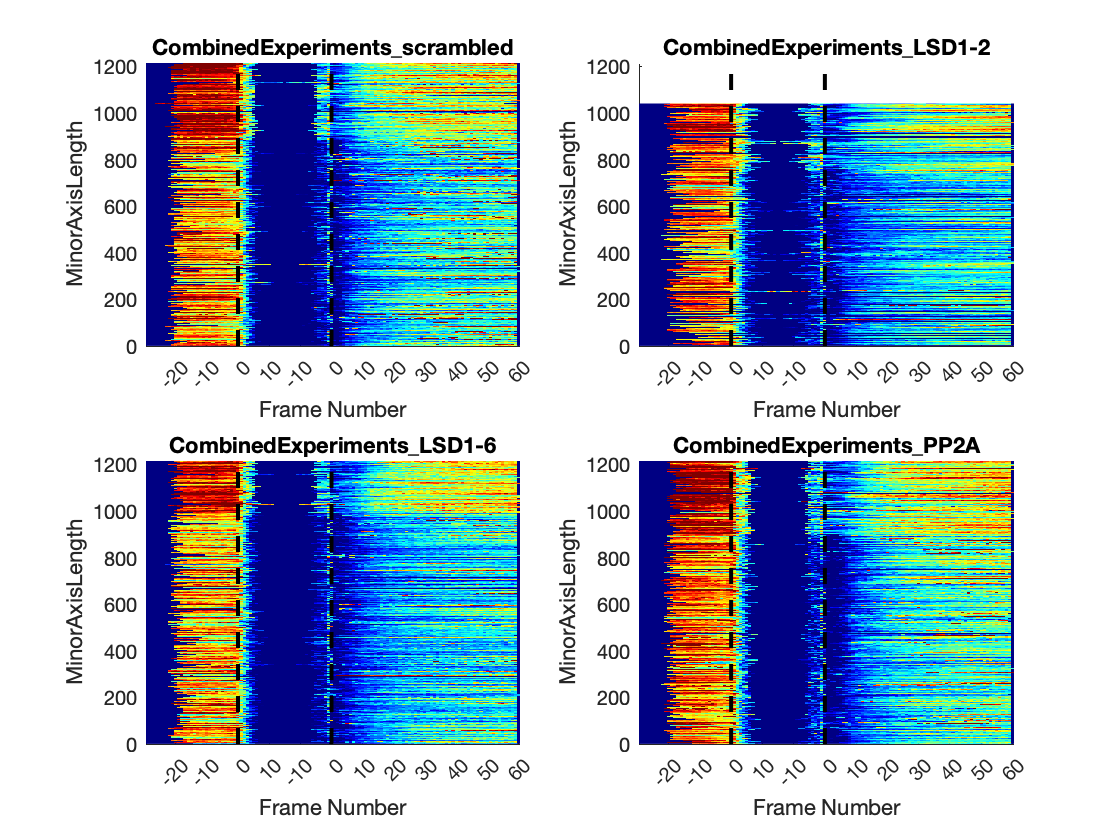

### LSD1_FusedProjects_CARSync_AdditionalFeatures_MinorAxisLength_LinePlots.png

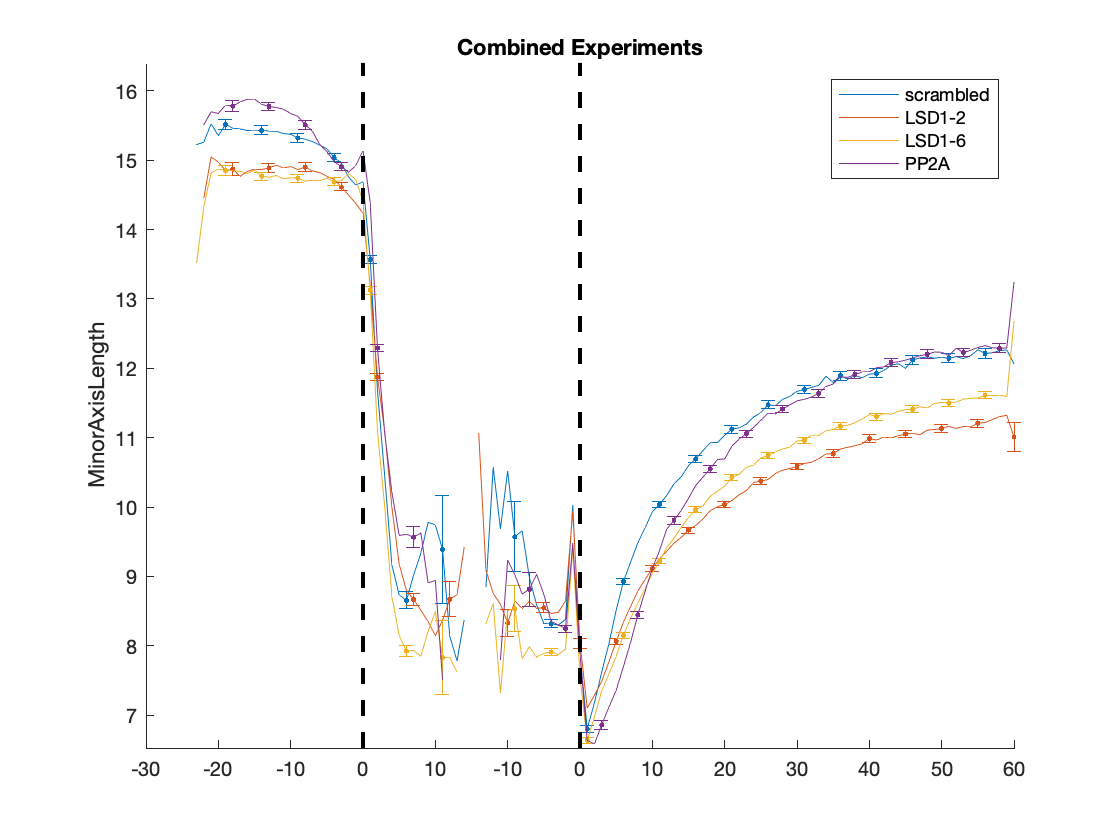

### LSD1_FusedProjects_CARSync_AdditionalFeatures_Orientation_HeatMaps.png

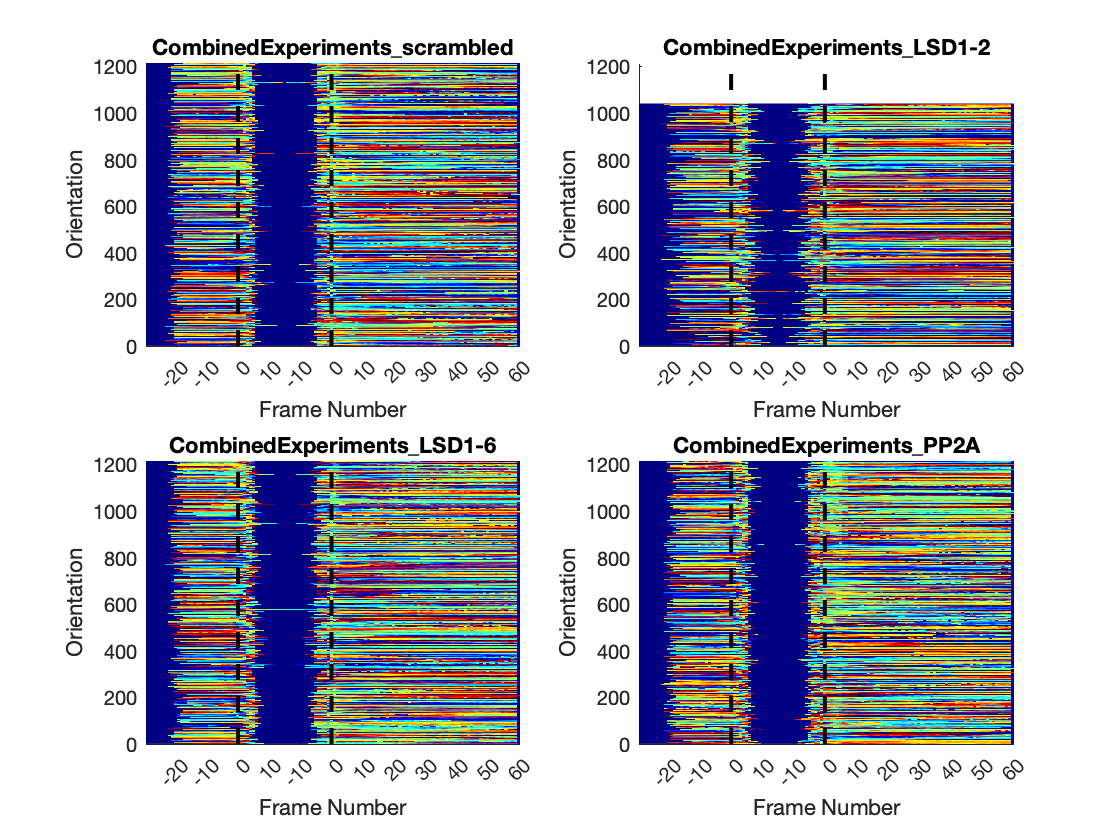

### LSD1_FusedProjects_CARSync_AdditionalFeatures_Orientation_LinePlots.png

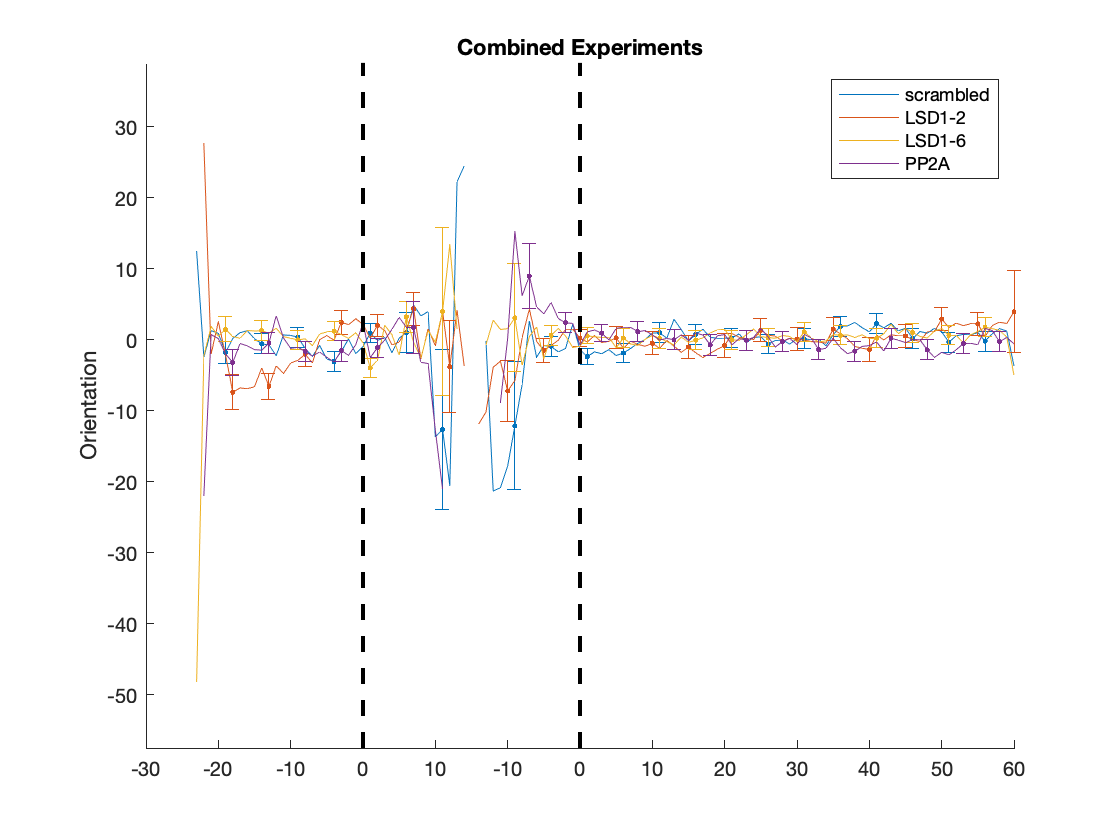

### LSD1_FusedProjects_CARSync_AdditionalFeatures_Ratio-MajorAxisLength-vs-MinorAxisLength_HeatMaps.png

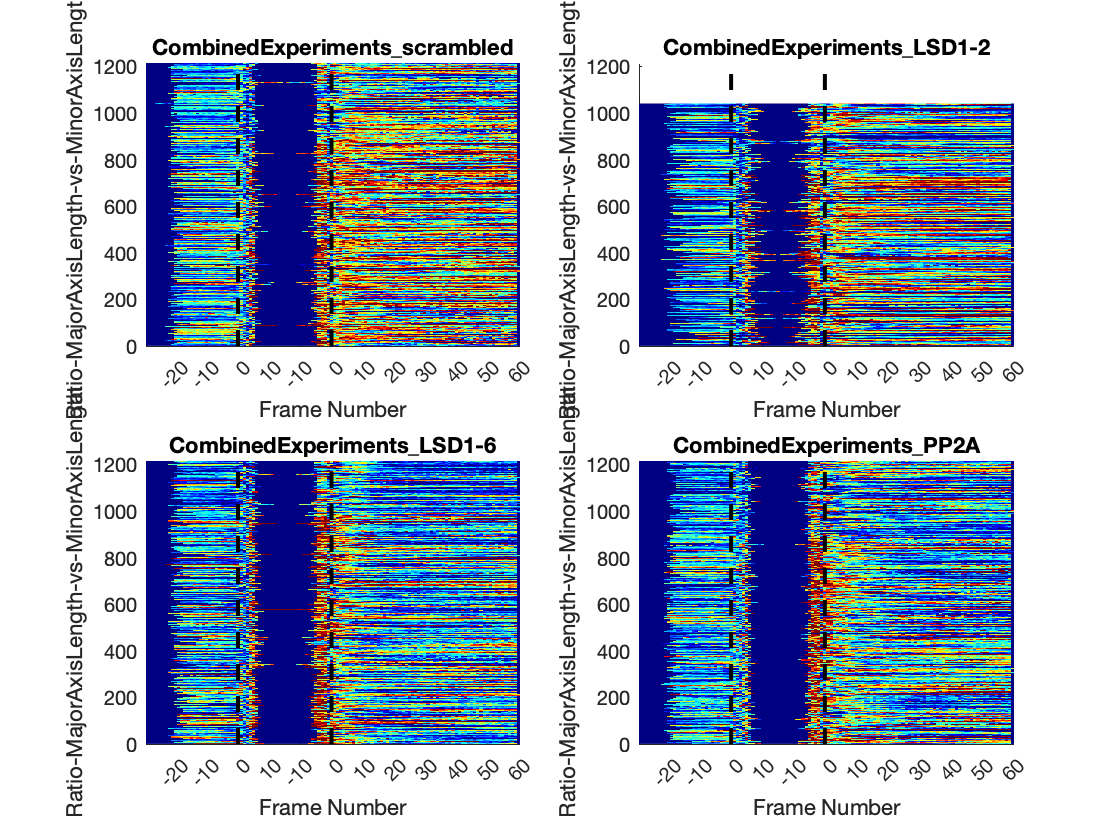

### LSD1_FusedProjects_CARSync_AdditionalFeatures_Ratio-MajorAxisLength-vs-MinorAxisLength_LinePlots.png

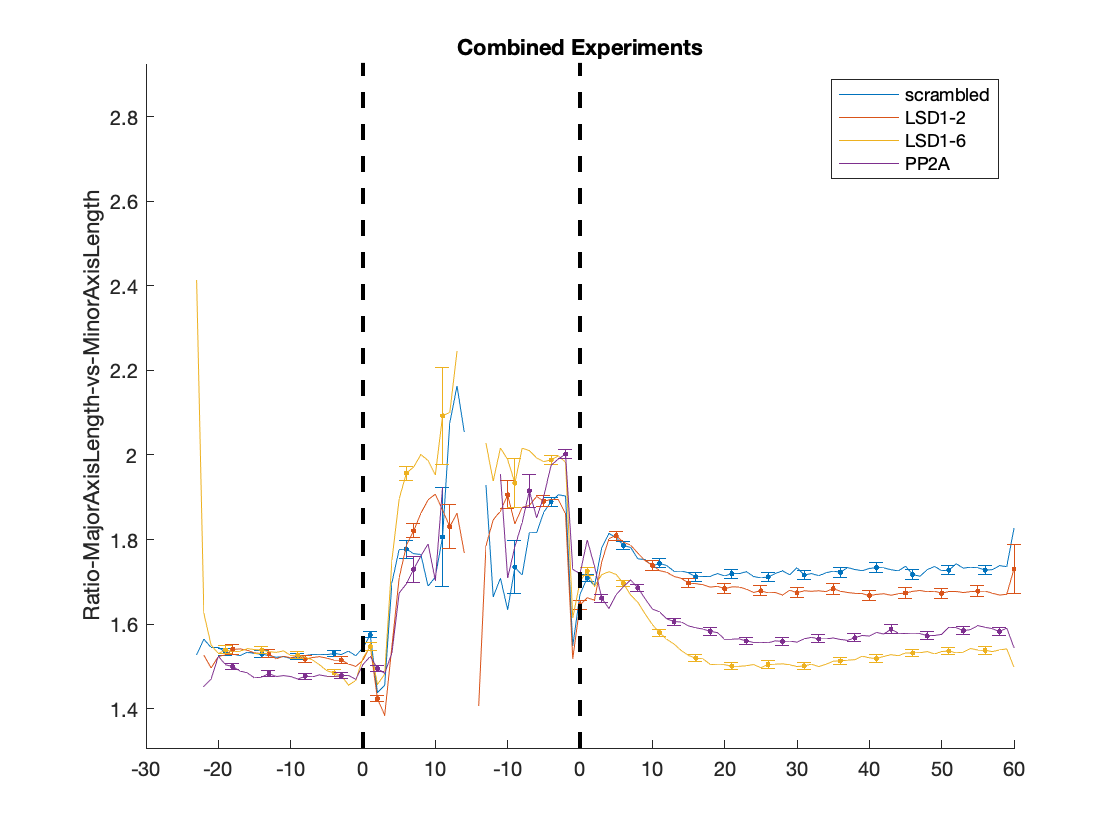

### LSD1_FusedProjects_CARSync_AdditionalFeatures_Ratio-MinorAxisLength-vs-MajorAxisLength_HeatMaps.png

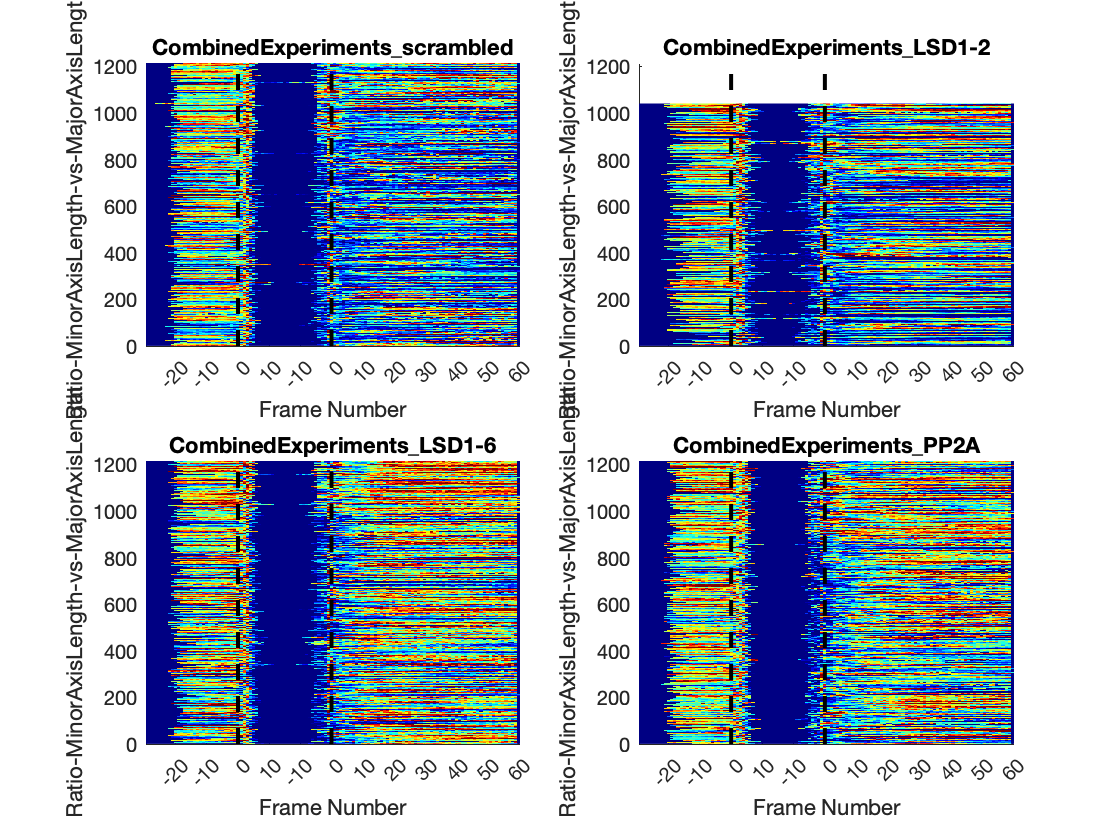

### LSD1_FusedProjects_CARSync_AdditionalFeatures_Ratio-MinorAxisLength-vs-MajorAxisLength_LinePlots.png

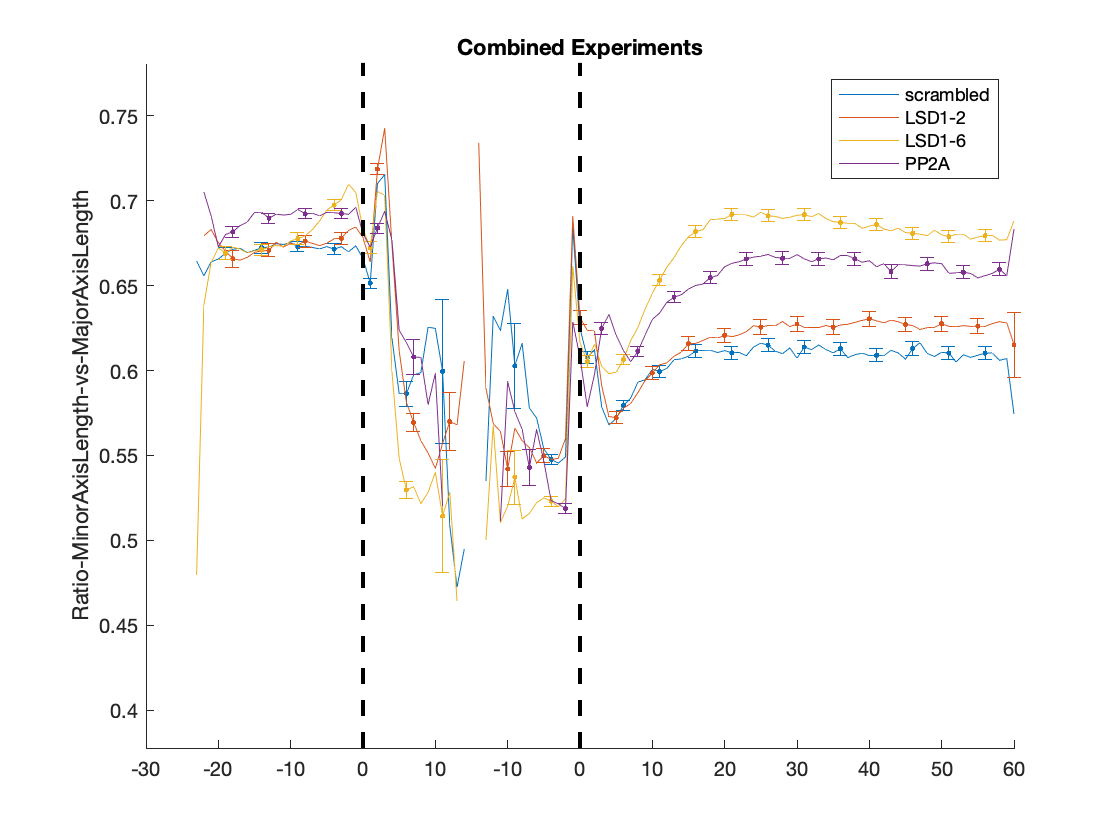

### LSD1_FusedProjects_CARSync_AdditionalFeatures_scale_HeatMaps.png

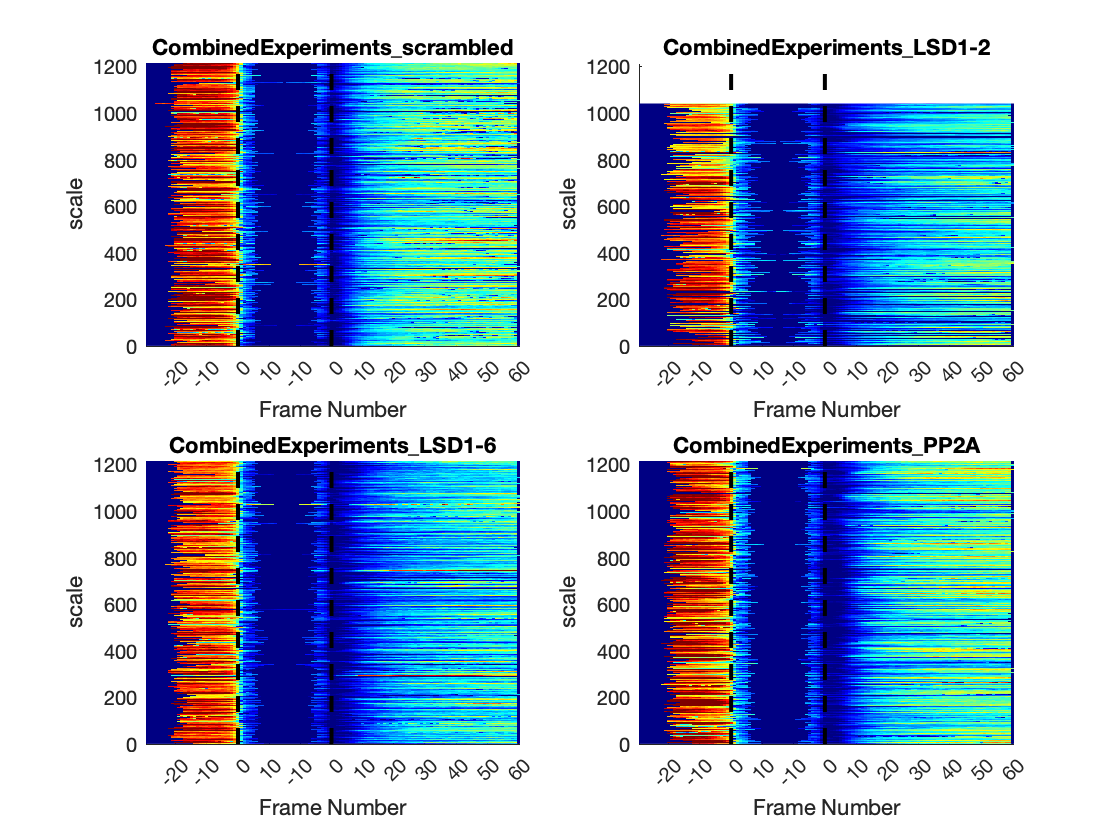

### LSD1_FusedProjects_CARSync_AdditionalFeatures_scale_LinePlots.png

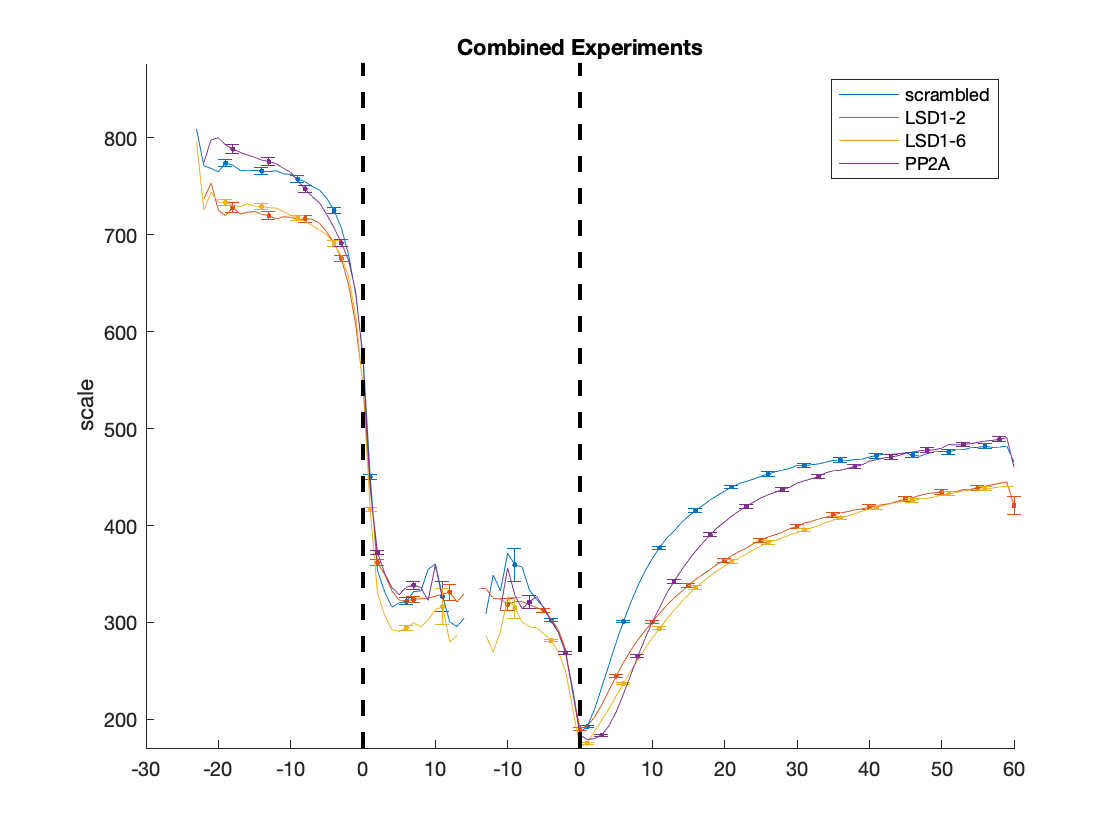

### LSD1_FusedProjects_CARSync_AdditionalFeatures_StdIntensity-Normalized_HeatMaps.png

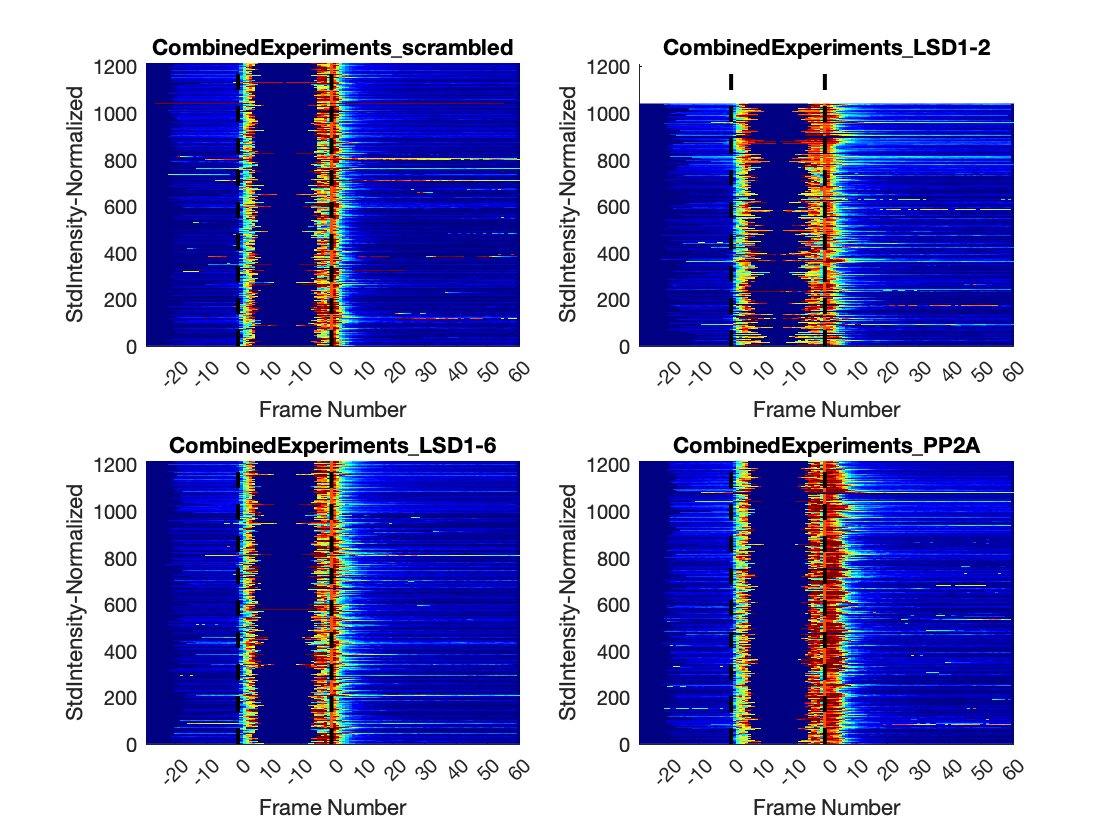

### LSD1_FusedProjects_CARSync_AdditionalFeatures_StdIntensity-Normalized_LinePlots.png

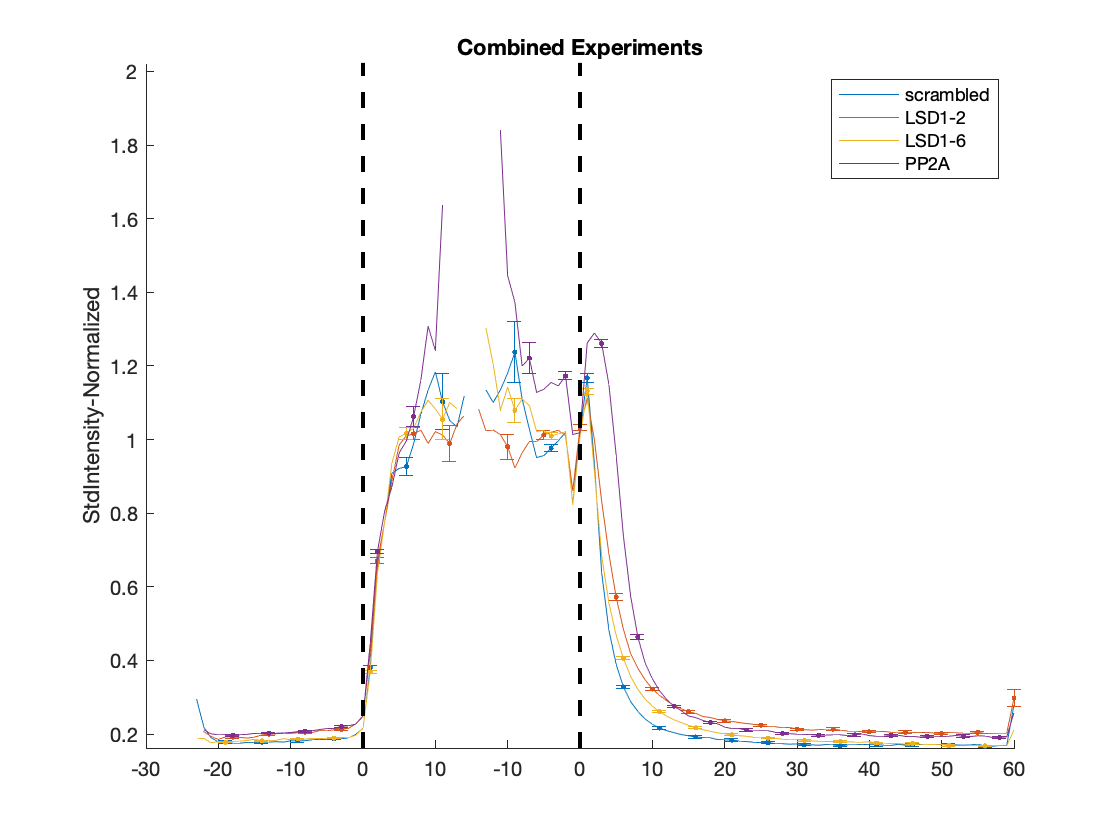

### LSD1_FusedProjects_CARSync_AdditionalFeatures_StdIntensity_HeatMaps.png

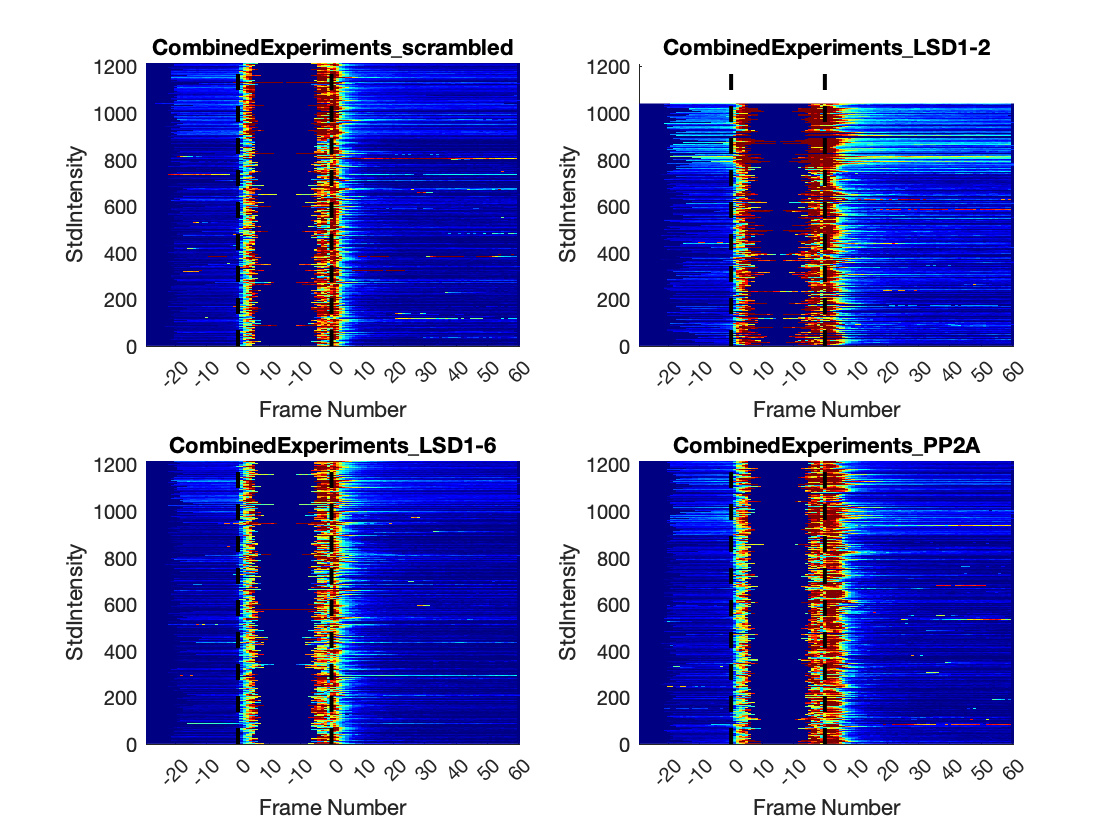

### LSD1_FusedProjects_CARSync_AdditionalFeatures_StdIntensity_LinePlots.png

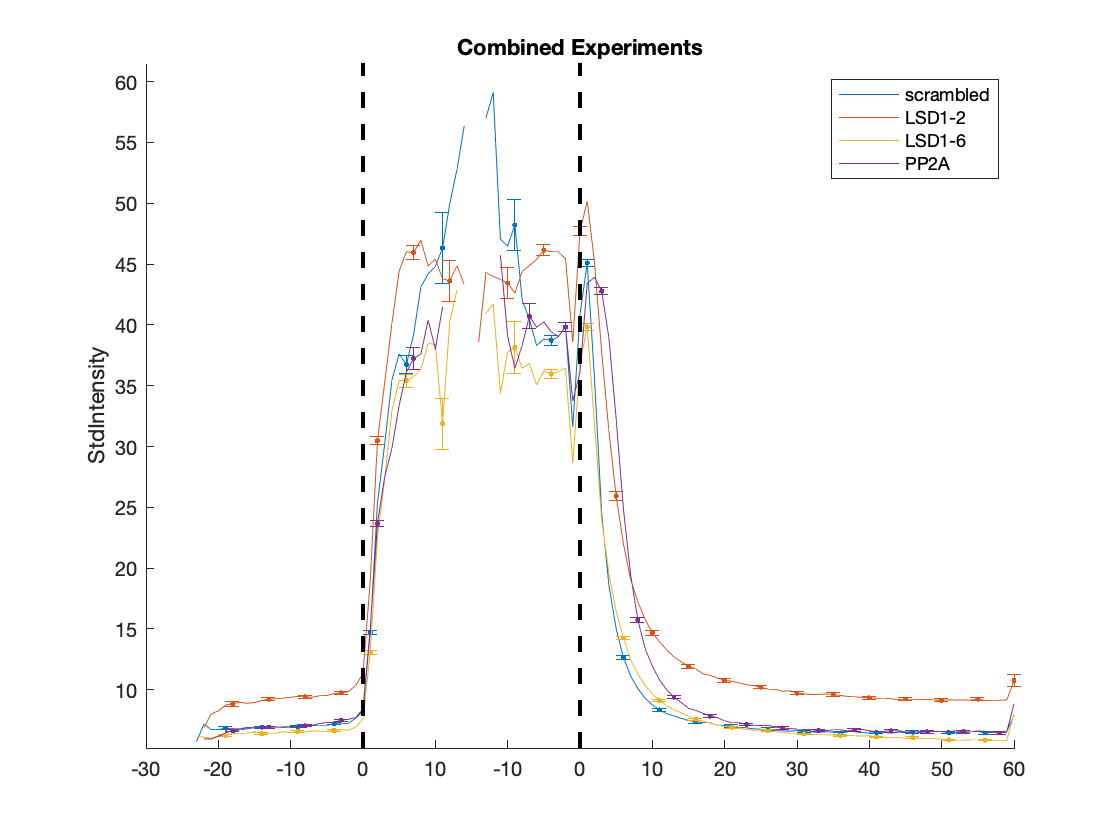

### LSD1_FusedProjects_CARSync_AdditionalFeatures_StdIntensityGradMag_HeatMaps.png

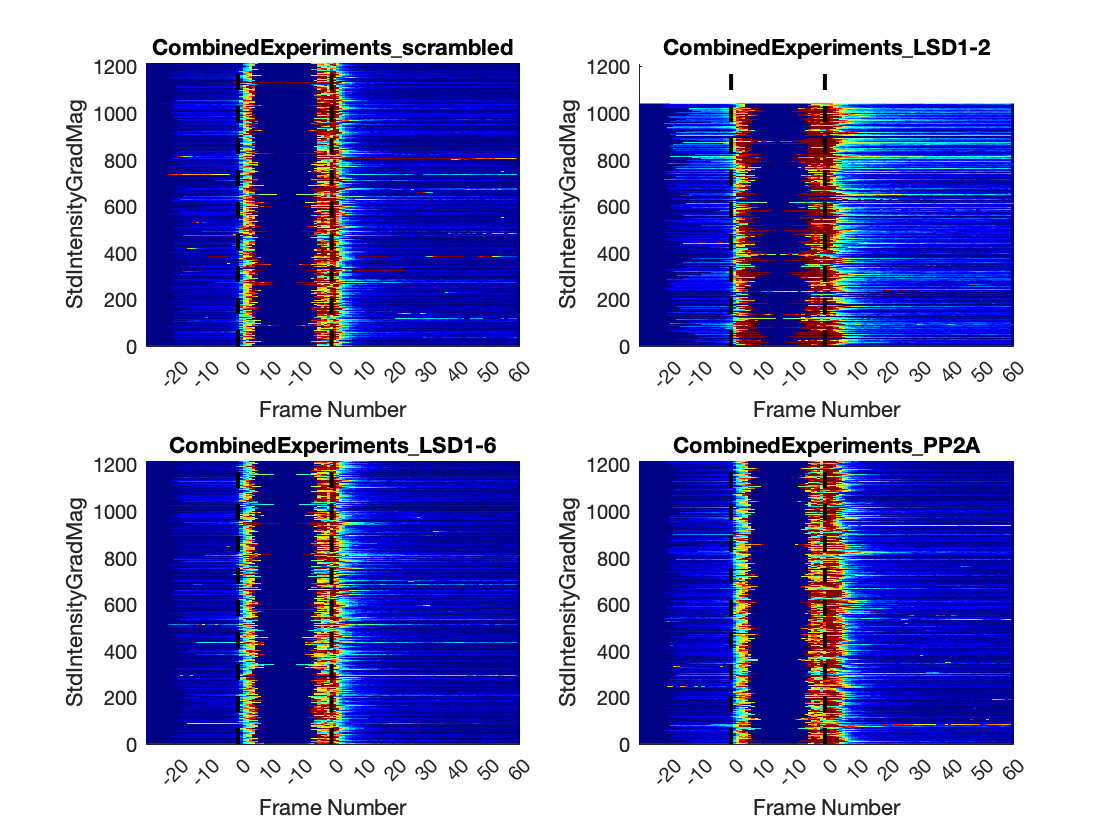

### LSD1_FusedProjects_CARSync_AdditionalFeatures_StdIntensityGradMag_LinePlots.png

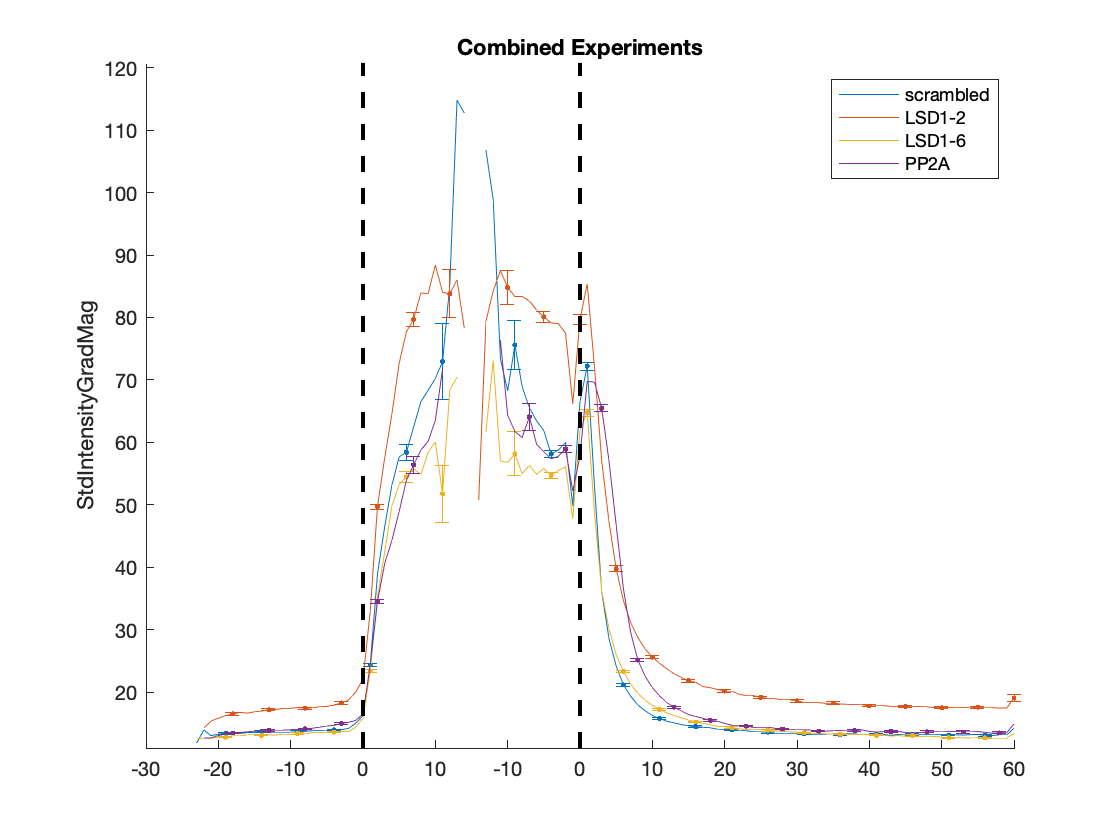

### LSD1_FusedProjects_CARSync_AdditionalFeatures_SumAverage_HeatMaps.png

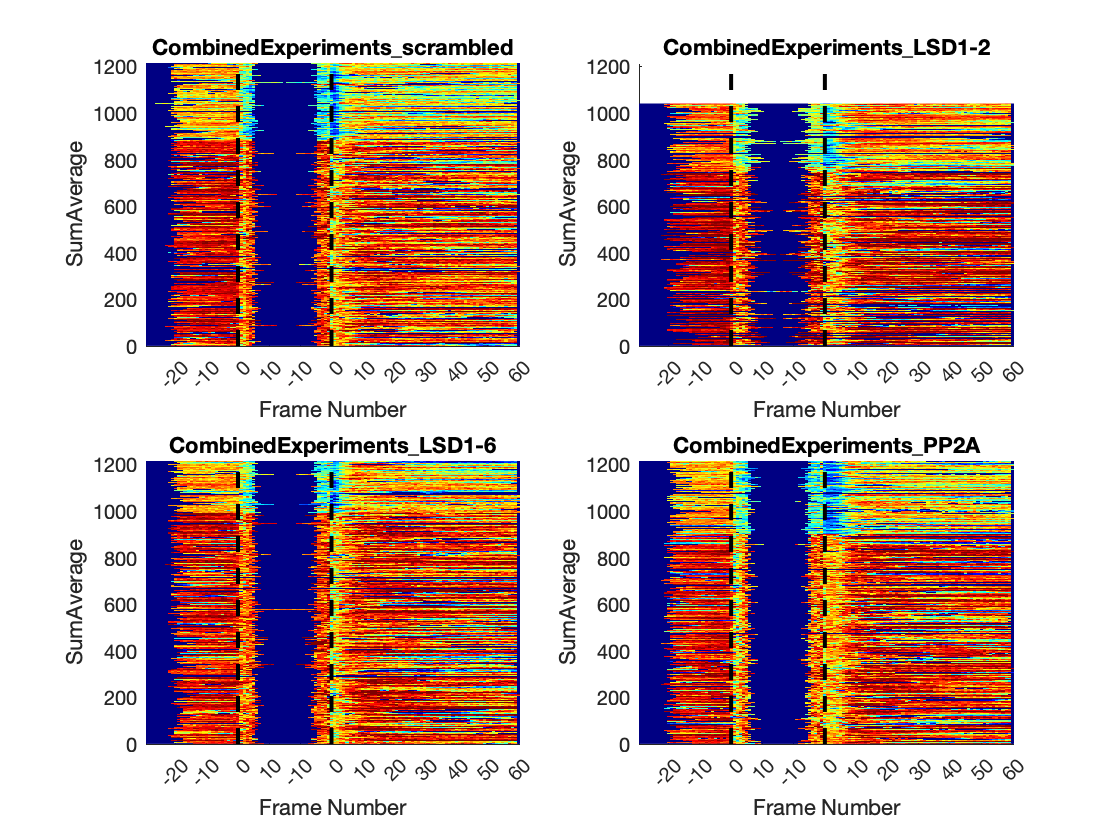

### LSD1_FusedProjects_CARSync_AdditionalFeatures_SumAverage_LinePlots.png

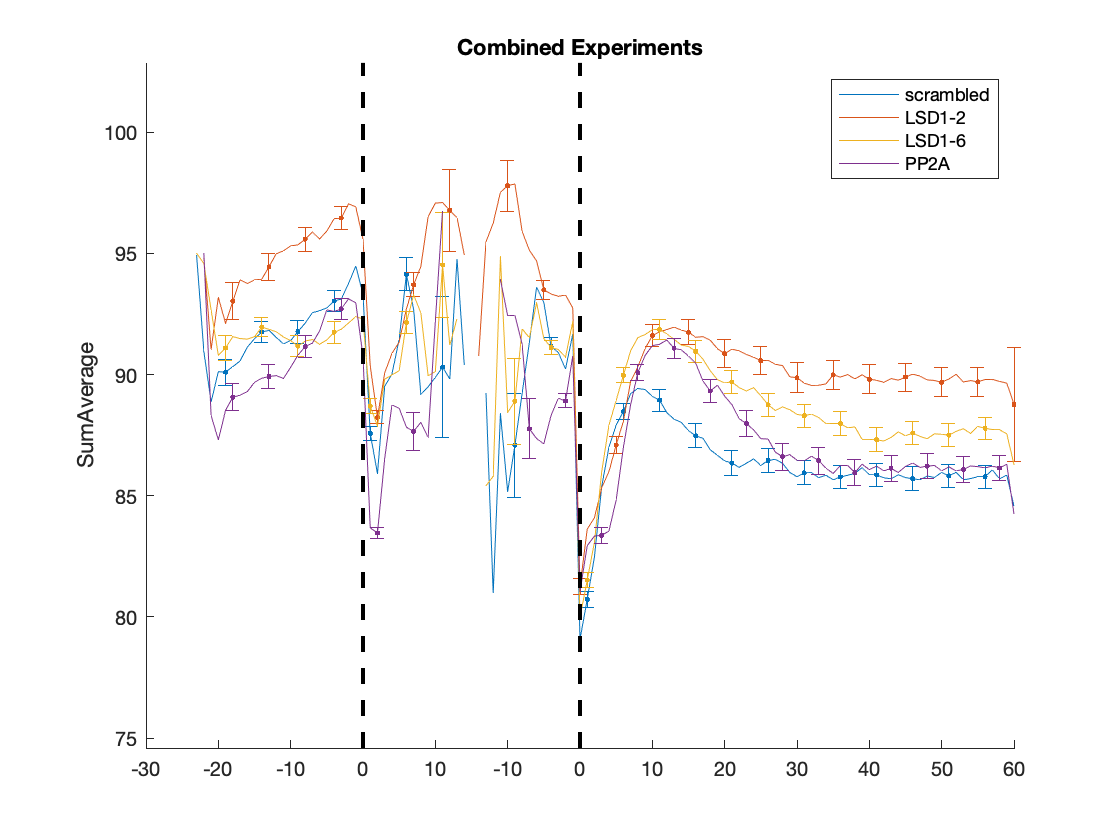

### LSD1_FusedProjects_CARSync_AdditionalFeatures_SumEntropy_HeatMaps.png

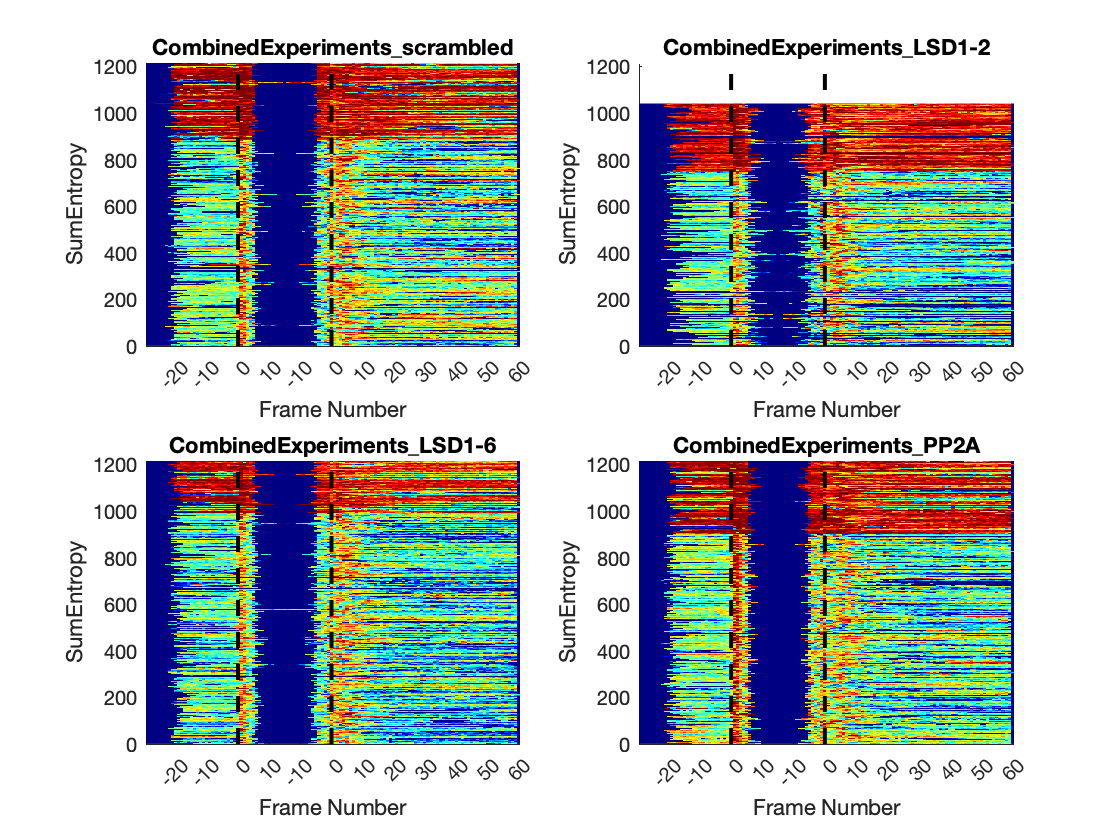

### LSD1_FusedProjects_CARSync_AdditionalFeatures_SumEntropy_LinePlots.png

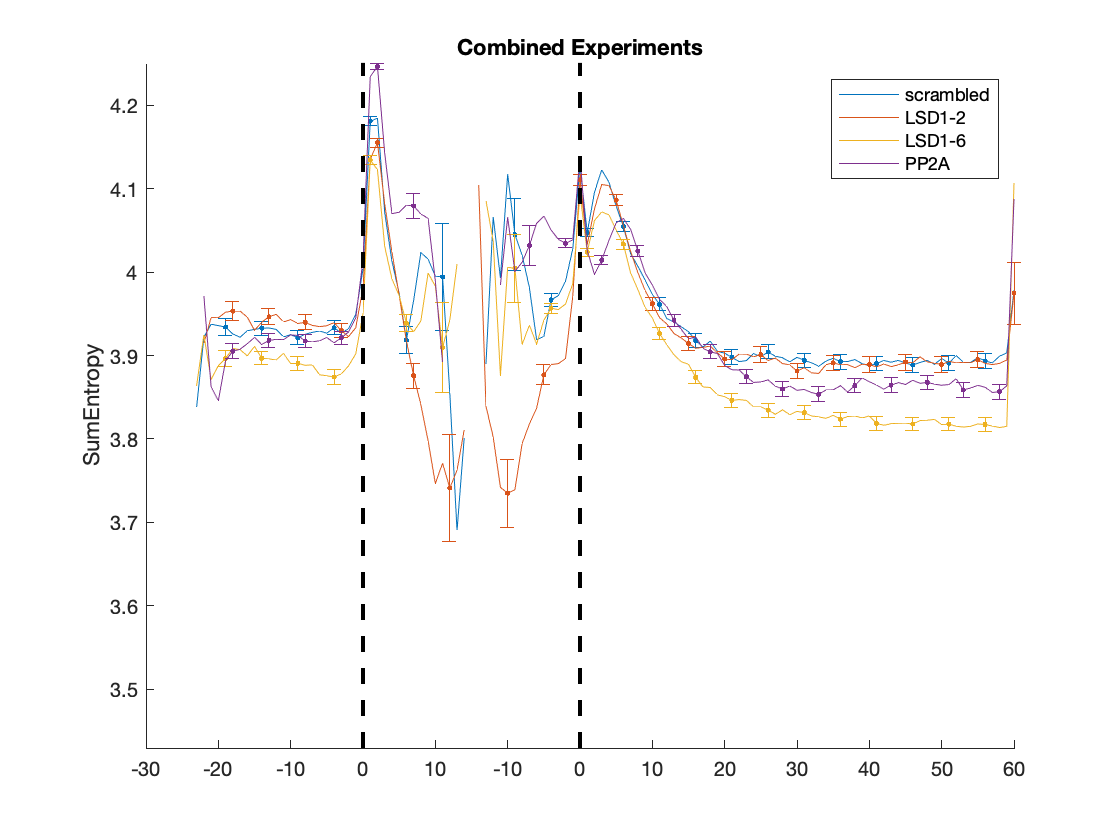

### LSD1_FusedProjects_CARSync_AdditionalFeatures_SumVariance_HeatMaps.png

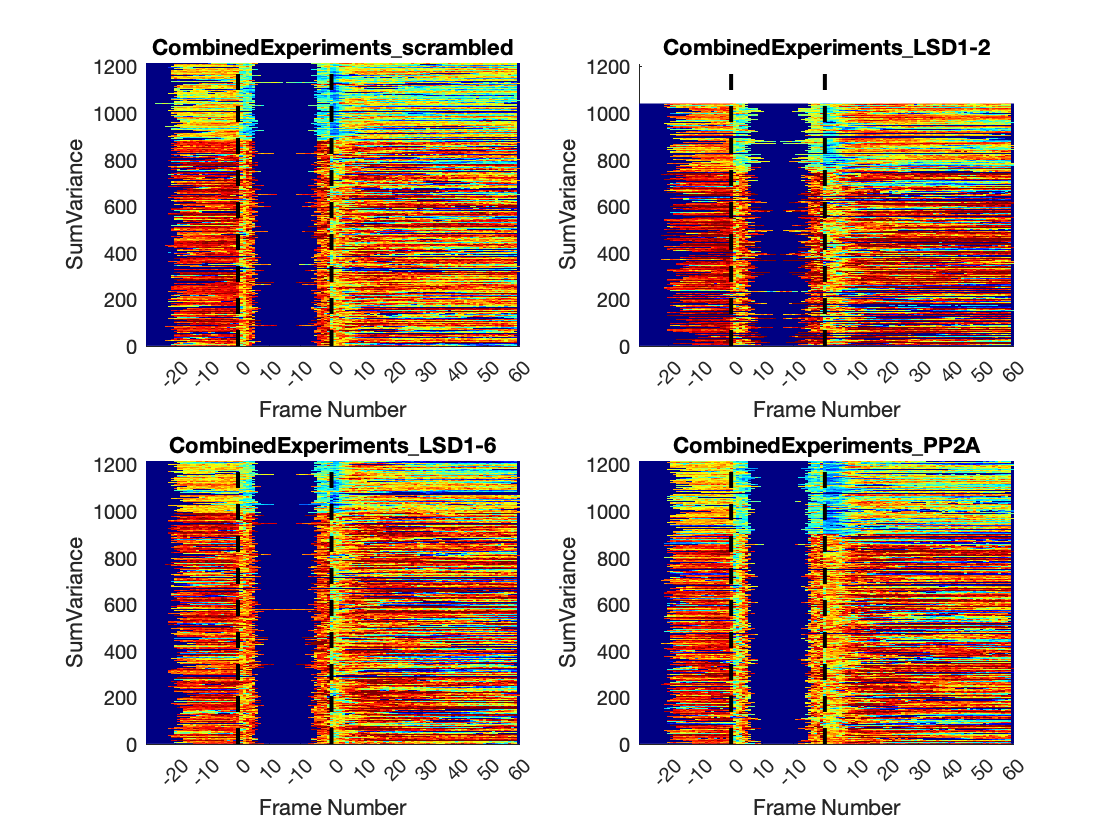

### LSD1_FusedProjects_CARSync_AdditionalFeatures_SumVariance_LinePlots.png

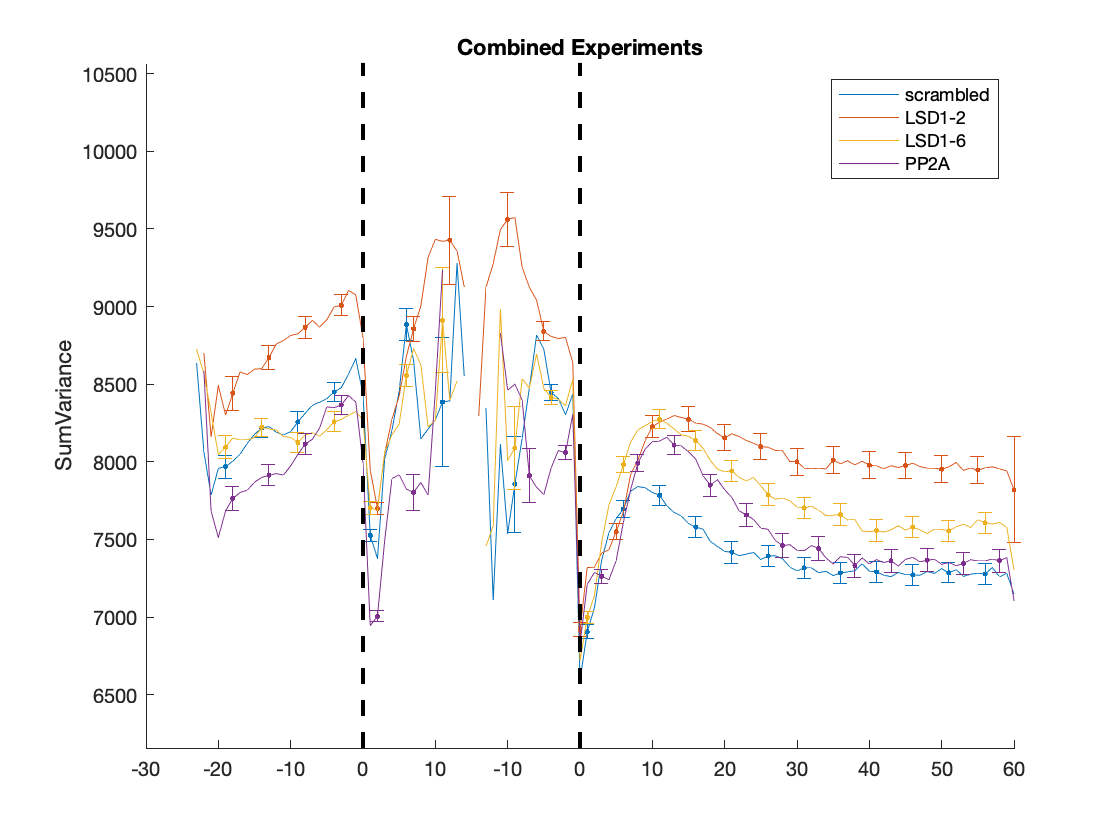

### LSD1_FusedProjects_CARSync_AdditionalFeatures_Variance_HeatMaps.png

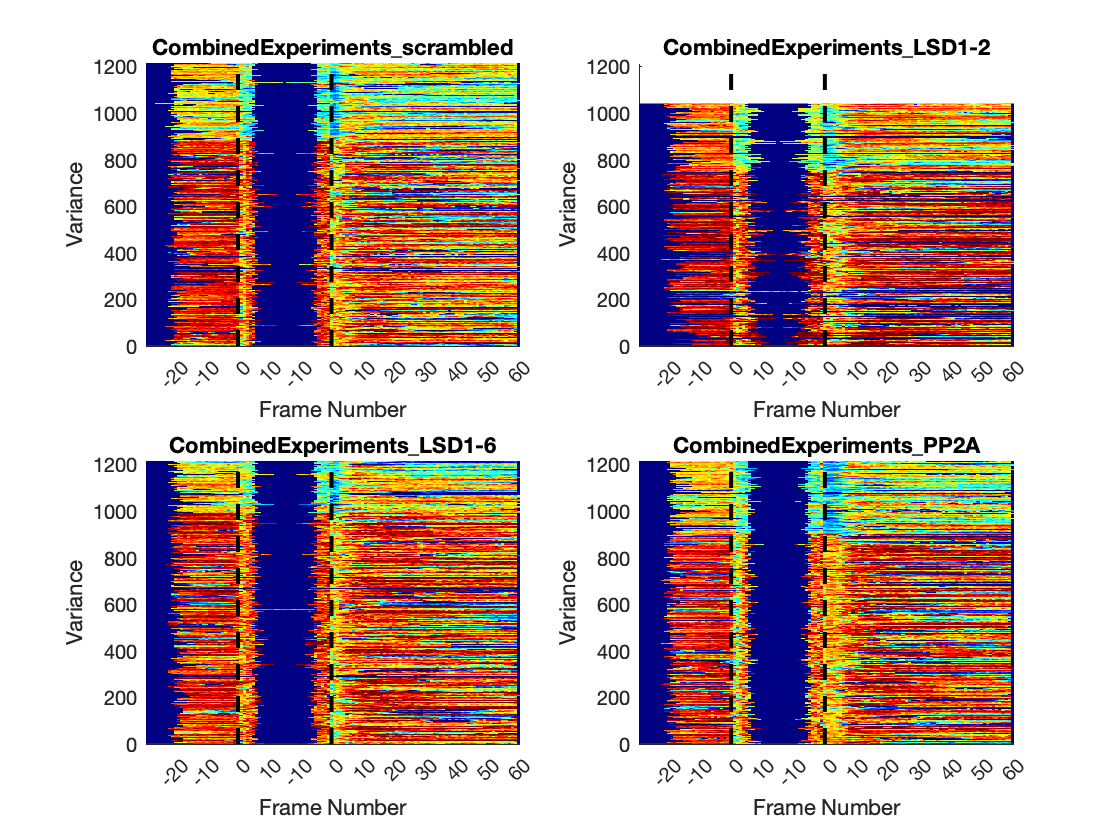

### LSD1_FusedProjects_CARSync_AdditionalFeatures_Variance_LinePlots.png

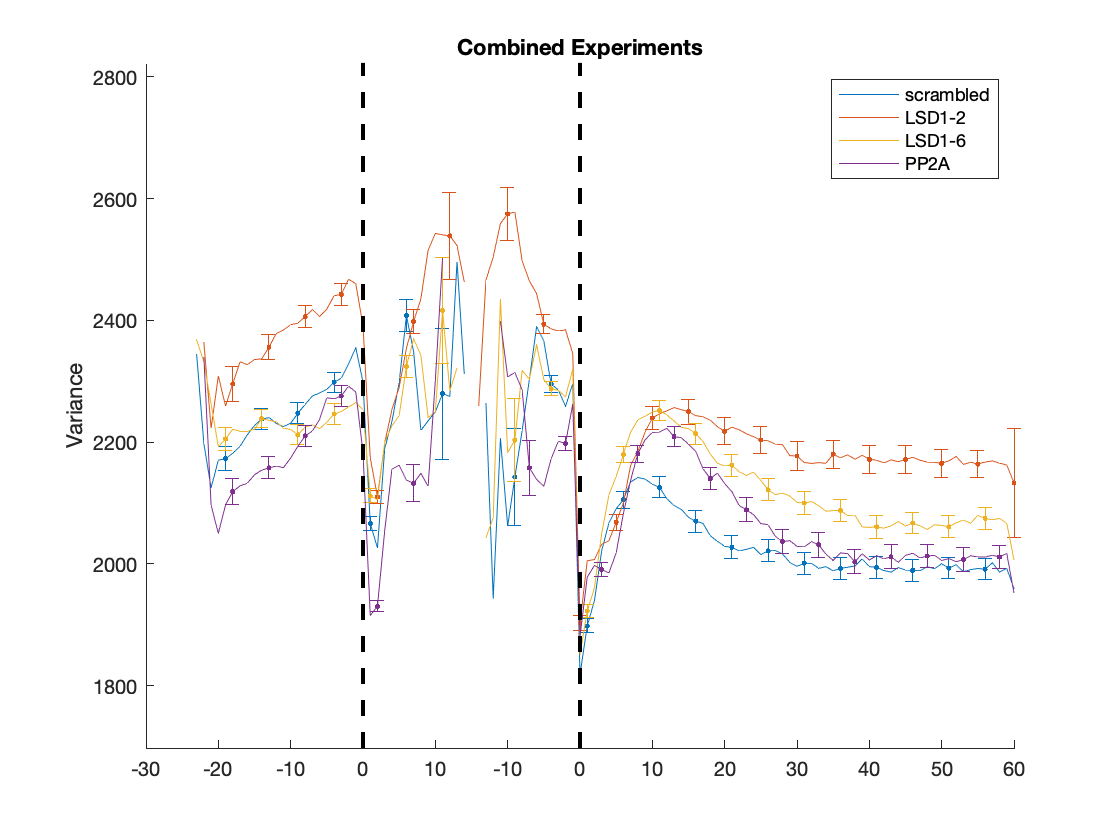

### LSD1_FusedProjects_CARSync_AdditionalFeatures_xpos_HeatMaps.png

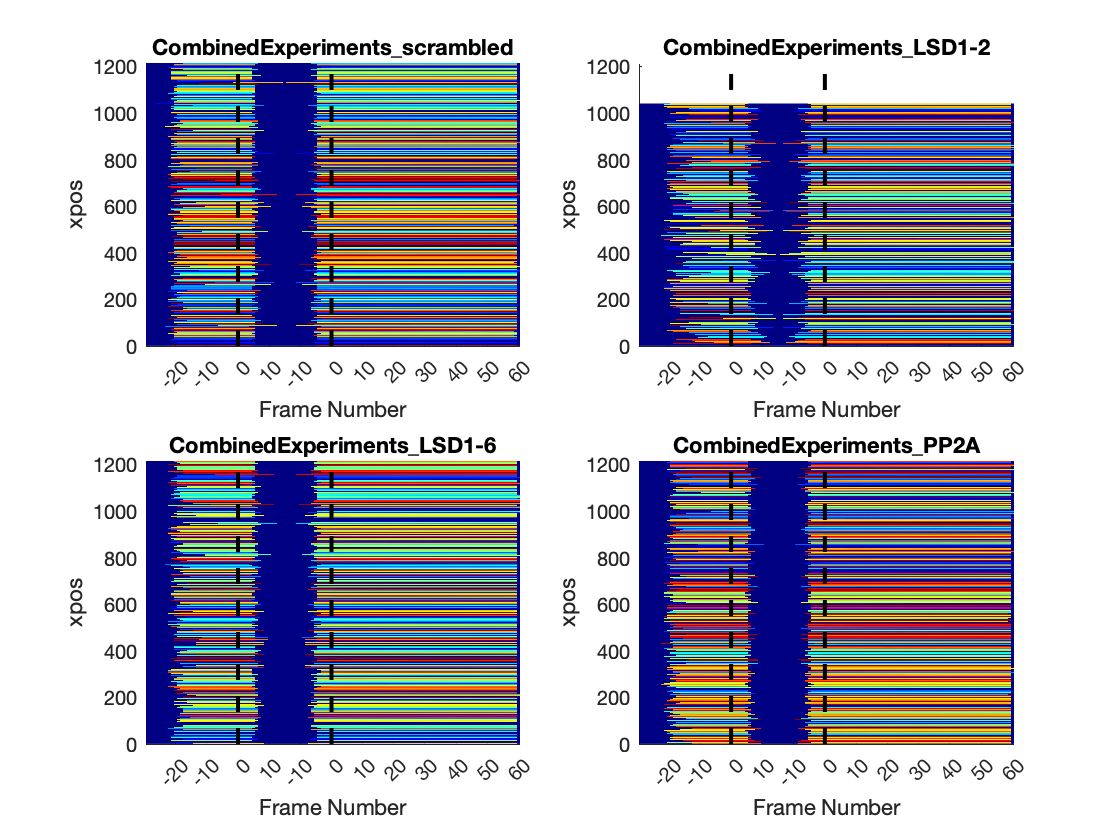

### LSD1_FusedProjects_CARSync_AdditionalFeatures_xpos_LinePlots.png

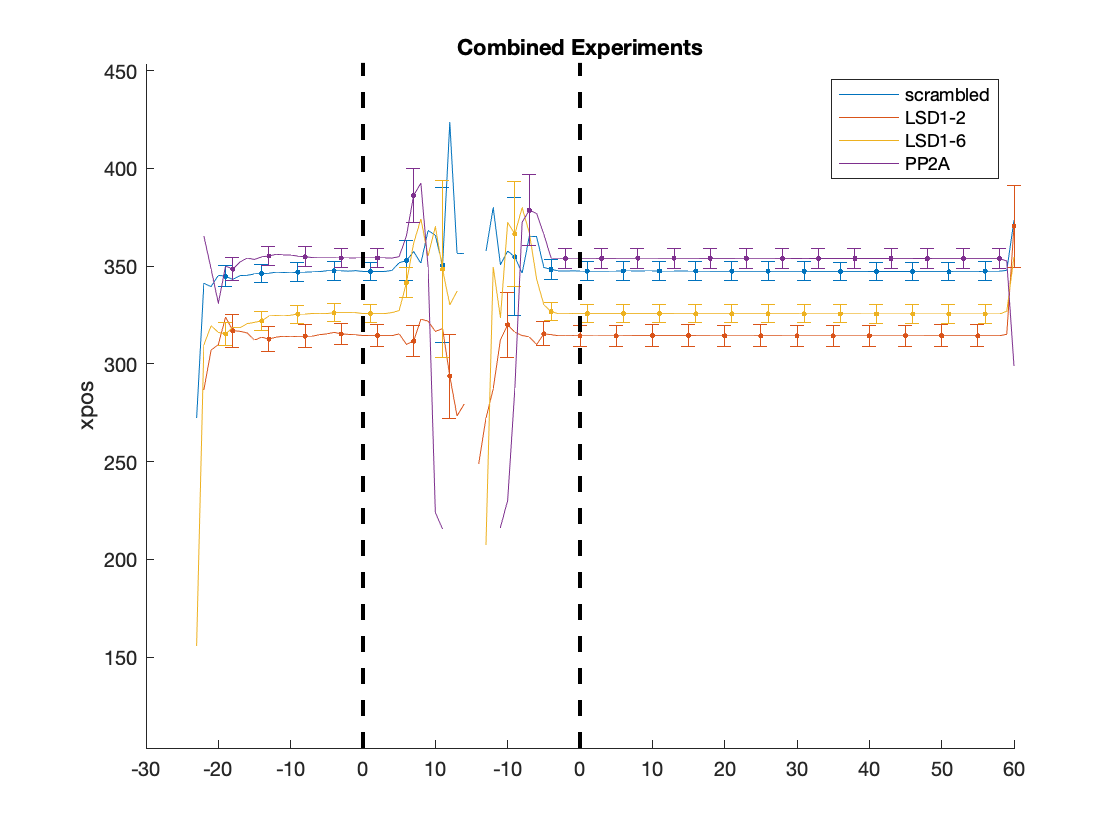

### LSD1_FusedProjects_CARSync_AdditionalFeatures_ypos_HeatMaps.png

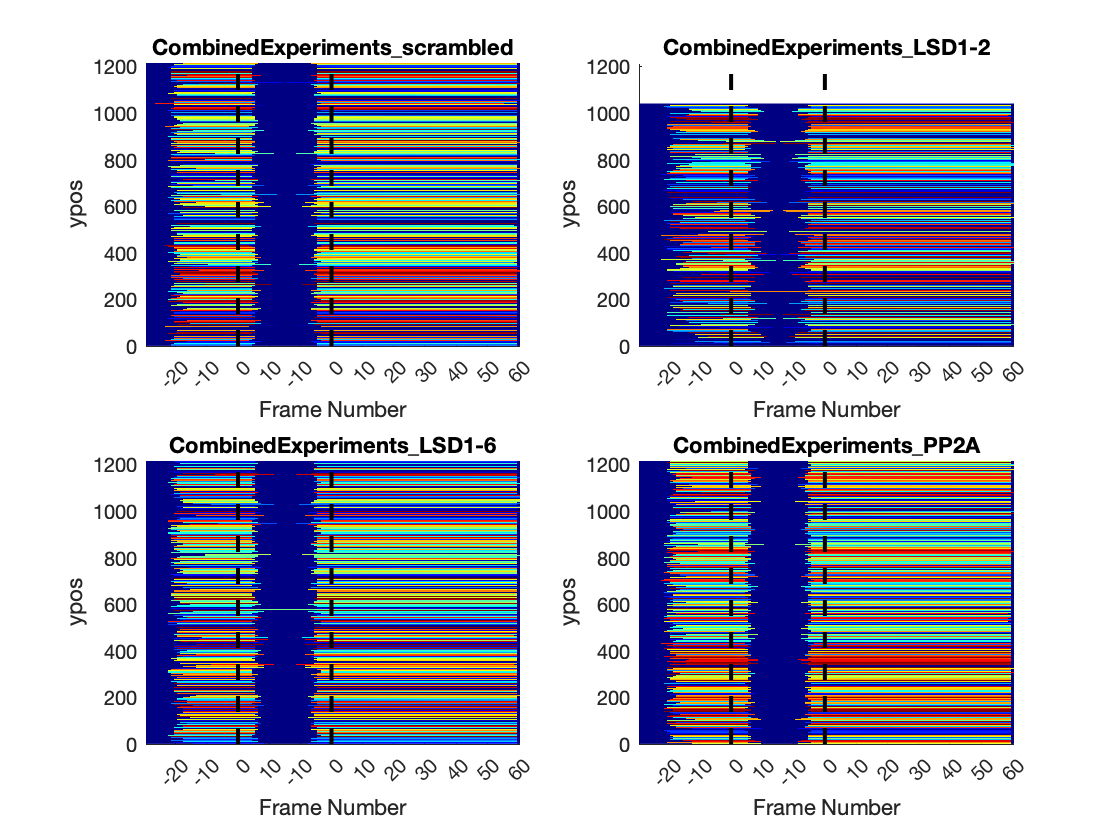

### LSD1_FusedProjects_CARSync_AdditionalFeatures_ypos_LinePlots.png

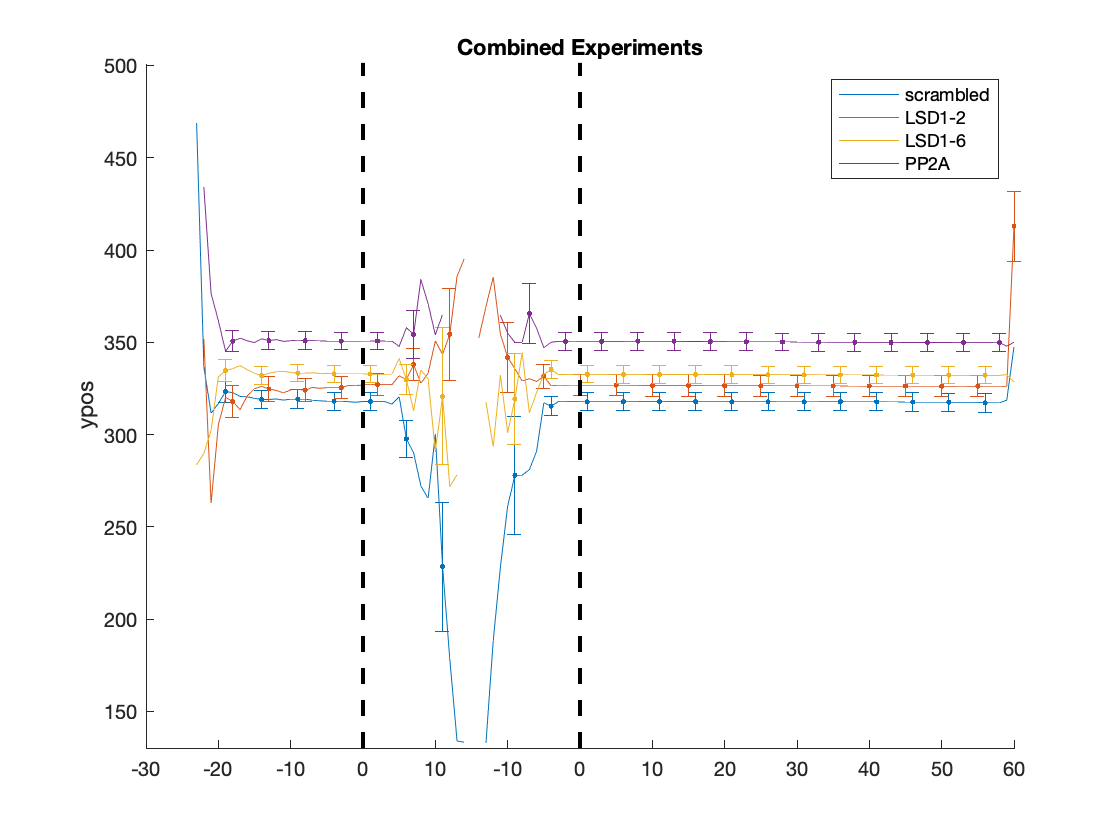

### RecQL4_FusedProjects_CARSync_AdditionalFeatures_AccumulatedOrientationDiffPMA_BoxPlots.png

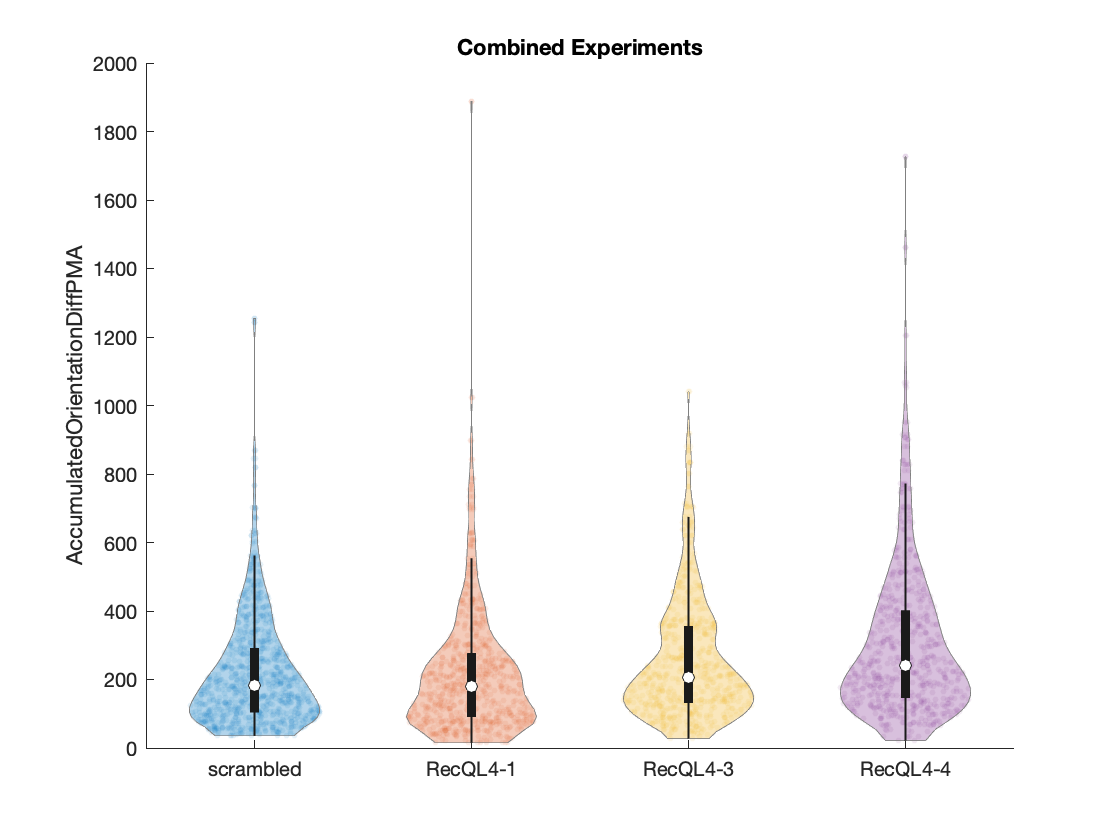

### RecQL4_FusedProjects_CARSync_AdditionalFeatures_AccumulatedOrientationDiffPMA_Histograms.png

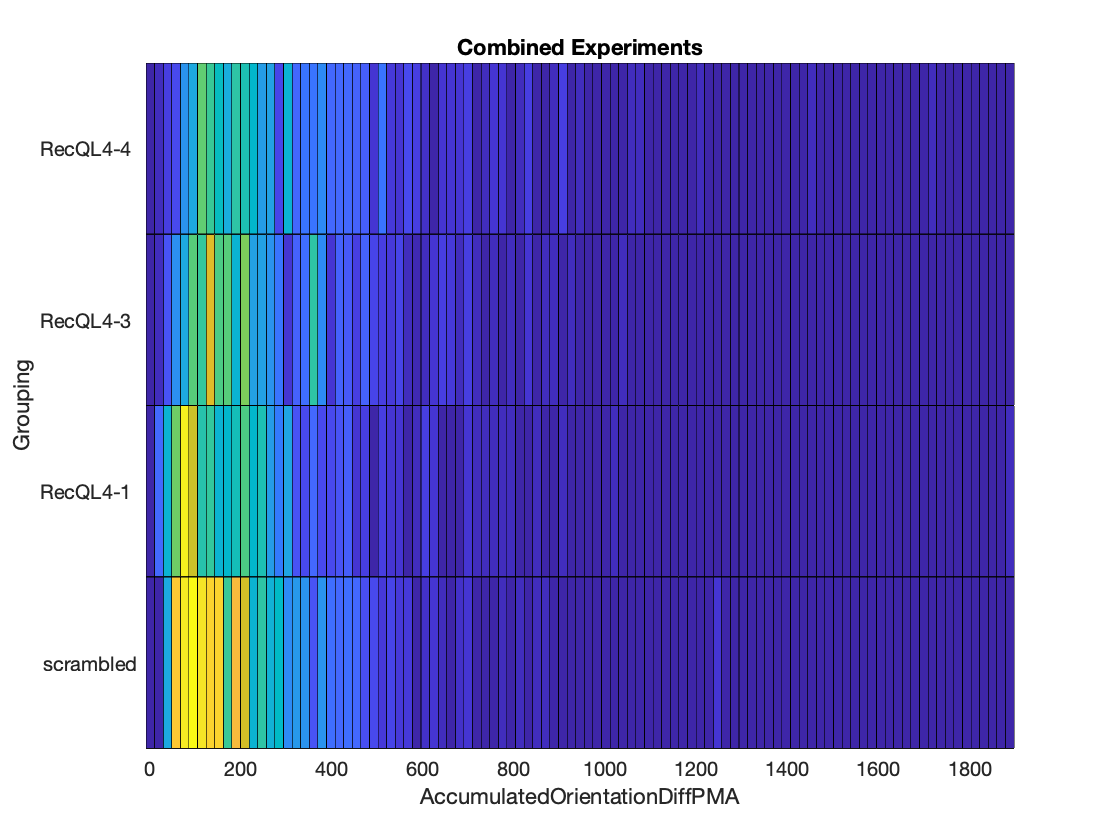

### RecQL4_FusedProjects_CARSync_AdditionalFeatures_AngularSecondMoment_HeatMaps.png

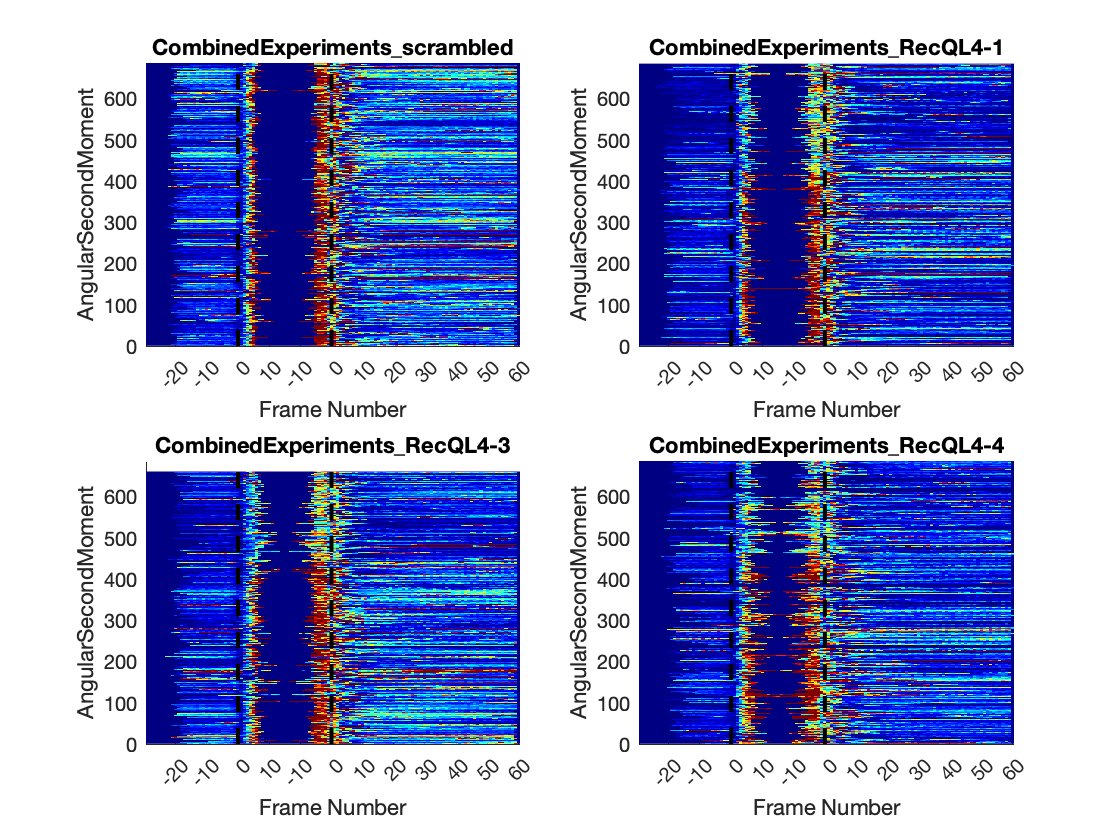

### RecQL4_FusedProjects_CARSync_AdditionalFeatures_AngularSecondMoment_LinePlots.png

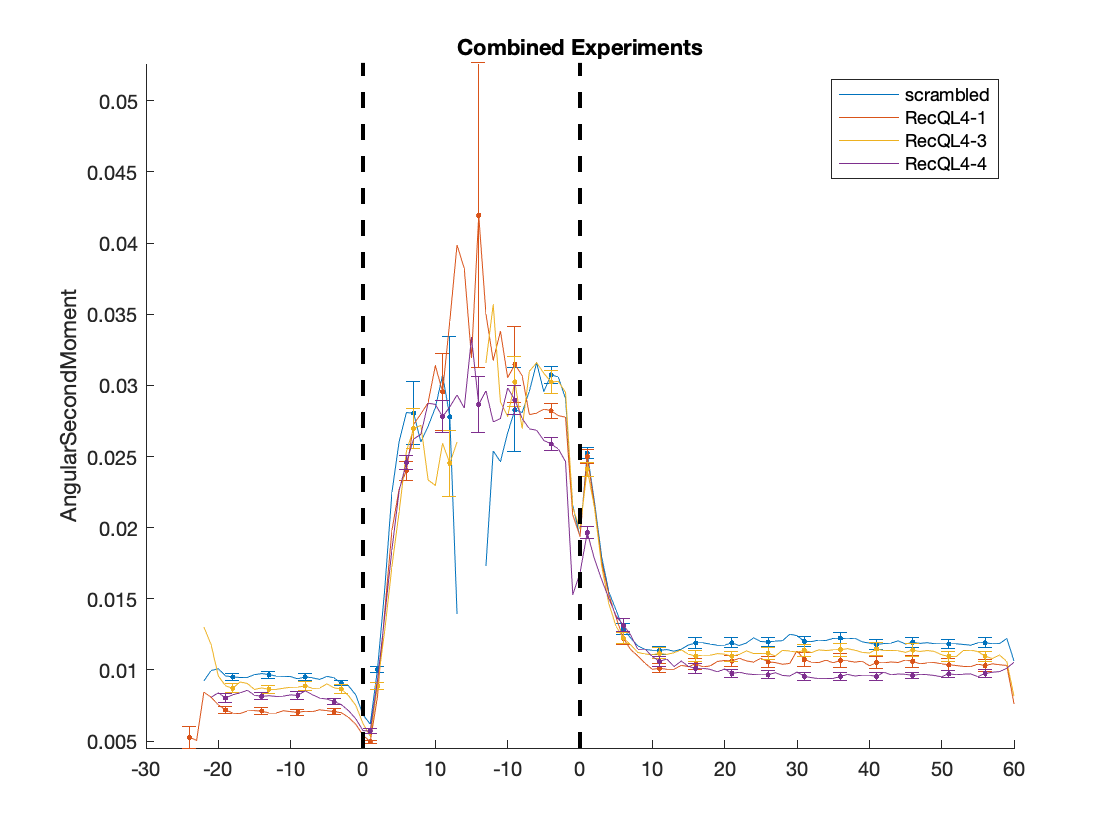

### RecQL4_FusedProjects_CARSync_AdditionalFeatures_Area_HeatMaps.png

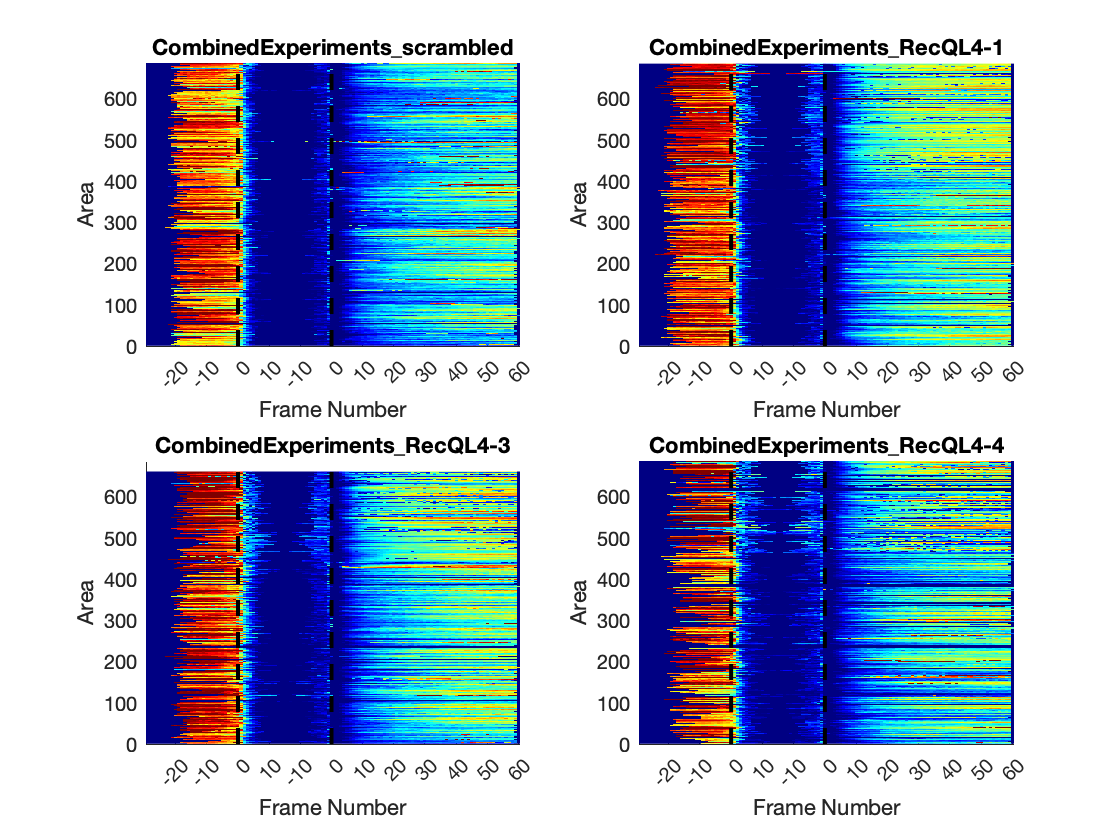

### RecQL4_FusedProjects_CARSync_AdditionalFeatures_Area_LinePlots.png

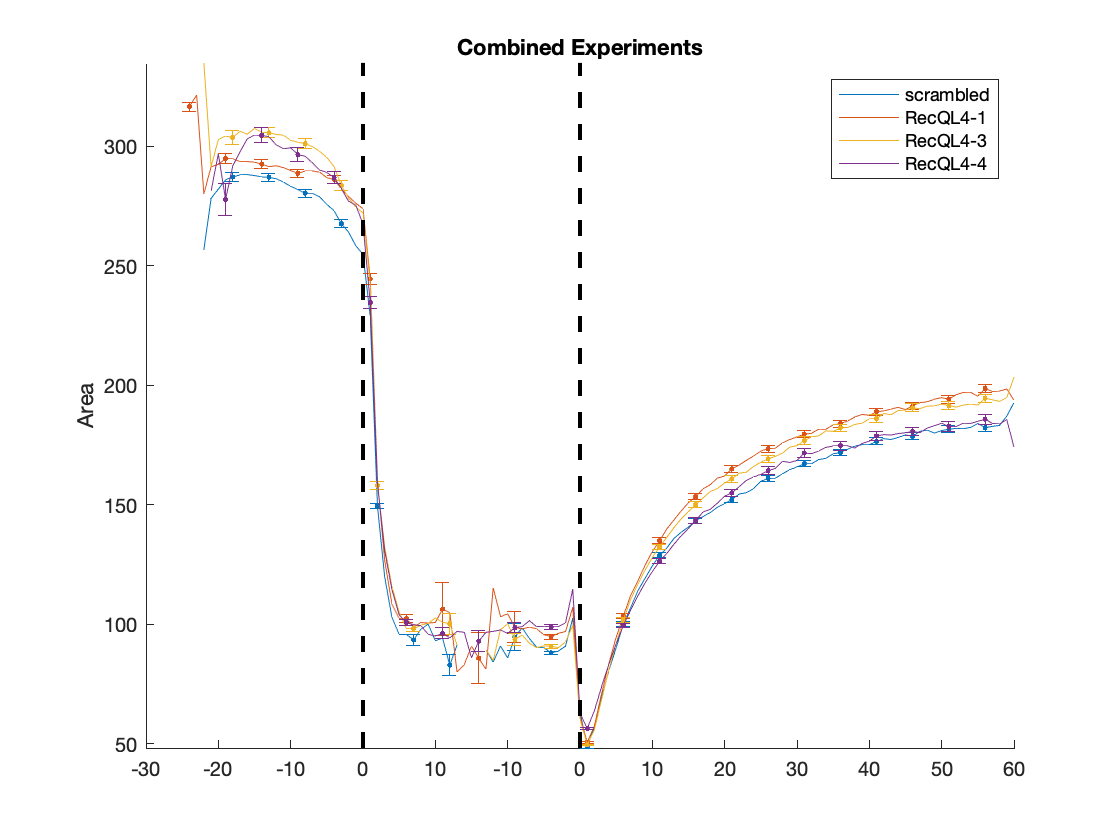

### RecQL4_FusedProjects_CARSync_AdditionalFeatures_Ch1-Ch2-MI-Ratio-Ext-disk-r=3_HeatMaps.png

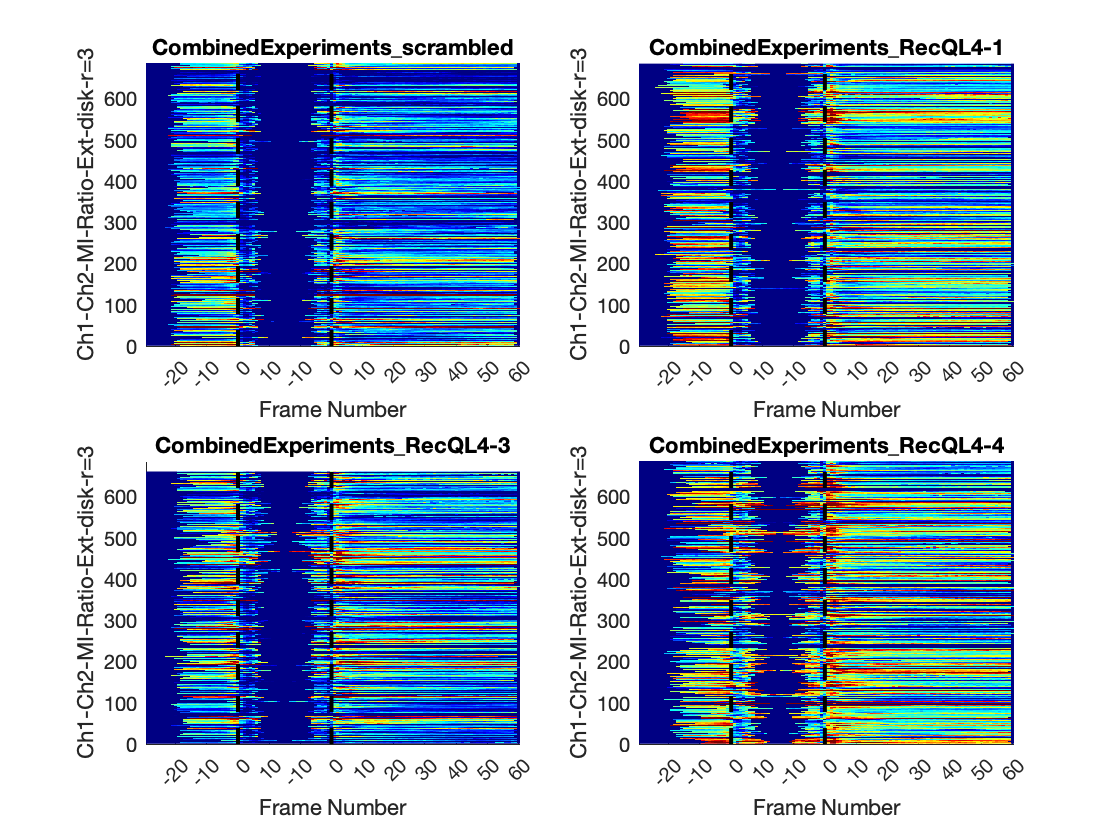

### RecQL4_FusedProjects_CARSync_AdditionalFeatures_Ch1-Ch2-MI-Ratio-Ext-disk-r=3_LinePlots.png

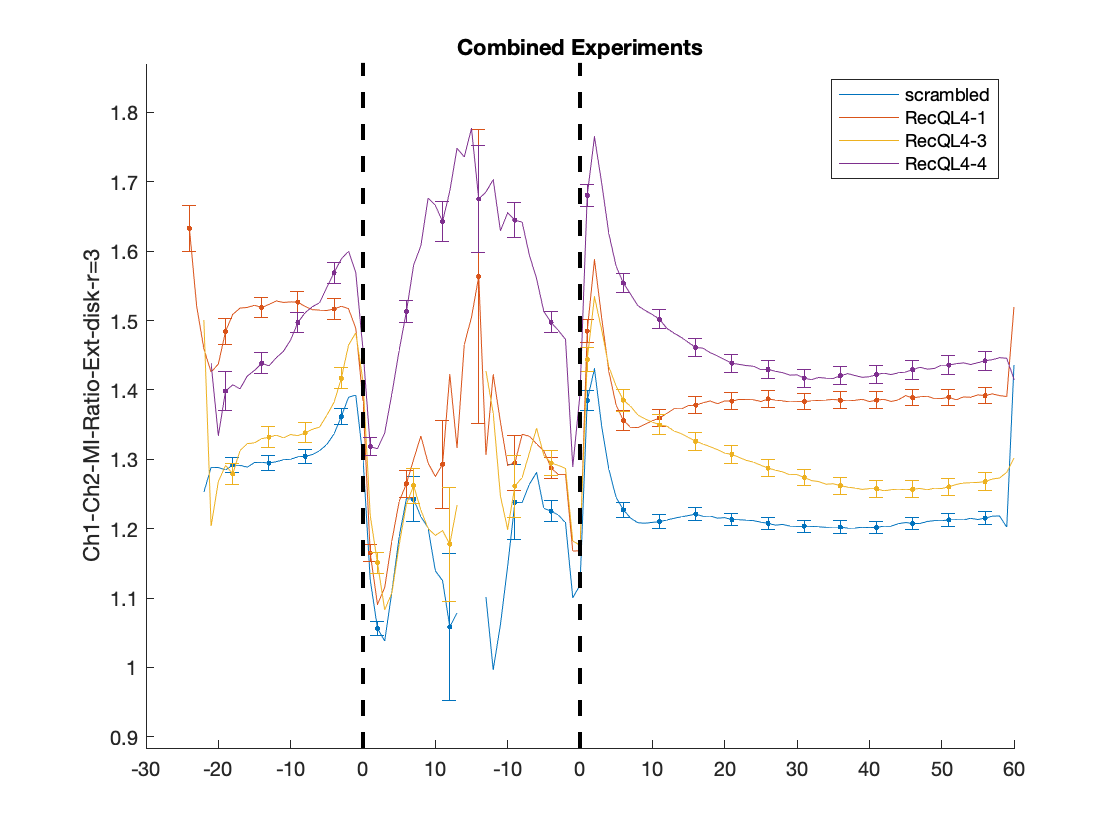

### RecQL4_FusedProjects_CARSync_AdditionalFeatures_Ch2-MaxIntensity-Ext-disk-r=3_HeatMaps.png

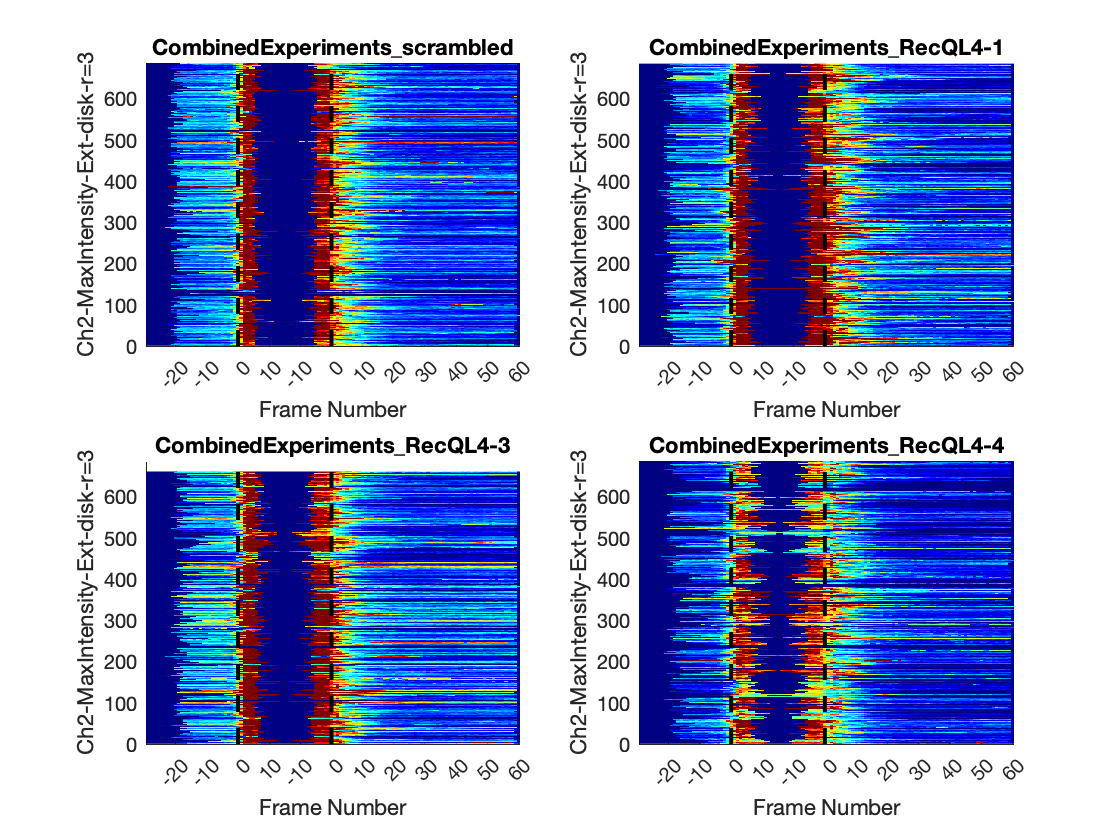

### RecQL4_FusedProjects_CARSync_AdditionalFeatures_Ch2-MaxIntensity-Ext-disk-r=3_LinePlots.png

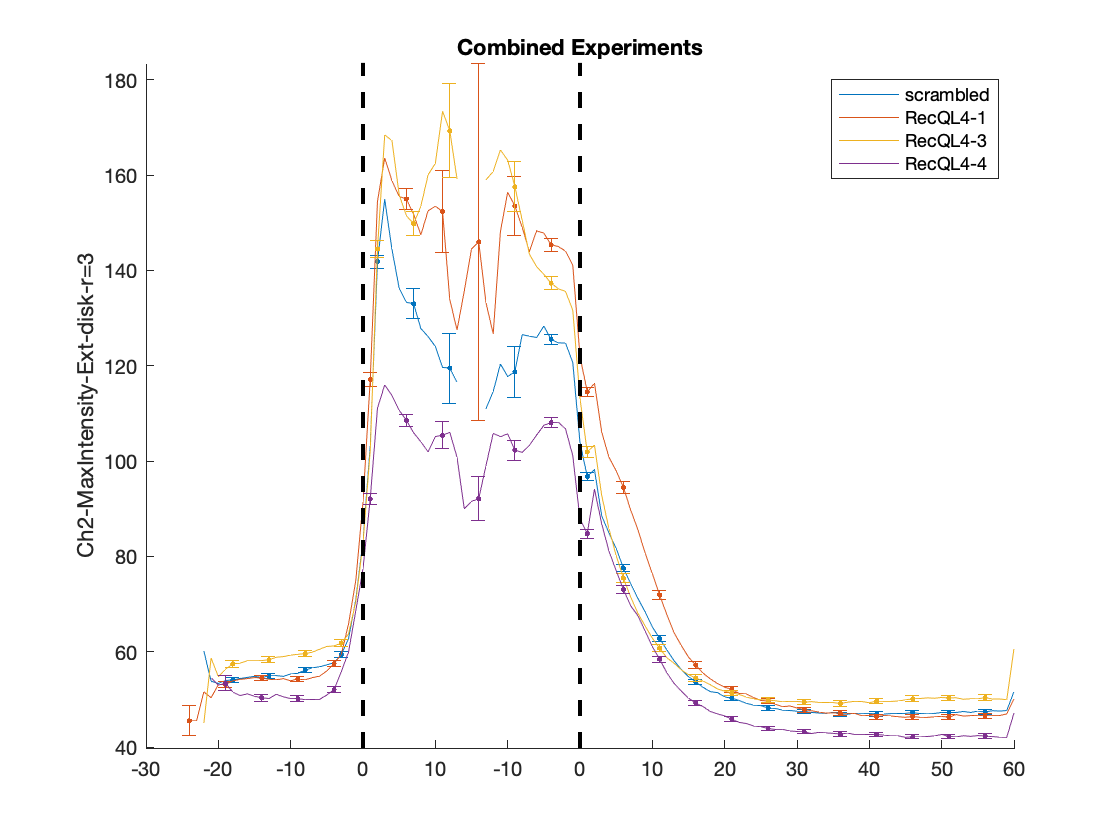
