## Supplementary material for "LiveCellMiner: A New Tool to Analyze Mitotic Progression": File S2: RecQL4_FusedProjects_CARSync_AdditionalFeatures_FeatureReport.htm

|  |  |  |  |  |  |  |  |  |  |  |  |  |  |  |  |
| --- | --- | --- | --- | --- | --- | --- | --- | --- | --- | --- | --- | --- | --- | --- | --- |
| **Time Series Name** | **Min** | **Max** | **Mean** | **Std** | **Median** | **n-Fold Inc. I->P (All)** | **n-Fold Inc. I->P (scrambled)** | **n-Fold Inc. I->P (RecQL4-1)** | **n-Fold Inc. I->P (RecQL4-3)** | **n-Fold Inc. I->P (RecQL4-4)** | **n-Fold Inc. I->A (All)** | **n-Fold Inc. I->A (scrambled)** | **n-Fold Inc. I->A (RecQL4-1)** | **n-Fold Inc. I->A (RecQL4-3)** | **n-Fold Inc. I->A (RecQL4-4)** |
| StdIntensity | 4.87 | 46.93 | 11.95 | 10.80 | 7.13 | 172.51 | 184.37 | 154.39 | 187.67 | 159.45 | 512.54 | 556.59 | 493.68 | 534.98 | 446.89 |
| StdIntensity-Normalized | 0.13 | 1.28 | 0.33 | 0.29 | 0.19 | 172.51 | 184.37 | 154.39 | 187.67 | 159.45 | 512.54 | 556.59 | 493.68 | 534.98 | 446.89 |
| StdIntensityGradMag | 10.33 | 70.98 | 21.85 | 15.73 | 14.70 | 129.98 | 137.28 | 126.62 | 135.79 | 117.20 | 316.46 | 338.18 | 321.29 | 328.56 | 267.92 |
| DifferenceVariance | 11.67 | 70.06 | 26.31 | 11.49 | 23.30 | 111.36 | 113.29 | 116.52 | 127.17 | 86.91 | 276.02 | 304.32 | 266.22 | 321.34 | 199.79 |
| Contrast | 11.75 | 70.26 | 26.43 | 11.51 | 23.41 | 110.86 | 112.75 | 116.04 | 126.57 | 86.58 | 274.40 | 302.43 | 264.86 | 319.20 | 198.83 |
| AngularSecondMoment | 0.00 | 0.05 | 0.01 | 0.01 | 0.01 | 7.47 | 3.28 | 8.56 | 3.69 | 16.15 | 223.25 | 194.99 | 282.70 | 231.30 | 191.78 |
| manualSynchronization | 1.00 | 3.00 | 2.47 | 0.80 | 3.00 | 100.00 | 100.00 | 100.00 | 100.00 | 100.00 | 200.00 | 200.00 | 200.00 | 200.00 | 200.00 |
| Ch2-StdIntensity-Ext-disk-r=3 | 3.88 | 34.84 | 9.22 | 7.53 | 6.03 | 195.05 | 193.35 | 239.84 | 166.41 | 178.84 | 183.60 | 166.21 | 238.21 | 160.37 | 173.74 |
| MeanIntensity-Normalized | 0.36 | 1.28 | 0.54 | 0.25 | 0.43 | 42.13 | 45.31 | 38.24 | 45.59 | 38.22 | 139.66 | 142.11 | 143.06 | 149.68 | 122.30 |
| MeanIntensity-NormalizedIntMean | 0.80 | 2.85 | 1.19 | 0.56 | 0.96 | 42.13 | 45.31 | 38.24 | 45.59 | 38.22 | 139.66 | 142.11 | 143.06 | 149.68 | 122.30 |
| MeanIntensity | 35.53 | 127.11 | 53.43 | 25.00 | 42.81 | 42.13 | 45.31 | 38.24 | 45.59 | 38.22 | 139.66 | 142.11 | 143.06 | 149.68 | 122.30 |
| Orientation | -68.89 | 69.66 | 0.95 | 36.73 | 0.98 | 12.99 | 64.32 | 16.15 | -104.06 | 55.99 | 134.24 | 129.24 | 32.09 | 262.31 | 119.74 |
| Ch2-MeanIntensity-Ext-disk-r=3 | 24.23 | 78.41 | 35.11 | 15.45 | 28.43 | 66.16 | 68.81 | 79.47 | 61.02 | 53.38 | 112.25 | 113.20 | 131.56 | 116.52 | 85.87 |
| InformationMeasureofCorrelationI | -0.48 | -0.19 | -0.27 | 0.07 | -0.25 | -20.82 | -25.33 | -14.50 | -20.13 | -21.85 | -102.14 | -114.52 | -93.79 | -99.41 | -96.16 |
| Area | 48.43 | 313.53 | 176.95 | 70.23 | 172.34 | -30.29 | -31.27 | -28.89 | -31.47 | -29.17 | -79.36 | -79.10 | -79.97 | -80.81 | -77.61 |
| Ch2-MaxIntensity-Ext-disk-r=3 | 39.58 | 166.34 | 66.39 | 31.40 | 52.93 | 106.33 | 109.03 | 131.33 | 98.11 | 84.13 | 77.26 | 72.83 | 103.43 | 74.19 | 58.75 |
| MinorAxisLength | 5.56 | 16.62 | 11.49 | 2.85 | 11.49 | -14.78 | -15.07 | -14.24 | -15.98 | -13.72 | -59.51 | -58.75 | -60.50 | -60.70 | -58.34 |
| MajorAxisLength | 10.82 | 26.50 | 19.62 | 3.89 | 19.76 | -14.76 | -15.63 | -14.59 | -13.95 | -14.52 | -50.64 | -51.17 | -50.22 | -52.79 | -48.13 |
| SumVariance | 3748.07 | 10421.46 | 7460.07 | 1634.92 | 7521.00 | -0.70 | 8.71 | -5.87 | 3.25 | -12.66 | 28.54 | 44.61 | 16.95 | 42.16 | 4.08 |
| Ratio-MajorAxisLength-vs-MinorAxisLength | 1.16 | 2.61 | 1.77 | 0.33 | 1.76 | 0.95 | 0.13 | 0.88 | 3.13 | -0.04 | 26.12 | 23.28 | 30.06 | 24.09 | 28.03 |
| Variance | 1052.52 | 2803.53 | 2036.17 | 430.65 | 2054.06 | -0.57 | 8.08 | -5.33 | 3.21 | -11.68 | 25.04 | 38.94 | 15.01 | 37.55 | 3.15 |
| DifferenceEntropy | 1.78 | 2.70 | 2.23 | 0.21 | 2.21 | 17.78 | 19.03 | 17.48 | 19.27 | 14.79 | 22.82 | 25.40 | 21.65 | 24.61 | 18.56 |
| InverseDifferenceMoment(Homogeneity) | 0.20 | 0.40 | 0.29 | 0.04 | 0.29 | -16.82 | -15.63 | -18.15 | -19.61 | -14.23 | -18.45 | -18.06 | -18.08 | -20.46 | -17.33 |
| Ratio-MinorAxisLength-vs-MajorAxisLength | 0.39 | 0.87 | 0.59 | 0.11 | 0.58 | 1.95 | 2.98 | 1.58 | -0.18 | 3.08 | -17.35 | -15.14 | -20.01 | -16.06 | -18.97 |
| InformationMeasureofCorrelationII | 0.87 | 1.00 | 0.93 | 0.03 | 0.93 | 4.48 | 5.67 | 3.03 | 4.36 | 4.46 | 8.35 | 10.23 | 6.66 | 7.93 | 7.93 |
| SumEntropy | 3.34 | 4.41 | 3.88 | 0.24 | 3.87 | 9.85 | 11.42 | 8.25 | 10.53 | 8.60 | 6.47 | 8.72 | 4.20 | 6.42 | 5.74 |
| Ch1-Ch2-MI-Ratio-Ext-disk-r=3 | 0.99 | 1.71 | 1.34 | 0.15 | 1.33 | -17.38 | -16.84 | -25.24 | -13.65 | -13.58 | -5.21 | -5.10 | -12.28 | -4.17 | 1.12 |
| MaximalCorrelationCoefficient | 0.76 | 0.93 | 0.85 | 0.04 | 0.85 | 2.76 | 4.26 | 0.64 | 2.45 | 3.19 | 4.44 | 6.40 | 2.66 | 3.39 | 4.61 |
| Correlation | 0.74 | 0.92 | 0.84 | 0.04 | 0.83 | 3.03 | 4.50 | 0.84 | 2.81 | 3.50 | 3.69 | 5.54 | 1.80 | 2.64 | 4.14 |
| Circularity | 0.62 | 0.99 | 0.85 | 0.07 | 0.86 | -14.80 | -13.47 | -14.85 | -16.29 | -15.11 | 2.76 | 2.64 | 1.08 | 5.02 | 2.45 |
| Entropy | 6.54 | 9.08 | 7.64 | 0.53 | 7.59 | 13.57 | 15.80 | 10.97 | 14.73 | 11.99 | 1.62 | 4.55 | -2.04 | 1.80 | 1.19 |
| SumAverage | 57.59 | 104.16 | 86.00 | 11.35 | 86.97 | -4.79 | -1.42 | -6.61 | -3.44 | -9.05 | 0.88 | 4.34 | -1.36 | 5.02 | -5.91 |
| **Single Feature Name** | **Min** | **Max** | **Mean** | **Std** | **Median** |  |  |  |  |  |  |  |  |  |  |
| IPToMALength\_Frames | 5.00 | 29.00 | 12.35 | 4.29 | 11.00 | - | - | - | - | - | - | - | - | - | - |
| IPToMALength\_Minutes | 15.00 | 87.00 | 37.05 | 12.87 | 33.00 | - | - | - | - | - | - | - | - | - | - |
| InterphaseMeanIntensity | 18.65 | 138.80 | 45.26 | 13.52 | 43.72 | - | - | - | - | - | - | - | - | - | - |
| AccumulatedOrientationDiffPMA | 15.04 | 1888.32 | 248.59 | 193.05 | 198.83 | - | - | - | - | - | - | - | - | - | - |
| Ch2-MeanIntensity-Ext-disk-r=3-intMean | 15.04 | 106.82 | 29.28 | 6.97 | 28.45 | - | - | - | - | - | - | - | - | - | - |
| Ch2-MeanIntensity-Ext-disk-r=3-pmaMean | 15.37 | 177.31 | 68.63 | 17.36 | 68.59 | - | - | - | - | - | - | - | - | - | - |
| Ch2-MeanIntensity-Ext-disk-r=3-atiMean | 14.70 | 87.68 | 30.03 | 5.89 | 29.56 | - | - | - | - | - | - | - | - | - | - |
